# Mechanistic dissection of global proteomic changes in rats with heart failure and preserved ejection fraction

**DOI:** 10.1101/2021.09.06.456608

**Authors:** Daniel Soetkamp, Aleksandra Binek, Romain Gallet, Geoffrey de Couto, Peter Kilfoil, Rowann Mostafa, Vidya Venkatraman, Ronald Holewinski, Amy D. Bradshaw, Joshua Goldhaber, Michael R. Zile, Eduardo Marbán, Jennifer E. Van Eyk

**Affiliations:** Smidt Heart Institute, Cedars-Sinai Medical Center, Los Angeles, California, USA; Advanced Clinical Biosystems Center, Cedars-Sinai Medical Center, Los Angeles, California, USA; Medical University of South Carolina and Ralph H. Johnson Veterans Affairs Medical Center, Charleston, South Carolina

**Keywords:** Heart failure and preserved ejection fraction (HFpEF), Diastolic dysfunction, Cardiosphere-Derived Cell (CDC)

## Abstract

**Introduction:** Heart failure with preserved ejection fraction (HFpEF) is characterized by diastolic dysfunction, pulmonary congestion and exercise intolerance. Previous preclinical studies show that treatment with cardiosphere-derived cells (CDCs) improves diastolic function, attenuates arrhythmias and prolongs survival in a rat HFpEF model. Here we characterize the myocardial proteome and diastolic function in HFpEF, with and without CDC therapy. As an initial strategy for identifying pathways worthy of further mechanistic dissection, we correlated CDC-responsive proteomic changes with functional improvements.

**Methods and Results:** Dahl salt-sensitive rats fed high-salt diet, with verified diastolic dysfunction, were randomly assigned to intracoronary CDCs or placebo. Dahl rats fed a low salt diet served as controls. Phenotyping was by echocardiography (E/A ratio) and invasive hemodynamic monitoring (time constant of relaxation Tau, and left ventricular end-diastolic pressure [LVEDP]). CDC treatment improved diastolic function as indicated by a normalized E/A ratio, a 33.3% reduction in Tau, and a 47% reduction of LVEDP. Mass spectrometry of left ventricular tissues (n=6/group) revealed changes in transcription and translation pathways in this rat HFpEF model and was also recapitulated in human HFpEF. These pathways were enhanced following CDC treatment in the animal model (205 proteins and 32 phosphorylated residues accounting for 37% and 19% of all changes, respectively). Among all CDC-sensitive pathways, 65% can be linked to at least 1 of 7 upstream regulators, among which several are of potential relevance for regulating protein expression. To probe newly-synthesized proteins AHA labeling was carried out in isolated rat cardiomyocytes obtained from HFpEF groups, with and without CDC therapy. Five of the initial upstream regulators (HNF4A, MTOR, MYC, TGFβ1, and TP53) were linked to proteins expressed exclusively (or increased) with CDC treatment. All 32 phosphorylated residues of proteins involved in transcription/translation altered specifically by CDC treatment had predicted kinases (Protein kinase C (PKC) being the most dominant) and known to be regulated by MYC, TGFβ1 and/or TP53. Western blot analysis of those 5 upstream regulators showed that TGFβ1, TP53, and Myc were significantly decreased in LV from CDC treated animals, whereas MTOR and HNF4A showed a significant increase compared to HFpEF alone. The cellular quantities of several upstream regulator correlated with indices of diastolic function (E/A ratio, Tau and/or LVEDP). Since CDCs act via the secretion of exosomes laden with signaling cargo, it is relevant that all 7 upstream regulators could, in principle, be regulated by proteins or miRNA that are present in CDC-derived exosomes.

**Conclusion:** We identified key cellular regulators of transcription and translation that underlie the therapeutic effects of CDCs in HFpEF, whose levels correlate quantitatively with measures of diastolic function. Among the multifarious proteomic changes associated with rat model of HFpEF which were also observed in human HFpEF samples, we propose that these regulators, and downstream effector kinases, be prioritized for further dedicated mechanistic dissection.

## Introduction

Heart failure with preserved ejection fraction (HFpEF) is characterized by diastolic dysfunction, pulmonary congestion, exercise intolerance and high mortality [1, 2]. Hypertrophy, fibrosis [3], abnormal Ca^2+^-handling [4, 5] and/or titin phosphorylation and isoform expression have all been posited as contributors [6-9]. Nevertheless, the mechanistic underpinnings of HFpEF remain poorly understood. Therapeutic trials in HFpEF patients have generally failed [10, 11] and there is no effective treatment. An alternative approach, using cardiosphere-derived cell (CDC) therapy, is now being tested clinically. The Regress-HFpEF trial (NCT02941705, clinicaltrials.gov), was motivated by signs of disease-modifying bioactivity in previous studies of ischemic, dilated and heritable cardiomyopathy [12-19], plus promising results in a Dahl salt sensitive rat model of HFpEF. There, intracoronary CDCs normalized diastolic function, reduced fibrosis and inflammation, and attenuated mortality despite persistent hypertension and hypertrophy [20].

Given their striking functional and histological benefits, CDCs may be useful in the mechanistic dissection of HFpEF. We reasoned that HFpEF would likely exhibit marked, global changes in protein expression and post-translational modifications, making it virtually impossible to identify causal changes; one way to narrow the search would be to focus on CDC-sensitive changes in which HFpEF abnormalities revert towards control levels. Thus, we sought to identify the cardiac proteome and phospho-pathways and key tissue regulators with HFpEF compared to control animals and then narrow down based on those that are targeted additionally by CDC therapy based on overlap of dyregulated pathway uncovered from analysis of human HFpEF myocaridum. Given that CDCs work by secreting exosomes rich in signaling cargo, we further related the observed CDC-induced proteomic changes to specific micro RNAs and proteins within CDC exosomes (CDCexo). Using mass spectrometry (MS)-friendly labeling techniques to capture newly synthesized proteins in isolated HFpEF myocytes upon CDCexo treatment or kinase modulators combined with proteomic approaches several CDC-responsive key regulators of the transcriptional and translational machinery and its phosphoregulation were pinpointed as mechanistic candidates worthy of prioritized further study.

## Methods

### Cell culture

H9C2 cells were cultured in DMEM high-glucose medium and supplemented with fetal calf serum (10%), 2 mM of L-glutamine, 100 U/mL of penicillin and 100 g/mL of streptomycin at 37 °C in 5% CO_2_. Neonatal rat ventricular myocytes (NRVMs) were isolated from P2 neonatal Sprague–Dawley rats as previously described [21]. The cells were plated on fibronectin-coated 6-well plates at a density of 1.5 million cells/well in Dulbecco’s Modified Eagle Medium (DMEM) containing 10% Fetal Bovine Serum (Gibco) media and incubated at 37 °C, with 5% CO_2_ for 24 h.

### Animals and isolated myocytes

All rats were treated according to the Guide for the Care and Use of Laboratory Animals published by the US National Institutes of Health (NIH publication no. 85–23, revised 1996) and approved by the Institutional Laboratory Animal Care and Use Committee of the NIH, Bethesda, MD, USA.

Male Dahl SS rats (Charles River) were either switched from a low salt diet (0.3% NaCl) to a high salt diet (8% NaCl) (diastolic dysfunction) or maintained on a low salt diet (control) at 7 weeks of age. After 6–7 weeks, rats with induced diastolic dysfunction and preserved systolic function were randomly assigned to allogeneic CDC treatment (injection of 5×10^5^ cells re-suspended in 100 μl PBS pH 7.4) or vehicle (PBS). CDCs were generated from explanted Wistar-Kyoto rat hearts (Charles River) hearts as described [22]. Echocardiography and/or hemodynamic measurements were performed to characterize heart phenotypes 4 weeks after treatment [20]. Adult cardiomyocytes were isolated from rat left ventricles as previously described [5].

Human samples were obtained from intraoperative left ventricular (LV) myocardial biopsies obtained from patients (n=21) recruited to undergo coronary artery bypass grafting with a preserved LV ejection fraction at Medical University of South Carolina. LV samples were flash frozen and stored at -80°C until use. Patients were grouped as control (n=9), and heart failure (HFpEF) (n=7) and have been previously assessed for myocardial stiffness [4] (see online table 8 for clinical parameters). The institutional review boards of participating institutions approved the protocol, and all patients gave written informed consent.

### Isolation of CDC exosomes

Extracellular vesicles (EVs) were enriched from CDC culture media as previously described [23]. In brief, rat CDCs were conditioned for 15 days in glucose-containing serum-free basal media to increase their potency. Cell debris was removed utilizing sterile vacuum filtration (0.45 um) and then enriched via ultrafiltration using 10 kDa MWCO Filtration unit (Vivacell). CDCexo with human origin were isolated by centrifugation for 10 min at 800 g to pellet cell debris, followed by centrifugation at 20,000 g for 30 min to pellet large vesicles and separate exosomes. Secreted proteins were removed by washing exosomes twice with PBS via ultracentrifugation with 100,000 g for 60 min.

### Sample handling and preparation for proteomics

Thawed rat or human LV tissue were homogenized, fractionated, reduced, alkylated, trypsin digested, and desalted with and without TiO_2_ enrichment prior to MS analysis. The latter enrichment allowed phospho-proteomics to be carried out [24]. Detailed methods are in the online supplements.

### Quantification of newly synthesized proteins (NSPs)

Adult cardiomyocytes harvested from HFpEF and control rats were washed twice with methionine depleted media after adhesion for 3 h. Cell were depleted of methionine for 1h and then 1 mM L-azidohomoalanine (AHA, Cambridge Isotope Laboratories) was added to the media with a dose of CDCexo (10^3^ EV/ cell) or PBS. After 5 h incubation at 37°C with 5% CO_2_ cell were washed twice with PBS containing proteases inhibitors (Thermo Fisher Scientific), spun down at 1000 g for 5 min at 4°C and snap frozen in liquid nitrogen. To assess effect of Protein kinase C (PKC) isoforms on transcription, H9C2 cells were starved in DMEM without FBS for 24 h before 50 um phenylephrine (PE) or DMSO (control) was added to induce hypertrophy. After 24 h, DMEM media was replaced with methionine depleted DMEM and fresh PE or DMSO for 1 h before addition of 1 mM AHA with either 1,2,3,4-Tetrahydro Staurosporine (PKCα inhibitor, Satan Cruz Biotechnology), PKCβ Inhibitor (CAS 257879-35-9, Santa Cruz Biotechnology), or 1 µm Rottlerin (PKC δ inhibitor, Santa Cruz Biotechnology) or buffer (control).

### Mass spectrometry and data analysis

Rat and human LV tissue, cell culture samples and CDC_exo_ were analyzed by liquid chromatography-tandem mass spectrometry (LC-MS) on a Dionex Ultimate 3000 NanoLC connected to an Orbitrap Elite (Thermo Fisher Scientific) equipped with an EasySpray ion source or on a Dionex Ultimate 3000 NanoLC connected to an Orbitrap Fusion™ Lumos™ Tribrid™ Mass Spectrometer (Thermo Fisher Scientific) equipped with an EasySpray ion source as previously reported [25]. The MS-based proteomics data will be deposited in the ProteomeXchange Consortium (http://www.proteomexchange.org) via the PRIDE partner repository [26]. Detailed methods are listed in the online supplements.

### Western blot

For Western blot (WB) analysis a total of 30 µg of frozen LV rat tissue (n=3-6) per treatment group was lysed. Concentrations of proteins of interests were quanttfied via antibody detection and HRP-conjugate chemiluminescence. Detailed methods are listed in the online supplements.

## Results

Control rats receiving the high salt diet develop the HFpEF phenotype featuring hypertension, LV hypertrophy, and diastolic dysfunction as indicated by reduced E/A ratio (23.7 ± 4.8%, p<0.05, n=7), prolonged Tau (64.9.7 ± 30.7%, p<0.05, n=7) and increased LVEDP (89.2 ± 15.6%, p<0.05, n=7). Four weeks after administration of a single bolus of CDCs, rats with HFpEF demonstrated improved LV stiffness with normalized E/A ratio (15.5 ± 3.8%, p<0.05, n=7), reduced Tau (38.5 ± 8.4%, p<0.05, n=7) and reduced LVEDP (61.4 ± 9.7%, p<0.05, n=7) compared to placebo. However, CDCs did not alter extent of hypertension or LV hypertrophy (Figure 1).

**Figure 1:**
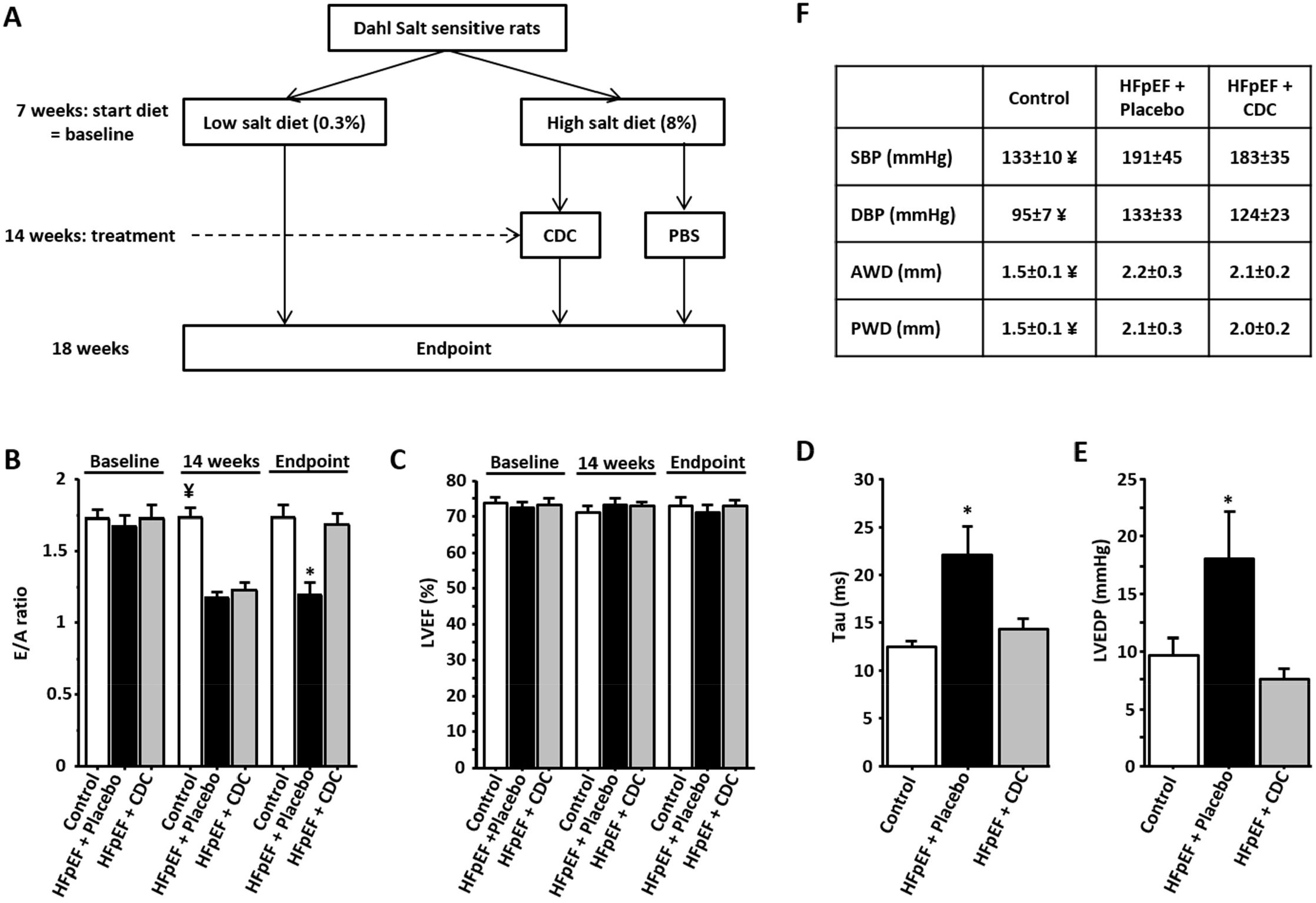
(A) Design of the study. (B) E/A ratio decreases after 7 weeks of high salt diet and normalizes 4 weeks after treatment in the CDC-treated but not in PBS-treated animals (n=3-6/group). (C) LVEF is preserved in all groups during follow-up. (D) Four weeks after treatment, Tau is increased in PBS-treated animals compared to control and CDC-treated animals. (E) LVEDP is increased in PBS-treated animals indicating heart failure but is normal in CDC-treated and control animals. (F) There are no differences in blood pressure between CDC-treated and PBS-treated animals which all have increased blood pressure compared to control. There are also no differences in cardiac hypertrophy between these 2 groups which both exhibit cardiac hypertrophy compared to control. * for p<0.05 vs. Control and CDCs. ¥ for p<0.05 vs. PBS and CDCs.

To identify underlying proteome changes, tryptic peptides obtained from LV tissue homogenates of control rats on low salt diet or high salt diet with or without CDC treatment were analyzed (n=5 per experimental group). To deal with the sparse annotation of the rat genome, we used both the rat annotated and unannotated and mouse annotated protein databases and following data search removed peptide and protein name redundancy. A total of 3058 unambiguous proteins and 3396 phosphorylated Ser, Thr and Tyr residues (n=6 per experimental group) were quantified based on the search against a combination of the three protein databases (for details on search approach see online supplement). 41% of total proteome (1262, p<0.05) proteins were different between HFpEF and control, with the largest number of differentially expressed proteins (348 proteins) being involved in regulating protein expression (supplemental table 1). Eqivalent proteomics analysis of LV obtained from human hearts, in which 3780 unambigous proteins were quantified revealed that HFpEF has a distinctive proteome compared to control donor hearts (Supplemental figure 1, Supplemental table 2). In human disease, the myocardial DEPs center on the tissue regulators Myc proto-oncogene protein (Myc), mammalian target of rapamycin (mTOR)/Rapamycin-insensitive companion of mammalian target of rapamycin (RICTOR) (Supplemental figure 1). Of interest, is the comparison between the proteome change and the pathways in human HFpEF and the animal model. In this case, 44% of the DEPs altered in human HFpEF compared to control hearts (172 of the total of 387 DEPs) were also altered in the rat HFpEF model. The overlapping DEPs, present in both species, involved cellular metabolism, fibrosis, inflammation, cell death, angiogenesis, and myofilament contraction (Supplemental figure 1, Supplemental table 2) although there were additional processes such as protein transport and protein synthesis that were changed differentially in human HFpEF compared to the rat model. Perhaps more important, there is a mechanistic convergence and agreement with respect to the primarly upstream regulators; Myc, Cellular tumor antigen p53 (TP53), HNF4A, and RICTOR with HFpEF regardless of species (supplemental figure 1C).

Focusing on CDC treatment response in the HFpEF animals, the 573 proteins were quantitatively altered following therapy compared to placebo treatment. Of those, 205 DEPs are involved in regulating protein expression (Supplemental Table 1). Specifically, transcriptional regulation, translational regulation, mRNA-processing, DNA organization and nuclear import/export (93, 62,42,19 and 6 DEPs, respectively) were the most represented GO functions. Upstream regulator analysis revealed 9 dominant proteins that collectively accounted for the potential regulation of the majority of the CDC-senstive DEPs and those specifically involved in translation and transcription (65% and 45%, respectively) (Table 1 and Figure 2A).

**Table 1:** List of upstream regulators identified by Ingenuity analysis of MS data. Shown are relevant gene ontology functions, the number of potential target proteins, that were changed following CDC treatment in LV from HFpEF rats compared to placebo, and the Ingenuity prediction for the upstream regulator to be activated or inhibited.

| Protein Name | Gene Name | Function | # Targets | Predicted Regulation by CDC |
| --- | --- | --- | --- | --- |
| Hepatocyte nuclear factor 4-alpha | HNF4A | Transcription factor | 125 | activated |
| Cellular tumor antigen p53 | TP53 | Cell cycle and apoptosis | 110 | inhibited |
| Transforming growth factor beta-1 proprotein | TGFβ1 | Cell growth and cell cycle | 86 | inhibited |
| Tumor necrosis factor | TNF | Inflammation | 85 | activated |
| Myc proto-oncogene protein | MYC | Cell cycle and apoptosis | 84 | N/A |
| Estrogen receptor | ESR1 | Transcription factor, cell proliferation and differentiation | 78 | N/A |
| Serine/threonine-protein kinase mTOR / Rapamycin-insensitive companion of mTOR | MTOR/RICTOR | Cell proliferation, survival, autophagy, and transcription | 56 | activated |
| Huntingtin | HTT | Cell signaling and transport | 53 | activated |

**Figure 2:**
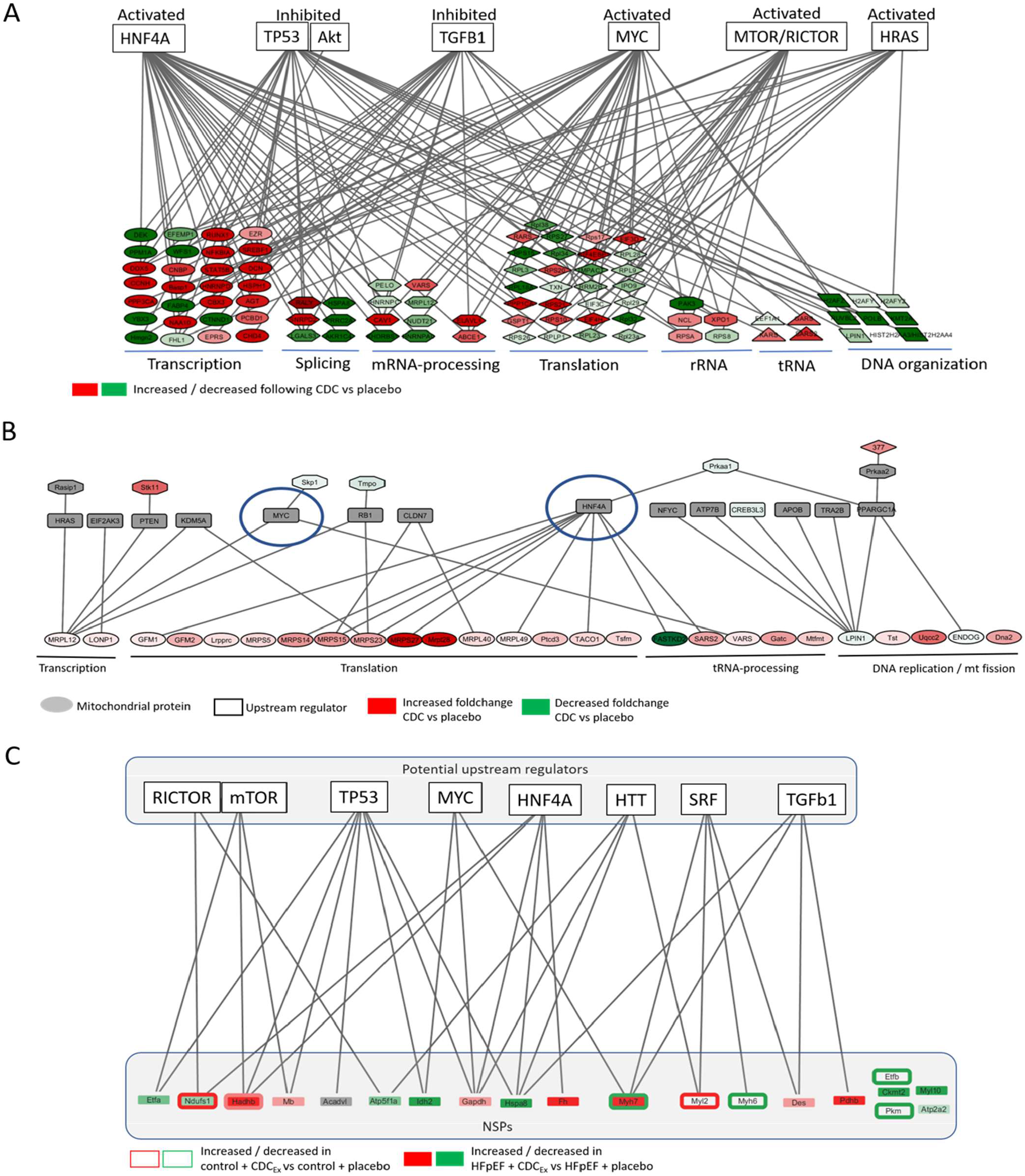
The graphic shows networks of potential upstream regulators identified based on the cohort of identified changed proteins from CDC treated HFpEF rats compared to placebo, that are able to influence protein expression (A), that are known to regulate mitochondrial protein expression and upstream regulators for NSPs significantly changed following CDCexo treatment of adult cardiomyocytes from HFpEF and control rats (C). The green color indicates a downregulation and the red color an upregulation. The intensity of the color represents the fold change intensity for CDC compared to placebo.

The specific quantification of NSPs in adult LV rat cardiomyocytes (LVMC) can be accomplished using AHA labeleing of proteins during their translation. AHA labeling of isolated LVMC revealed 16 proteins with significantly changed concentration following 5h of CDCexo treatment of LVMC isolated from HFpEF rats compared to 7 NSPs following CDCexo treatment in control cardiomyocytes (Table 2). Upstream regulator analysis revealed 7 upstream regulators that were predicted to regulate 75% of the CDC-induced NSPs. 5 were the same as those identified in the static proteome analysis to be involved intranscriptional and translational protein regulation, specifically mTOR, RICTOR, TP53, Myc, HNF4A, and Transforming growth factor beta-1 (TGFβ1) (Figure 2C).

**Table 2:** MS data of NSPs significantly changed following CDC treatment analysis. Shown are the foldchanges (log2) following CDCexo treatment of adult cardiomyocytes from LV of HFpEF abd control rats compared to placebo (n=6). If a NSP was detected soley in one group a foldchange (log2) of 4 was assigned.

| Protein Name | Gene Name | Cellular location | Foldchange (log2)<br>CDC+HFpEF vs<br>Placebo+HFpEF | p-value<br>CDC+HFpEF vs<br>Placebo+HFpEF | Foldchange (log2)<br>CDC+control vs<br>Placebo+control | p-value<br>CDC+control vs<br>Placebo+control |
| --- | --- | --- | --- | --- | --- | --- |
| Very long-chain specific acyl-CoA dehydrogenase, mitochondrial | Acadvl | Mitochondria | -4.00 | NA | NA | NA |
| Sarcoplasmic/endoplasmic reticulum calcium ATPase 2 | Atp2a2 | SR | -0.92 | 0.063 | NC | NS |
| ATP synthase subunit alpha, mitochondrial | Atp5f1a | Mitochondria | -1.84 | 0.022 | NC | NS |
| Creatine kinase S-type, mitochondrial | Ckmt2 | Mitochondria | -4.00 | NS | NC | NS |
| Desmin | Des | Myofilament | 1.39 | 0.024 | NC | NS |
| Electron transfer flavoprotein subunit alpha, mitochondrial | Etfa | Mitochondria | -2.01 | 0.006 | NC | NS |
| Electron transfer flavoprotein subunit beta | Etfb | Mitochondria | NC | NS | -1.86 | 0.033 |
| Fumarate hydratase, mitochondrial | Fh | Mitochondria | 4.00 | NA | NC | NS |
| Glyceraldehyde-3-phosphate dehydrogenase | Gapdh | Cytosol | 1.40 | 0.033 | NC | NS |
| Trifunctional enzyme subunit beta, mitochondrial | Hadhb | Mitochondria | 4.00 | NA | 0.48 | 0.025 |
| Heat shock cognate 71 kDa protein | Hspa8 | Cytosol | -4.00 | NA | NC | NS |
| Isocitrate dehydrogenase [NADP], mitochondrial | Idh2 | Mitochondria | -4.00 | NA | NC | NS |
| Myoglobin | Mb |  | 1.34 | 0.056 | NC | NS |
| Myosin-6 | Myh6 | Cytosol |  |  | -1.58 | 0.084 |
| Myosin-7 | Myh7 | Myofilament | 3.07 | 0.043 | -1.44 | 0.067 |
| Myosin regulatory light chain 10 | Myl10 | Myofilament | -4.00 | NA | NC | NS |
| Myosin regulatory light chain 2, ventricular/cardiac muscle isoform | Myl2 | Myofilament | NA | NA | 1.14 | 0.027 |
| NADH-ubiquinone oxidoreductase 75 kDa subunit, mitochondrial | Ndufs1 | Mitochondria | -1.40 | 0.006 | 0.85 | 0.005 |
| Pyruvate dehydrogenase E1 component subunit beta, mitochondrial | Pdhb | Mitochondria | 4.00 | NA | NC | NS |
| Pyruvate kinase PKM | Pkm | Cytosol | NC | NS | -1.67 | 0.060 |
| ADP/ATP translocase 1 | Slc25a4 | Mitochondria | -4.00 | NA | NC | NS |

NA = no appropriate NC = no change NS= not significant

The upstream regulatory analysis postulated that TP53 and TGFβ1 are inhibited following CDC treatment. Based on westerns blots (WB) of LV obtained from the various animal experimental groups, both TGFβ1 (−54.5 ± 11.3%, p<0.05) and in its downstream target SMAD2 (reduced by 22 ± 14%, p<0.05) were decreased in following CDC treatment (Supplemental figure 1A and B). As well, WB confirmed a reduction of TP53 in LV when compared to placebo (−20.3 ± 1.8%, p<0.05) along with a reduction in its phosphorylation at Ser15 (−22.2 ± 3.5%, p<0.05, Supplemental figure 1D and E). Our MS data further supports a reduction of TP53 activity as both Ampk and Hmgb1, known TP53 activators, are reduced in quantity (MS data showed a reduction by 30.4 ± 11.5% and 20.3 ± 10.5%, p<0.05, respectively, Supplemental table 1). On the other hand, based on the upstream regulatory analysis and according to the direction of their changed target proteins, AKT (also called protein kinase B or PKB) and HNF4A are postulated to be activated. The WB analysis supports a reduced total quantity of AKT following CDC treatment (−15 ± 9.8%, p<0.05) (Supplemental figure 1F). HNF4A and MTOR were increased in quantity based on WB in the LV obtained from CDC treated HFpEF compared to placebo(15 ± 3.2% and 98 ± 15.6%, respectively, p<0.05) (Supplemental figure 2H and 2K). Interestingly, the changes in the protein quantity of each upstream regulators correlates with the functional data for E/A ratio, Tau, and/or LVEDP. Figure 3 shows protein levels in each individual heart plotted against *in vivo* functional data obtained from each corresponding animal (based on WB). TGFβ1 positively correlated with LVEDP (R^2^=+0.59). TP53 positively correlated with Tau (R^2^=+0.622) and LEVDP (R^2^=+0.54), but it is the phosphorylated form of TP53 that is positively correlated with E/A ratio (R^2^=+0.53). HNF4A showed a positive correlation with E/A ratio (R^2^=+0.42) and a negative correlation with LEVDP (R^2^=0.-471). Myc negatively correlated with E/A ratio (R^2^=-0.43) (Supplemental figure 2).

**Figure 3:**
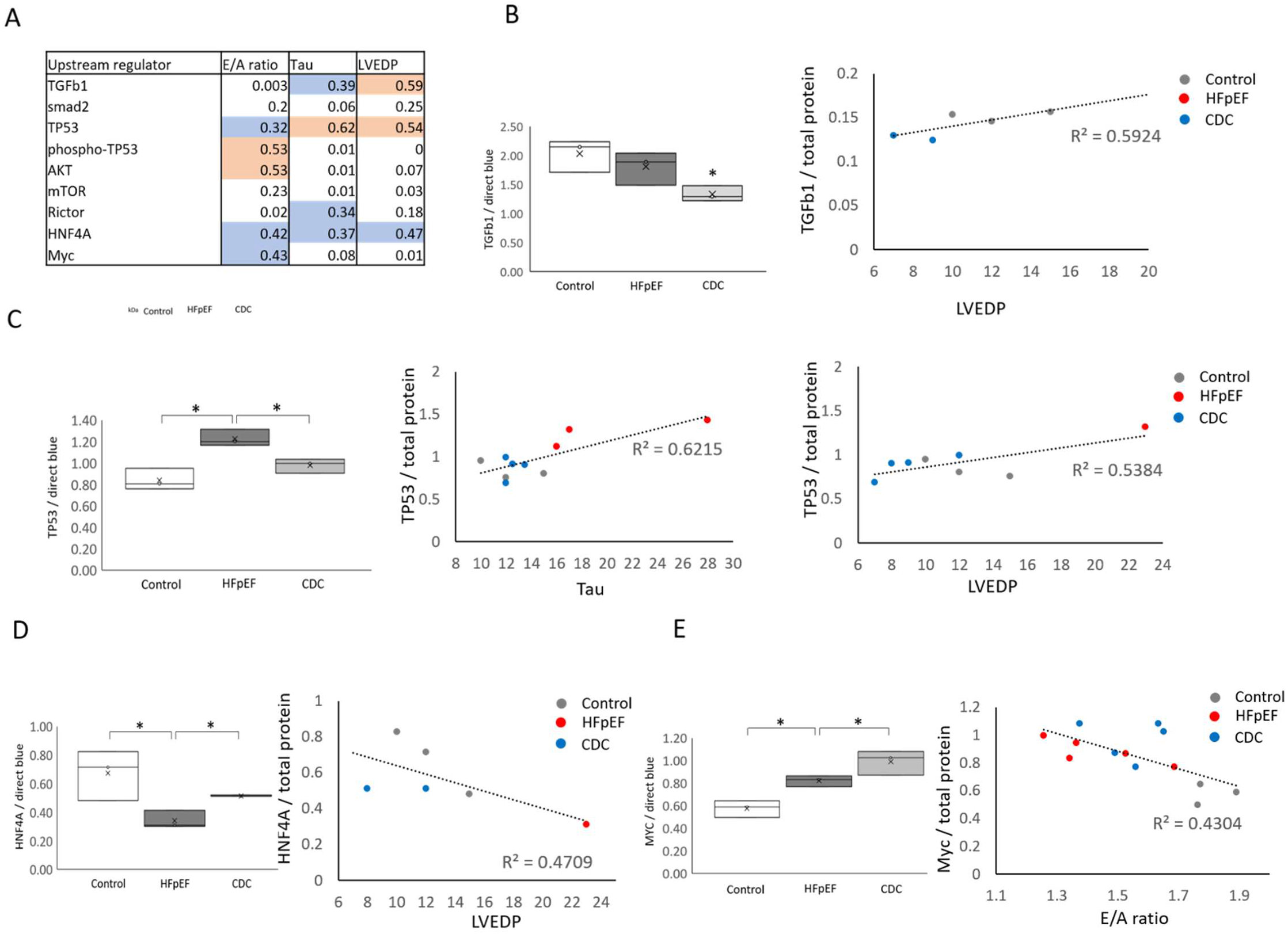
The Graphs are showing the correlation between functional data (E/A ratio, Tau, and LVEDP) and key regulatory protein concentrations (WB) of specific animals (HFpEF + placebo, HFpEF + CDC, and control) for TGFβ1 (B), TP53 (C), HNF4A (D) and Myc (E).

Besides protein regulating nuclear DNA encoded protein expression, there were an additional 26 DEPs following CDC treatment that are involved in mitochondrial DNA encoded protein expression. These proteins regulating mitochondrial transcription include cold shock domain-containing protein C2 (Csdc2), mitochondrial FAST kinase domain-containing protein (Fastkd2), histone-lysine N-methyltransferase 2A (Kmt2a), Heat shock cognate 71 kDa protein (Hspa8), Catenin delta-1 (Ctnnd1 or P120-catenin), RNA-binding protein Raly (Raly), Pterin-4-alpha-carbinolamine dehydratase 2 (Pcbd2), Splicing factor 1 (Sf1), and Brain acid soluble protein 1 (Basp1) (Supplemental table 4). Interestingly, upstream regulator analysis for the mitochondrial transcriptional regulatory proteins include HNF4A and Myc which could be directly involved in regulating 26% and 8% of proteins regulating mitochondrial protein expression, respectively (Figure 2B).

There was also extensive phosphorylation of proteins known to be involved in transcription (e.g. transcription factors, ribosomes) with an increase in over 100 modified residues in HFpEF rats compared to control animals and 32 of these changed phosphorylation with CDC compared to placebo treatment (Supplemental Table 2). A well-established algorithm (GPS 3.0) was used to identify potential kinases based on the probability of targeting the changes in phosphorylated residues plus information from the MS data (e.g. total or phospho-enriched peptide samples across treatment groups) and allows us to prioritize candidates based on the following criteria i) altered enzyme concentration, ii) phosphorylation changes, iii) alterations in known kinase regulatory proteins, and iv) the amount of potential target phospho-sites in diastolic dysfunction with CDC-treatment compared to control (Figure 4A). 5 kinases (Protein kinase C (PKC), AKT, mTOR, Glycogen synthase kinase (GSK) and Protein kinase A (PKA)) were identified within our proteomic dataset to be regulated in LVs obtained from the CDC treated HFpEF compared to placebo treated group. Collectively, they share consensus amino acid sequences with 31 of those phospho-sites. PKC was the most dominant kinase potentially targeting 90.6% of the CDC -induced changes phospho-sites (90.6%). For detail supporting the role specific kinases based on proteomic data (e.g. quantity or phosphorylation status of a specific kinase or one of their regulatory proteins) see supplemental table 5.

**Figure 4:**
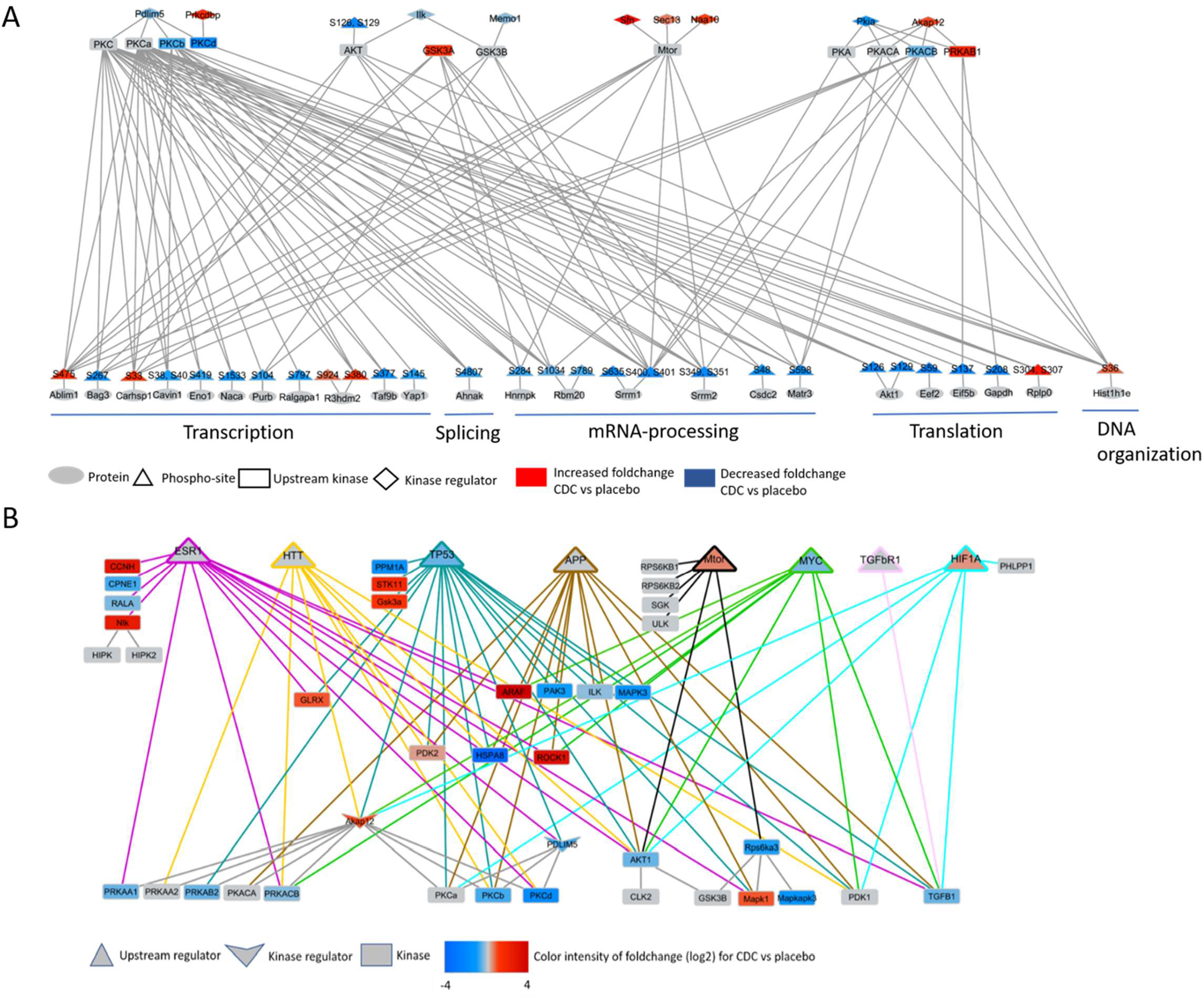
Network of potential upstream kinases with consensus sequences for altered phosphorylated proteins regulating protein expression in HFpEF following CDC treatment compared to placebo. Only kinases with MS data support for their changed activity concentration, phosphorylation or known regulators were included in the figure (A). A upstream regulator analysis of the mot relevant kinases with the most identified shared consensus sequences with CDC changed phosphorylation sites was completed. Shown are the most relevant potential upstream regulators that can regulate kinases of relevance for regulating protein expression (B).

In order to identify regulatory elements within the CDCexo cargo that are able to trigger these key regulators, TP53, mTOR, MYC, TGFβ1, and HNF4A, the isolated CDCexo were also analyzed by LC-MS (Supplemental table 5). In isolated CDCexo,there has been approximately 400 unique proteins and 229 different miRNAs previously identified[27]. Of these, 31 proteins and 6 miRNAs have been linked directly to the regulation of the 6 upstream regulators predicted to mediate the CDC response (Figure 5 and supplemental table 6). To validate these CDCexo-biologics, we quantified NSPs, using AHA labeling, following inhibition of multiple different PKC isoforms using 3 different small molecule inhibitors for PKCα, PKCβ, and PKCδ in H9C2 cells. The inhibition of PKCα, beta, and delta led to a significant change of 12, 25 and 15 proteins (with 22 proteins, 57 and 25 protiens trending, p<0.1), respectively. Furthermore, 44% of the NSPs changed following PKC inhibition were also changed following CDC treatment of HFpEF rats with PKCβ inhibition having the highest impact on cellular protein synthesis (Supplemtal figure 4 and supplemental table 6).

**Figure 5:**
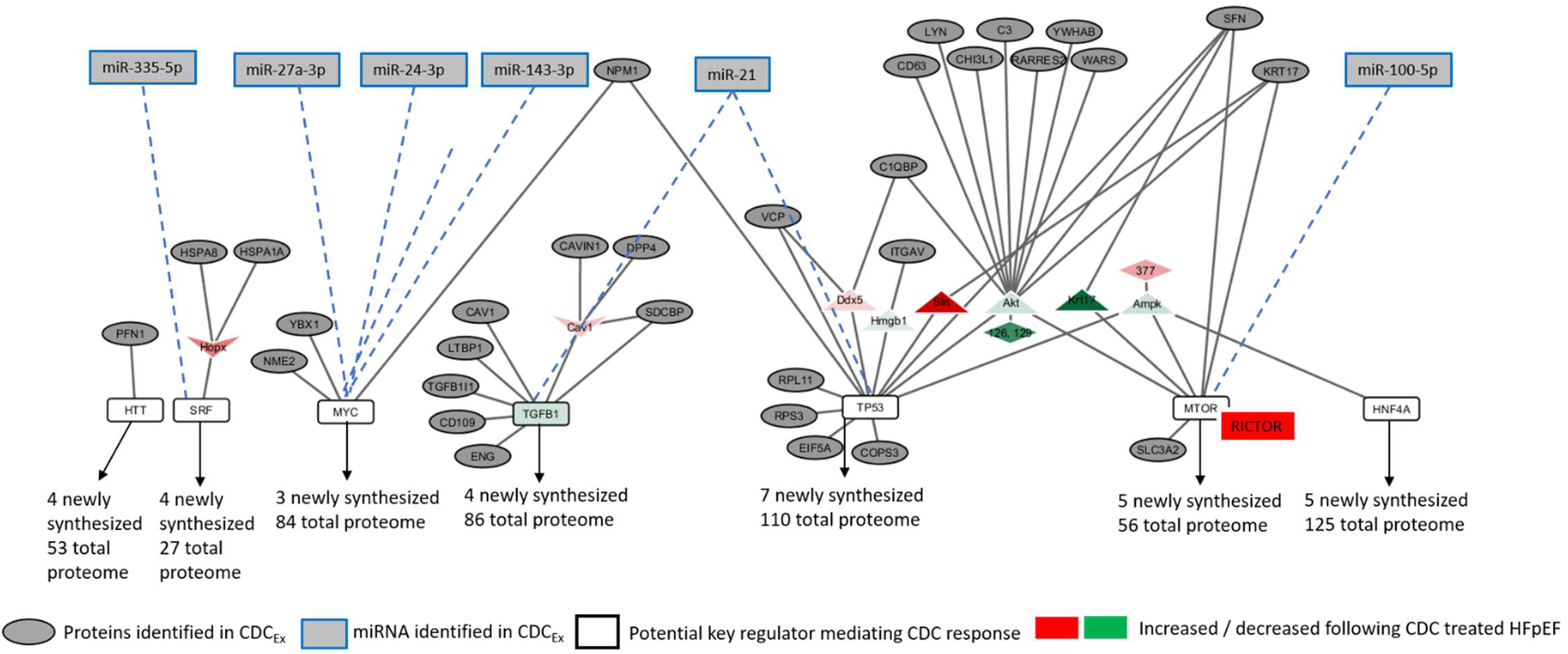
Network of proteins and miRNA identified in CDCexo cargo that are known to regulate the identified key regulators of CDC therapy. The green color indicates a downregulation and the red color an upregulation following CDC treatment. The intensity of the color represents the fold change intensity for CDC compared to placebo.

We performed upstream regulator analysis of those kinases to identify those that could induce the extensive phosphorylation. TP53 can potentially target 14 relevant kinases, among them GSK3A, PKA, PKCβ, and TGFβ1. Myc can potentially target 7 kinases, e.g. AKT1, PDK1, PKA, and also TGFβ1. TGFβ1 as well, could regulate 8 kinases, e.g. AKT1, and PKCα (Figure 4C). Interestingly, miR-142-3p which was found in CDCexo, was also identified as an upstream regulator for PKCα.

## Discussion

In this study, we dissected the underlying proteomic changes reprogramming the transcriptional- and translational machinery following CDC treatment of HFpEF, and revealed a number of novel findings including i) extensive changes in concentration and phosphorylation of proteins regulating nuclear and mitochondrial protein expression; ii) quantification of NSPs following CDCexo treatment revealing 5 potential key mediators of CDC therapy; iii) characterization of the CDCexo cargo revealing 24 proteins and 4 miRNAs that are known regulators for the 5 identified key mediators of CDC response. Furthermore, 4 of these key upsteam regulator were also perturbed in the rat HFpEF model and recapitulated in the human HFpEF hearts when compared to healthy controls supporting that reversal or modulation of these pathways could be important in human disease.

Briefly, of the large proteome changes with rat HFpEF model compared to control animals, the majority (348 DEPs) are involved in regulating protein expression and CDC treatment reversed 125 of these back towards levels found in healthy animals. The proteomic profile suggests that cellular functions like fibrosis, inflammation, and cell death are increased, whereas other cellular functions like calcium handling, contraction, myofilament compositon, and protein transport seem dysregulated in rat HFpEF. In comparison the proteomic profile of human HFpEF showed strong similarities regarding physiological cellular functions, e.g. fibrosis, metabolism, inflammation, cell death, and calcium handling. Further, the proteomic profile of human HFpEF indicates, that TP53 signaling is congruently regulated compared to the rat HFpEF model.

Importantly, the CDC-sensitive DEPs in the HFpEF rat model were also enriched in transcription and translation and involved in regulation of both nuclear and mitochondrial encoded proteins suggesting a potential coordination. Interestingly a large portion of the regulated proteome is associated with cell cycle regulation, inflammatory signaling and cell death. These include proteins like Cold-inducible RNA-binding protein (Cirbp; increased following CDC treatment) and Eukaryotic translation initiation factor 4E-binding protein 1 (Eif4ebp1; increased following CDC treatment), known for positive regulation of cell cycle G1/S phase transition [28, 29], whereas core histone macro-H2A.1 (H2afy; decreased following CDC treatment) negatively regulates the cell cycle G2/M phase transition [30]. Besides their role as a transcriptional co-activator, high mobility group protein B1 and B2 (Hmgb1 and Hmgb2; both decreased following CDC treatment) are associated with activation of innate immune response, negative regulation of apoptotic cell clearance, and pro-inflammatory response [31-34]. Elongation factor 1-alpha 1 (Eef1a1; decreased following CDC treatment) positively regulates cell death [35] and eukaryotic translation initiation factor 3 subunit C (Eif3c) including cell cycling, differentiation and apoptosis (Supplemental table 1) suggesting that CDC induced reprogramming of the transcriptional and translational machinery activates cell proliferation, reduces inflammation and cell death, which is supported by several studies investigating CDC therapy [20, 27].

The quantification of NSPs is a read out of transcription and translational alterations found in HFpEF and then altered following 5 h of CDCexo treatment. These proteins are mainly those that have a rather rapid turnover, like mitochondrial proteins (ATP synthase subunit alpha, NADH-ubiquinone oxidoreductase 75 kDa subunit, and ADP/ATP translocase 1), myofilaments (Desmin, Myosin regulatory light chain 2, ventricular/cardiac muscle isoform, and Myosin-6), and metabolic enzymes (Glyceraldehyde-3-phosphate dehydrogenase, Isocitrate dehydrogenase [NADP], and Fumarate hydratase) (Table 2). The data limitations quantifying NSPs in adult cardiomyocytes arose due to the fragile state of cardiomyocytes isolated from HFpEF rats that survive for only 12 h in culture leaving a short timeframe for capturing NSP and thus explaining the low amount of significantly changed NSP. Nevertheless, using this data as a bases for identification of upstream regulators using Ingenuity upstream regulator analysis confirmed 5 proteins of the 7 that were also identified for regulating the changes of the transcriptional and translational machinery.

PKC, GSK, mTOR, AKT, and PKA are the dominant kinases predicted based on bioinformatic consensus sequence analysis to be involved in the large reversal of the phosphorylation status of HFpEF that occur following CDC treatment. Specifically, PKC isoforms α and β share the majority consensus sequences with CDC treatment changed phospho-sites (90%). It is worth mentioning that the predominant kinases with the most potential phospho-targets are also known potential downstream targets of mTOR, MYC, and/or TP53 or vice versa. For instance, PKCα protects cells from apoptosis by suppressing the TP53-mediated activation of IGFBP3 [36] and PKCβ is likely involved in the inhibition of the insulin gene transcription, via regulation of MYC expression [37, 38]. Interestingly, the *in vitro* experiments performed in this study quantifying NSPs following PKC isoform inhibition showed a strong overlap with the proteomic data acquired from CDC treated HFpEF rats, especially for PKCβ. It is possible, that PKC functions as a downstream mediator within CDC therapy regulating protein expression and thus could be an interesting target for a CDC replacement therapy.

It is of interestwhether these primary tissue regulators, TGFβ1, TP53, Myc, MTOR and HNF4A, are correlating with specific dysfunctions seen in HFpEF (including TP53 in human HFpEF) and if those functional parameters are reversing in correlation those primary regulaters following CDC treatment. For example, the quantity of TGFβ1 in each of the animals used in this study positively correlate with Tau and LVEDP leading to the assumption that CDC induced reduction of TGFβ1 signaling is contributing to a reduction of myocardial stiffness and thus reducing the accompanying increased LVEDP in HFpEF. It is known that excessive TGFβ1 signaling is contributing to pathologic fibrosis, scarring and matrix deposition in a variety of diseases including diabetes, cardiac hypertrophy, and heart failure [39-42]. Gallet et. al. concluded that CDC treatment improved diastolic function through limiting fibrosis and inflammation in this rat model [20]. In line with this conclusion, the proteomic data of this study shows a reduction of TGFβ1 following CDC treatment. Upon TGFβ1 secretion the downstream cellular signaling is dependent on TGFβ1 receptors, serine-threonine kinases, and SMAD proteins. SMADS are involved in the formation of functional transcription complexes inducing and mediating the TGFβ signal to target genes [43-45]. Here we also showed, that SMAD2 was reduced following CDC treatment suggesting a CDC induced downregulation of TGFβ1 signaling. Our data suggests that CDC treatment is contributing to a normalization of the HFpEF phenotype by inhibiting TGFβ1-signalling, which could be achieved via miR-21 and miR-143-3p identified in CDCexo. It is shown, that miR-21 improves LV remodeling and decreases apoptosis in myocardial cells [46]. Recently, treatment with miR-21 post-MI has been shown to reduce hypertrophy, fibrosis, and cell apoptosis, whereas it promoted angiogenesis in the remote myocardium. Thus suggesting that miR-21 has therapeutic capacities that are o relevance or post-MI remodeling and heart failure [47]. Further, miR-21 has a role in adipogenic differentiation mediated through modulation of TGFβ-signaling [48]. But its role in fibrosis in remains controversial. Overexpression of miR-21 in fibroblasts promoted TGF-β1-induced expression of Collagen-1, alpha-smooth muscle actin (α-SMA), and F-actin, whereas inhibition of miR-21 attenuated the fibrotic process [49].

Interestingly, MiR-143-3p suppresses proliferation and ECM protein deposition by inhibition of TGFβ1 via the negative regulation of NFATc1 signaling in airway smooth muscle cells [50]. Additionally, some proteins identified in the CDCexo cargo are known to negatively regulate TGFβ1 signaling, e.g. CD109 antigen (CD109) (inhibits TGFβ in human keratinocytes) [51] and Caveolin-1 (CAV1) (internalizes the TGFβ-receptor 1 from membrane rafts) [52]. Other protein identified in CDCexo positively regulate TGFβ1 signaling, e.g. Latent-transforming growth factor beta-binding protein 1 (LTBP1) (regulates integrin-dependent activation of TGFβ) [53-55], Endoglin (ENG) (involved in the TGFβ/BMP signaling cascade leading to the activation of SMAD transcription factors) [56-59], and Integrin alpha-V (ITGAV) (activates TGFβ1 via R-G-D-dependent release of TGFβ1 from regulatory Latency-associated peptide) [55, 60, 61]. It remains to be elucidated, whether the CDCexo protein cargo is sufficient to alter host cell signaling.

TP53 quantity in LV of rats positively correlated with Tau and LVEDP whereas levels of activated phosphorylated TP53 (Ser15) positively correlated with the E/A ratio strongly suggestive of a beneficial improvement of diastolic function with a CDC mediated reduction of TP53. TP53 was found to be overexpressed in myocytes of patients with dilated cardiomyopathy [62]. TP53 is involved in regulating cell cycle inhibiting cell division via controlling crucial genes. The CDC mediated downregulation of TP53 could reduce cell death and increase cell division in HFpEF. Following CDC treatment mTOR was increased and RICTOR, a subunit of mTOR complex 2 regulating cell growth and survival, was strongly increased following CDC treatment. The mTORC1 pathway regulates protein synthesis and cell growth promoting proliferation in response to mitotic signals [63, 64]. The balance of the stress-response and pro-growth pathway is important for optimal cell proliferation. Thus, our data suggests CDC treatment leads to reduced cell death, increased cell division and cell growth. The exact mechanism involving CDC regulation of the mTOR/TP53 pathway remains to be investigated. Additional supporting information for a CDC mediated downregulation of TP53 and mTOR was found. Several TP53 and mTOR regulators were changed following CDC treatment. Among them Hmgb1 which facilitates binding of TP53 to DNA [65]. AKT was changed following CDC, which activates mTORC1 signaling via TSC2 phosphorylation [66]. Further, Ampk was altered following CDC. The kinase phosphorylates transcription regulator TP53 and negatively regulates the mTORC1 complex by phosphorylating RPTOR and mTORC1 inhibitor TSC2 [67]. Thus, the identified downregulation of TP53 and upregulation of mTOR following CDC therapy could be achieved via a downregulation of AKT. The group of proteins involved in p53-AKT-mTOR signaling seems to be especially strong targeted by the CDCexo with 18 identified biological compounds that are known regulators. NPM1 an activator of cell cycle G2/M phase transition was identified in CDCexo, regulating TP53 [68, 69]. COPS3 a component of the COP9 signalosome complex targets p53 for degradation by the ubiquitin system [70]. C1QBP is involved in the inhibition of innate immune response via the PI3K-AKT/PKB pathway [71], SFN activates TP53 via MDM2 degradation, and when bound to KRT17, regulates protein synthesis and epithelial cell growth by stimulating AKT/mTOR pathway [72]. Interestingly, AKT is downregulated following CDC treatment. The CDCexo cargo carried 10 proteins that activate AKT. Further research is needed to investigate the CDC mediated fine tuning of this pathway. We also identified miR-100-5p and miR-21 within CDCexo. Studies showed that those microRNAs regulate the PI3K/AKT/mTOR pathway. MiR-100-5p affects the mTOR mediated autophagy [73] and MiR-21 plays a role in skeleton muscle development via targeting the TGFβ1 expression [74].

Two additional transcription factors, Myc and HNF4A, have not been well studied in HFpEF yet are indicated in this work to play a role in mediating CDC therapy. Studies showed that Myc activates growth-related genes [75] and regulates the self-renewal of embryonic stem cells [76]. In this study, Myc (which is sensitive to CDC treatment) could be regulated via 3 proteins and 4 miRNAs identified within CDCexo. The microRNAs, miR-24-3p, miR-27a-3p, miR-143-3p, and miR-34a have been reported to target the Myc gene [77-80]. Studies showed that miR-24-3p regulates cell proliferation (mESC and human smooth muscle cells) via inhibition of Myc at the post-transcriptional level [81, 82]. On the other hand, miR-24-3p and miR-27-3a were found to promote cell proliferation in glioma cells via inhibition of Myc antagonist MXI1, a member of Mad (Mxi1) family [77]. MiR-143-3p directly targets ERK5 and initiates cell-growth via activation of Myc [83] and is involved in apoptosis (miR-143-3p inhibits the proliferation, migration and invasion in osteosarcoma by targeting FOSL2). Overexpression of miR-34a decreased MYC levels in lymphoma cell lines leading to inhibition of cell proliferation, and induced apoptosis [84, 85]. Proteins identified within CDCexo that regulate Myc include: a transcriptional activator for the Myc gene nucleoside diphosphate kinase B (NME2) [86, 87], Y-box-binding protein 1 (YBX1) stabilizing Myc mRNA [88], and NPM1 which in complex with Myc increases its activity and enhances transcription of Myc target genes [89]. HNF4A was reduced in HFpEF rats and increased to control levels with CDC therapy. This study showed a negative correlation of HNF4A concentrations with Tau and LVEDP suggesting that a CDC mediated increase in HNF4A contributes to a reduction in cardiac stiffness and diastolic pressure. Transcription actor HNF4A is not well-studied,thus, it is not surprising, that we weren’t able to identify potential regulators within CDCexo. Further, research is needed to characterize HNF4A role in mediating CDC therapy.

There are several limitations of our study that are noteworthy. The results we present here are correlational and indirect without demonstrating that specific manipulation of identified key mechanisms are impacting HFpEF pathology or will alter myocardial function. The primary upstream regulators are inferred based on global proteome changes and quantities were validated by WB. The accuracy of the protein measurements is influenced by both analtyical and biological variability. With human disease there are additional confounders with respect to age, enthicity, and therapeutic state which could explain network directionality differences between human HFpEF and the rat HFpEF model.

In conclusion our data suggest that CDC therapy is associated with i) reprogramming of cardiac cells via alterations within the transcriptional and translational machinery; ii) seven potential cellular key regulatory elements targeted by CDC therapy; iii) for 5 of the 7 potential key elements numerous proteins and miRNA were identified within the CDCexo cargo, that are documented to play a role in their regulation. The most promising targets for a CDC replacement therapy would be i) TP53 and mTOR signaling regulating cell cycle, cell growth and apoptosis in an opposing manner and ii) TGFβ1 which is known to play a role in cardiac disease and cell proliferation and fibrosis was reduced following CDC treatment in our MS data. Taken together with the two identified upstream regulators MYC and HNF4A we identified 4 key regulators mediating CDC therapy. Those 4 key regulators are very promising targets for a CDC a replacement therapy for treating HFpEF and other related diseases (Figure 6).

**Figure 6:**
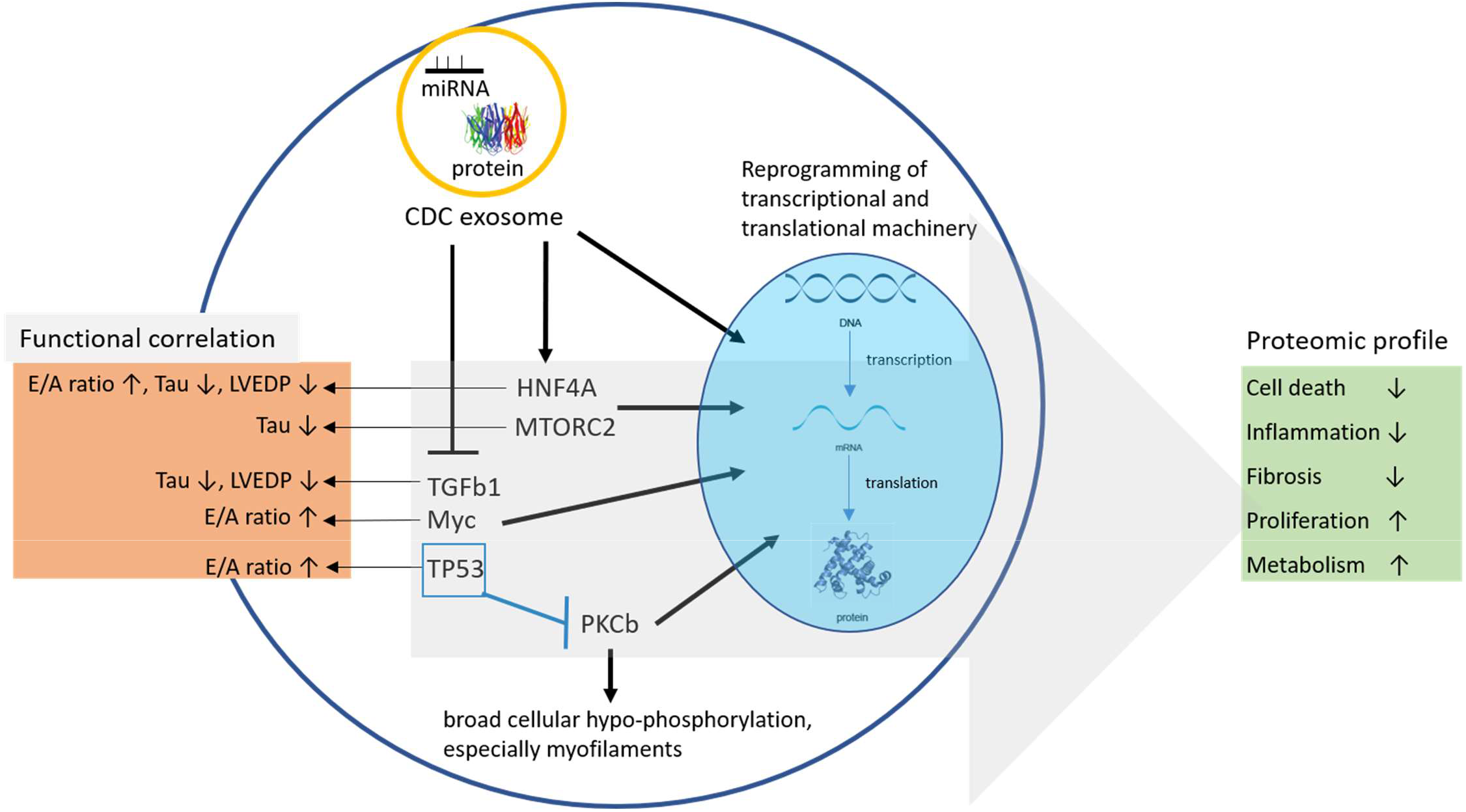
Overview of discovered key elements of CDC induced transcriptional and transitional reprogramming of HFpEF.

## Abbreviations

AHA: L-azidohomoalanine
Ca^2+^: Calcium
CDC: Cardiosphere-derived cell
CDCexo: CDC exosomes
DEP: differential expressed protein
DMEM: Dulbecco’s Modified Eagle Medium
E/A: ratio
EV: extracellular vessicle
FC: Fold change
HF: Heart failure
HFpEF: Heart failure and preserved ejection fraction
HFrEF: Heart failure with reduced ejection fraction
LC-MS: Liquid chromatography-mass spectrometry
LV: Left ventricle
LVEDP: left ventricular end-diastolic pressure
MS: Mass spectrometry
mTOR: mammalian target of rapamycin
Myc: Myc proto-oncogene protein
NRVMs: Neonatal rat ventricular myocytes
NSPs: Newly synthesized proteins
PE: Phenylephrine
PKC: Protein kinase C
RICTOR: mammalian target of rapamycin
Ser: Serine (S)
SS: Salt sensitive
TGF β-1: Transforming growth factor beta-1
Thr: Threonine (T)
TP53: Cellular tumor antigen p53
Tyr: Tyrosine (Y)
WB: Western blot

## Acknowledgements and Sources of Funding

This work was generously supported by a U.S. Department of Defense Grant to E.M, MZ and J.E.V.E.; the National Institutes of Health grants T32 HL116273 to J.I.G. and E.M.; American Heart Association generously support this work with a Postdoctoral Fellowship Award to D.S.; Smidt Heart Institute E.M.; the Erika J. Glazer chair in Women’s Heart Health to J.E.V.E.; the Barbra Streisand Women’s Heart Center to J.E.V.E.; the Advanced Clinical Biosystems Institute to J.E.V.E. at Cedars-Sinai Medical Center; and the VA Merit CX001608 to A.B.; R01 HL123478-01 to M.K., RO1 HL144927 to M.K., and Department of Defence Grant W81XWH-16-1-0592 to M.K.

