## Supplementary material for "Mechanistic dissection of global proteomic changes in rats with heart failure and preserved ejection fraction": Online supplements

**Online Supplements – Methods**

**Sample handling and preparation for proteomics**

Freshly harvested rat left ventricle (LV) tissue was quickly rinsed in ice-cold homogenization buffer (300 mM sucrose, 250 mM HEPES-NaOH, 1 mM EDTA, pH 8.0) supplemented with a broad-spectrum protease inhibitor (Thermo Fisher Scientific, Waltham, MA) for total protein analysis, or phosphatase inhibitor cocktail (Thermo Fisher) for protein phosphorylation were snap frozen in liquid nitrogen. Left ventricle tissue was fractionated into cytosolic-, myofilament-, and in-soluble-enriched fractions by the "in sequence" method as previously described [1]. The protein concentration was determined by BCA (Pierce) and one hundred μg of each fraction (total protein quantification) or 1 mg of protein of each fraction (phosphorylation quantification) were reduced using 1 mM TCEP, and alkylated using 4 mM iodoacetamide in a filter-aided sample preparation (FASP 10 kDA, Promega). Samples were digested for 15–18h at 37°C using ultra-grade Trypsin (Promega) at a 1:100 enzyme: protein ratio. Samples were desalted using Oasis HLB plates (30 μm and 5 mg sorbent, Waters), vacuum dried and stored at -80°C until analysis. For analyzing phosphorylated peptides, samples were enriched post-digestion and desalted using $TiO_2$ beads as previously described [2].

**Mass spectrometry and data analysis**

Tissue and cell culture samples were analyzed by liquid chromatography-tandem mass spectrometry (LC-MS/MS) on a Dionex Ultimate 3000 NanoLC connected to an Orbitrap Elite (Thermo Fisher Scientific) equipped with an EasySpray ion source or on a Dionex Ultimate 3000 NanoLC connected to an Orbitrap Fusion™ Lumos™ Tribrid™ Mass

Spectrometer (Thermo Fisher Scientific) equipped with an EasySpray ion source as previously reported [3].

Raw MS/MS data files were converted to mzXML format using MSconvert version v.3.0.6002 from ProteoWizard [4] for peaklist generation and used for database searching using two engines, X!Tandem [5] algorithm version 2013.06.15.1 and Comet [6] algorithm version 2014.02 rev.2. The dataset was searched against the concatenated target/decoy [7] rat (8,231 proteins) and mouse (32,741 proteins on March 06, 2017) in the Uniprot database [8] and additionally included non-reviewed protein IDs of the rat database (163,172 proteins on April 07, 2016). The latter was required as some key contractile proteins were missing from the annotated rat database (e.g. titin). The uniqueness of identified sequences assigned to non-reviewed proteins was validated by manual Bastp search, thus filtering multiple assigned peptides. Protein isoforms were only reported if a peptide comprising of an amino acid sequence that is unique to the isoform was identified. The MS/MS proteomics data will be deposited in the ProteomeXchange Consortium (http://www.proteomexchange.org) via the PRIDE partner repository [9].

Protein quantification was carried out by averaging the raw peptide intensity among technical replicates and summed among cellular sub-fractions. The peptide signal intensity was normalized to the median of the overall sample signal intensity and protein level abundance inference was calculated using the linear mixed effects model built into the open sources MSSTATs (v3.2.2) software suite [10]. For quantification of PTMs, intensities were normalized to the relative signal intensity of sample loading instead. Phosphorylated residues of peptides with missed cleavages or different charge states were summed before comparing MS1 intensity. Phosphorylation was quantified by modified (+80) peptide abundance. Statistical significance was determined at a nominal *p-value* ≤0.05. Upstream kinases potentially targeting significantly changed phosphorylated residues were identified using the Group-based prediction system v3.0 software.

**Western blot**

A total of 30 μg of frozen LV rat tissue (n=3) was minced and lysed in 2% SDS using a glass homogenizer and cell debris pelleted by centrifugation at 13,000g for 10 min at 4°C. Protein concentration was determined using BCA assay as above, Samples were separated by electrophoresis on a 4–10% gradient Bis–Tris SDS-Gels (Bio-Rad) and transferred to PVDF membranes and control stained with direct blue. After blocking, membranes were incubated with primary antibodies: rabbit anti-Rictor (2140, Cell Signaling), mouse monoclonal anti-HNF-4 α (ab41898, Abcam), rabbit anti-Mtor (2972, Cell Signaling), rabbit anti-phospho-mTOR (Ser2448) (2971, Cell Signaling), rabbit anti-Akt (9272, Cell Signaling), rabbit monoclonal anti-phospho-Akt I (phospho S473) (ab81283, Abcam), rabbit polyclonal anti-p53 (ab131442, Abcam), rabbit anti-Phospho-p53 (Ser15) (9284, Cell Signaling), rabbit monoclonal anti-c-Myc (ab32072, Abcam), rabbit anti-Smad2 (3122, Cell Signaling), rabbit anti-phospho-Smad2 (Ser465/467) (138D4, Cell Signaling), and mouse monoclonal anti-TGF-β I (MAB240, R&D Systems). The corresponding IgG HRP-conjugate combined with chemiluminescent substrate (Bio-Rad) and was scanned by a luminescent image quantifier and was within the linear range (ImageQuant LAS 4000, GE Healthcare). Protein quantity was normalized to direct blue staining, phosphorylation was normalized to the total of the same protein. An independent two-tailed t-tests ($p<0.05$) were used to compare experimental differences.

### Online Supplements – Results

**Supplemental table 1:** List of MS data of proteins able to regulate protein expression. Shown are the fold change values (log2) of HFpEF compared to control and for CDC compared to placebo treated HFpEF for total protein concentration (n=3–4).

| Protein name | Gene name | Function | Fold-change (log2) HFpEF vs control | p-value HFpEF vs control | Fold-change (log2) CDC vs placebo | p-value CDC vs placebo |
|---|---|---|---|---|---|---|
| Alanine--tRNA ligase, cytoplasmic | Aars | Translation, tRNA | 0.37 | 0.018 | NC | NS |
| Alanyl-tRNA editing protein Aarsd1 | Aarsd1 | Translation, tRNA | -0.19 | 0.079 | 0.20 | 0.054 |
| Protein ABHD14B | Abhd14b | Transcription | -0.43 | 0.010 | NC | NS |
| Actin-binding LIM protein 1 | Ablim1 | Transcription | -0.88 | 0.002 | 0.62 | 0.012 |
| Double-stranded RNA-specific editase B2 | Adarb2 | mRNA processing | NC | NS | -1.81 | 0.000 |
| Proteasomal ubiquitin receptor ADRM1 | Adrm1 | Transcription | NC | NS | 0.58 | 0.024 |
| Agrin | Agrn | Transcription | -0.73 | 0.005 | NC | NS |
| Angiotensinogen | Agt | Transcription | -0.97 | 0.000 | 0.48 | 0.005 |
| Estradiol 17 beta-dehydrogenase 5 | Akr1c6 | Splicing | 3.66 | 0.000 | -1.87 | 0.005 |
| RNA demethylase ALKBH5 | Alkbh5 | mRNA processing | -3.03 | 0.059 | NC | |
| Aly/REF export factor 2 | Alyref2 | Nuclear export | -1.87 | 0.011 | 1.59 | 0.028 |
| Acidic leucine-rich nuclear phosphoprotein 32 family member E | Anp32e | Histone | 0.75 | 0.038 | NC | NS |
| Annexin A3 | Anxa3 | Transcription | 0.36 | 0.002 | NC | NS |
| C->U-editing enzyme APOBEC-2 | Apobec2 | DNA demethylation, mRNA processing | 0.47 | 0.001 | NC | NS |
| Androgen receptor | Ar | Transcription | 2.93 | 0.000 | NC | NS |
| ADP-ribosylation factor 4 | Arf4 | Transcription | 0.69 | 0.003 | NC | NS |
| Ataxin-2-like protein | Atxn2l | mRNA processing | 1.45 | 0.000 | -1.16 | 0.000 |
| Barrier-to-autointegration factor | Banf1 | DNA organization | 1.82 | 0.000 | -0.57 | 0.068 |
| Brain acid soluble protein 1 | Basp1 | Transcription | -3.18 | 0.055 | 3.75 | 0.008 |
| Breast cancer type 1 susceptibility protein homolog | Brca1 | Nuclear import, Transcription | -1.54 | 0.002 | NC | NS |
| Basic leucine zipper and W2 domain-containing protein 1 | Bzw1 | Transcription | -0.56 | 0.011 | NC | NS |
| Basic leucine zipper and W2 domain-containing protein 2 | Bzw2 | Translation | -0.45 | 0.005 | NC | NS |
| Calcium-regulated heat stable protein 1 | Carhsp1 | mRNA processing | 0.42 | 0.010 | NC | NS |

| | | | | | | |
|---|---|---|---|---|---|---|
| Caveolin-1 | Cav1 | mRNA processing | -1.37 | 0.000 | 0.80 | 0.000 |
| Cyclin-H | Ccnh | Transcription | -3.45 | 0.000 | 2.63 | 0.000 |
| Cyclin-L1 | Ccnl1 | Splicing | -0.81 | 0.023 | NC | |
| Cadherin-13 | Cdh13 | Transcription | NC | | -0.41 | 0.001 |
| CUGBP Elav-like family member 1 | Celf1 | Splicing | 0.57 | 0.013 | NC | NS |
| Centrosomal protein of 290 kDa | Cep290 | Transcription | -2.53 | 0.000 | NC | NS |
| Cellular nucleic acid-binding protein | Cnbp | Transcription | 1.21 | 0.000 | NC | NS |
| Cellular nucleic acid-binding protein Isoform 2 | Cnbp | Transcription | 0.86 | 0.000 | 0.41 | 0.037 |
| Collagen alpha-1(I) chain | Col1a1 | Transcription | 2.73 | 0.023 | NC | |
| Copine-1 | Cpne1 | Transcription | NC | | -0.70 | 0.001 |
| Cyclic AMP-responsive element-binding protein 3-like protein 3 | Creb3l3 | Transcription | 0.78 | 0.035 | -0.57 | 0.09 |
| cAMP-responsive element modulator | Crem | Transcription | 1.72 | 0.002 | -0.85 | 0.066 |
| CREB-regulated transcription coactivator 2 | Crtc2 | Transcription | 1.30 | 0.013 | NC | NS |
| Ketimine reductase mu-crystallin | Crym | Transcription | NC | NS | 1.00 | 0.001 |
| Cold shock domain-containing protein C2 | Csdc2 | Transcription, mRNA processing, mRNA stabilization | NC | NS | -7.11 | 0.000 |
| Cysteine and glycine-rich protein 3 | Csrp3 | Transcription | 1.54 | 0.000 | -0.63 | 0.000 |
| Catenin delta-1 | Ctnnd1 | Transcription | 1.53 | 0.002 | -2.66 | 0.000 |
| Aspartate--tRNA ligase, cytoplasmic | Dars | tRNA | NC | | 0.25 | 0.020 |
| Decorin | Dcn | Transcription | -1.39 | 0.000 | 0.64 | 0.022 |
| ATP-dependent RNA helicase DDX1 | Ddx1 | Transcription, mRNA processing, Splicing, RNA helicase | -0.27 | 0.006 | NC | NS |
| ATP-dependent RNA helicase DDX39A | Ddx39a | Splicing, Nuclear export, RNA helicase | -0.70 | 0.000 | NC | NS |
| Spliceosome RNA helicase Ddx39b | Ddx39b | Transcription, Splicing, Nuclear export, Translation, RNA helicase | -0.50 | 0.000 | NC | NS |
| Probable ATP-dependent RNA helicase DDX4 | Ddx4 | DNA methylation, RNA helicase | -0.86 | 0.005 | 0.55 | 0.042 |
| Probable ATP-dependent RNA helicase DDX5 | Ddx5 | Transcription, RNA processing, Splicing, RNA helicase | -0.88 | 0.000 | 0.59 | 0.003 |
| Protein DEK | Dek | Transcription | 2.16 | 0.000 | -1.00 | 0.017 |

| | | | | | | |
|---|---|---|---|---|---|---|
| Density-regulated protein | Denr | Translation initiation | NC | NS | -0.28 | 0.001 |
| Deoxyguanosine kinase, mitochondrial | Dguok | mt DNA synthesis | 0.49 | 0.041 | NC | NS |
| Probable ATP-dependent RNA helicase DHX40 | Dhx40 | Splicing, RNA helicase | -1.17 | 0.000 | 0.91 | 0.000 |
| DNA replication ATP-dependent helicase/nuclease DNA2 | Dna2 | mt DNA replication | -2.75 | 0.000 | 0.97 | 0.000 |
| DnaJ homolog subfamily B member 11 | Dnajb11 | RNA processing | 2.07 | 0.001 | -1.11 | 0.047 |
| Dr1-associated corepressor | Drap1 | Transcription | NA | NS | 0.59 | 0.004 |
| Evolutionarily conserved signaling intermediate in Toll pathway, mitochondrial | Ecsit | Transcription, mt Transcription | -0.76 | 0.015 | NC | NS |
| Elongation factor 1-alpha 1 | Eef1a1 | Translation | 0.70 | 0.000 | -0.20 | 0.005 |
| Elongation factor 1-alpha 2 | Eef1a2 | Translation, tRNA | 0.10 | 0.065 | -0.14 | 0.005 |
| Elongation factor 1-gamma | Eef1g | Translation | 0.41 | 0.000 | NC | NS |
| Elongation factor 2 | Eef2 | Translation | 0.36 | 0.000 | NC | NS |
| EGF-containing fibulin-like extracellular matrix protein 1 | Efemp1 | Transcription | 0.75 | 0.031 | -0.66 | 0.040 |
| Histone-lysine N-methyltransferase EHMT1 | Ehmt1 | Transcription | 0.64 | 0.052 | NC | NS |
| Eukaryotic translation initiation factor 2 subunit 1 | Eif2s1 | Translation initiation | -0.34 | 0.054 | NC | NS |
| Eukaryotic translation initiation factor 2 subunit 3, Y-linked | Eif2s3y | Translation initiation | -0.63 | 0.000 | NC | NS |
| Eukaryotic translation initiation factor 3 subunit A | Eif3a | Translation initiation | -0.52 | 0.032 | NC | NS |
| Eukaryotic translation initiation factor 3 subunit C | Eif3c | Translation initiation | -0.32 | 0.003 | -0.20 | 0.035 |
| Eukaryotic translation initiation factor 3 subunit D | Eif3d | Translation initiation | -0.86 | 0.001 | 0.46 | 0.043 |
| Eukaryotic translation initiation factor 3 subunit E | Eif3e | Translation initiation | 1.63 | 0.000 | -0.29 | 0.059 |
| Eukaryotic translation initiation factor 3 subunit G | Eif3g | Translation initiation | -0.69 | 0.014 | NC | NS |
| Eukaryotic initiation factor 4A-III | Eif4a3 | Splicing, mRNA stabilization, mRNA transport, Translation | -2.96 | 0.000 | NC | NS |
| Eukaryotic translation initiation factor 4E | Eif4e | Transcription, Translation initiation | 0.86 | 0.000 | NC | NS |
| Eukaryotic translation initiation factor 4 gamma 2 | Eif4g2 | Translation initiation | 0.59 | 0.017 | -0.95 | 0.000 |
| Eukaryotic translation initiation factor 4 gamma 3 | Eif4g3 | Translation initiation | 1.67 | 0.002 | NC | NS |
| Eukaryotic translation initiation factor 4H | Eif4h | RNA helicase, Translation initiation | -1.02 | 0.000 | 0.51 | 0.036 |

| | | | | | | |
|---|---|---|---|---|---|---|
| Eukaryotic translation initiation factor 5A-1 | Eif5a | Translation | -0.35 | 0.014 | NC | NS |
| Eukaryotic translation initiation factor 6 | Eif6 | Translation initiation, mt Translation initiation | 1.40 | 0.000 | NC | NS |
| ELAV-like protein 1 | Elavl1 | mRNA stabilization | -2.45 | 0.000 | 1.26 | 0.023 |
| Elongin-B | Elob | Transcription | NC | NS | -0.44 | 0.003 |
| Elongin-C | Eloc | Transcription | -0.33 | 0.009 | NC | NS |
| Endonuclease G, mitochondrial | Endog | mt DNA replication | -0.25 | 0.030 | -0.25 | 0.021 |
| EPM2A-interacting protein 1 | Epm2aip 1 | Transcription | -0.46 | 0.011 | NC | NS |
| Eukaryotic peptide chain release factor subunit 1 | Etf1 | Translation termination, mRNA catabolism | NC | NS | 0.91 | 0.008 |
| RNA-binding protein EWS | Ewsr1 | Transcription | 0.71 | 0.002 | -0.43 | 0.029 |
| Ezrin | Ezr | Transcription | -0.23 | 0.002 | 0.18 | 0.007 |
| Fatty acid-binding protein, adipocyte | Fabp4 | Transcription | 0.67 | 0.001 | -1.13 | 0.000 |
| FAST kinase domain-containing protein 2, mitochondrial | Fastkd2 | mt ribosomal assembly | 4.85 | 0.000 | -5.68 | 0.000 |
| 40S ribosomal protein S30 | Fau | Translation | 0.21 | 0.026 | NC | NS |
| Bis(5'-adenosyl)-triphosphatase | Fhit | Transcription | -0.56 | 0.001 | NC | NS |
| Four and a half LIM domains protein 1 | Fhl1 | Transcription | 1.38 | 0.000 | -0.31 | 0.058 |
| Four and a half LIM domains protein 2 | Fhl2 | Transcription | -0.73 | 0.042 | NC | NS |
| Synaptic functional regulator FMR1 | Fmr1 | Splicing, RNA stabilization, Histone phosphorylation, miRNA translation | 1.09 | 0.025 | NC | NS |
| Fragile X mental retardation syndrome-related protein 1 | Fxr1 | mRNA transport, Translation | 0.36 | 0.049 | NC | NS |
| Ras GTPase-activating protein-binding protein 1 | G3bp1 | RNA helicase, DNA helicase | 0.31 | 0.042 | -0.32 | 0.025 |
| Glycine--tRNA ligase | Gars | tRNA | NC | | 0.35 | 0.023 |
| Glutamyl-tRNA(Gln) amidotransferase subunit C, mitochondrial | Gatc | mt translation | -1.20 | 0.023 | 1.02 | 0.036 |
| Elongation factor G, mitochondrial | Gfm1 | mt translation | -0.34 | 0.000 | 0.26 | 0.002 |
| Ribosome-releasing factor 2, mitochondrial | Gfm2 | mt translation | -0.81 | 0.010 | 0.80 | 0.007 |
| Lactoylglutathione lyase | Glo1 | transcription | -0.26 | 0.027 | 0.20 | 0.064 |
| Guanine nucleotide-binding protein G(s) subunit alpha isoforms Xlas | Gnas | transcription | -0.89 | 0.000 | NC | NS |

| | | | | | | |
|---|---|---|---|---|---|---|
| G-rich sequence factor 1 | Grsf1 | mt mRNA processing, mt tRNA processing | -0.57 | 0.010 | NC | NS |
| Eukaryotic peptide chain release factor GTP-binding subunit ERF3A | Gspt1 | Translation termination | NC | NS | 0.56 | 0.006 |
| General transcription factor 3C polypeptide | Gtf3c1 | rRNA and tRNA transcription | -1.13 | 0.038 | 1.12 | 0.026 |
| GTP-binding protein 1 | Gtpbp1 | mRNA degradation, mRNA stability | 2.24 | 0.000 | NC | NS |
| Core histone macro-H2A.2 | H2afy2 | Transcription, rRNA transcription, Histone | 1.00 | 0.000 | -0.34 | 0.048 |
| Histone H2A.Z | H2afz | Transcription, Histone | 2.28 | 0.000 | -1.21 | 0.000 |
| Histone H2B type 1 | H2bc1 | Histone, DNA organization | 0.28 | 0.011 | NC | NS |
| HBS1-like protein | Hbs1l | Translation | -1.21 | 0.041 | NC | NS |
| Host cell factor 2 | Hcfc2 | Transcription | 2.39 | 0.039 | NC | NS |
| Histidine triad nucleotide-binding protein 1 | Hint1 | Transcription | 0.54 | 0.003 | NC | NS |
| Histidine triad nucleotide-binding protein 3 | Hint3 | DNA organization | NC | NS | -1.65 | 0.003 |
| Histone H1.1 | Hist1h1a | Transcription, Histone | 2.06 | 0.000 | NA | NS |
| Histone H1.5 | Hist1h1b | DNA methylation, Histone, Transcription | 0.36 | 0.019 | NA | NS |
| Histone H1.2 | Hist1h1c | Transcription, Histone | 1.73 | 0.000 | -1.35 | 0.000 |
| Histone H1t | Hist1h1t | Transcription, Histone | 2.82 | 0.000 | -1.92 | 0.000 |
| Histone H2A type 1 | HIST1H2AA | Histone, DNA organization | 0.81 | 0.000 | -0.84 | 0.000 |
| Histone H2A type 1-F | Hist1h2af | Histone, Transcription | 1.05 | 0.000 | -0.68 | 0.000 |
| Histone H2B type 1-A | Hist1h2ba | Histone, Transcription | -0.81 | 0.002 | 0.40 | 0.081 |
| Histone H2B type 1-C/E/G | Hist1h2bc | Histone, Transcription | 0.32 | 0.005 | NC | NS |
| Histone H3.2 | Hist1h3b | Histone, Transcription | 3.33 | 0.003 | 1.64 | 0.092 |
| Histone H4 | Hist1h4b | Histone, Transcription | 0.81 | 0.000 | -0.52 | 0.000 |
| Histone H2A type 2-C | Hist2h2ac | Histone, Transcription | 2.76 | 0.000 | -1.70 | 0.001 |
| High mobility group protein HMG-I/HMG-Y | Hmga1 | mRNA processing, Transcription | -1.52 | 0.000 | NC | NS |
| High mobility group protein B1 | Hmgb1 | Transcription | 0.65 | 0.001 | -0.33 | 0.055 |
| High mobility group protein B2 | Hmgb2 | Transcription | 1.41 | 0.000 | -1.33 | 0.000 |
| Non-histone chromosomal protein HMG-17 | Hmgn2 | Transcription, Histone | NA | NS | -1.74 | 0.001 |
| Heterogeneous nuclear ribonucleoprotein A0 | Hnrnpa0 | mRNA stabilization | 0.90 | 0.001 | -0.91 | 0.001 |

| | | | | | | |
|---|---|---|---|---|---|---|
| Heterogeneous nuclear ribonucleoprotein A1 | Hnrnpa1 | Transcription, mRNA export, Splicing | -0.33 | 0.020 | NA | NA |
| Heterogeneous nuclear ribonucleoproteins A2/B1 | Hnrnpa2 b1 | mRNA packing, mRNA export, mRNA stability | -0.75 | 0.000 | NC | NS |
| Heterogeneous nuclear ribonucleoprotein A3 | Hnrnpa3 | mRNA transport, mRNA processing, splicing | -0.16 | 0.043 | NC | NS |
| Heterogeneous nuclear ribonucleoprotein C | Hnrnpc | Spicing | 0.46 | 0.009 | -0.31 | 0.048 |
| Heterogeneous nuclear ribonucleoprotein D0 | Hnrnpd | Transcription, mRNA stability | 0.99 | 0.004 | -0.62 | 0.041 |
| Heterogeneous nuclear ribonucleoprotein D-like | Hnrnpdl | Transcription | NA | NA | 2.00 | 0.000 |
| Heterogeneous nuclear ribonucleoprotein F | Hnrnpf | mRNA processing, splicing | 1.47 | 0.004 | NC | NS |
| Heterogeneous nuclear ribonucleoprotein H | Hnrnph1 | mRNA processing, splicing | -0.38 | 0.000 | NC | NS |
| Heterogeneous nuclear ribonucleoprotein H2 | Hnrnph2 | mRNA processing | 0.63 | 0.038 | NC | NS |
| Heterogeneous nuclear ribonucleoprotein L | Hnrnpl | mRNA processing, splicing | -0.51 | 0.000 | NC | NS |
| Heterogeneous nuclear ribonucleoprotein M | Hnrnpm | Splicing, Nuclear import | -1.03 | 0.003 | NC | NS |
| Homeodomain-only protein | Hopx | Transcription | -2.56 | 0.001 | 1.75 | 0.009 |
| Heterochromatin protein 1-binding protein 3 | Hp1bp3 | Transcription | 0.79 | 0.003 | NC | NS |
| Histidine-rich glycoprotein | Hrg | Transcription | -0.57 | 0.001 | 0.28 | 0.048 |
| Heat shock cognate 71 kDa protein | Hspa8 | Splicing, RNA processing | 4.57 | 0.000 | -4.00 | 0.001 |
| Heat shock protein 105 kDa | Hsph1 | Transcription | -0.45 | 0.001 | 0.60 | 0.000 |
| Immunoglobulin-binding protein 1 | Igbp1 | Transcription | 0.37 | 0.010 | NC | NS |
| Protein IMPACT | Impact | Translation | 1.13 | 0.000 | -0.75 | 0.002 |
| Importin-9 | Ipo9 | Nuclear import | 2.57 | 0.000 | -1.46 | 0.001 |
| Interferon regulatory factor 2-binding protein 2 | Irf2bp2 | Transcription | 0.39 | 0.053 | -0.34 | 0.020 |
| Inosine triphosphate pyrophosphatase | Itpa | chromosome organization | 0.43 | 0.032 | -0.39 | 0.037 |
| KN motif and ankyrin repeat domain-containing protein 2 | Kank2 | Transcription, mt Transcription | -0.67 | 0.002 | NC | NS |
| Lysine--tRNA ligase | Kars | tRNA | NC | NS | 0.27 | 0.004 |
| Histone acetyltransferase KAT6A | Kat6a | Histone acetylation, Transcription | 6.26 | 0.000 | NC | NS |
| KH domain-containing, RNA-binding, signal transduction-associated protein 1 | Khdrbs1 | mRNA stability, mRNA export | NC | NS | -1.45 | 0.000 |

| | | | | | | |
|---|---|---|---|---|---|---|
| Far upstream element-binding protein 2 | Khsrp | Transcription, mRNA stability, mRNA processing | -0.33 | 0.001 | NC | NS |
| Histone-lysine N-methyltransferase 2A | Kmt2a | Histone acetylation, Transcription | 7.20 | 0.000 | -4.10 | 0.000 |
| Endoribonuclease LACTB2 | Lactb2 | mt mRNA processing | 0.62 | 0.003 | NC | NS |
| Leucine--tRNA ligase, cytoplasmic | Lars | tRNA | NC | NS | -0.72 | 0.000 |
| Protein LBH | Lbh | Transcription | 0.30 | 0.058 | 0.25 | 0.09 |
| Galectin-3 | Lgals3 | Splicing, mRNA processing | 1.91 | 0.000 | -0.82 | 0.072 |
| LIM domain-containing protein 1 | Limd1 | miRNA gene silencing, Transcription, mRNA processing | NC | NS | -1.85 | 0.001 |
| Hormone-sensitive lipase | Lipe | Transcription | 0.75 | 0.008 | 1.15 | 0.002 |
| LIM and cysteine-rich domains protein 1 | Lmcd1 | Transcription | 0.46 | 0.003 | NC | NS |
| Lon protease homolog, mitochondrial | Lonp1 | mt gene expression/transcription | NC | NS | 0.28 | 0.003 |
| Phosphatidate phosphatase LPIN1 | Lpin1 | Transcription, Histone deacetylation, mt Transcription | 0.54 | 0.000 | -0.42 | 0.000 |
| Leucine-rich PPR motif-containing protein, mitochondrial | Lrpprc | mRNA export, Transcription, mt Transcription, mRNA stability | -0.32 | 0.000 | 0.33 | 0.000 |
| Leucine-rich repeat flightless-interacting protein 1 | Lrrfip1 | Transcription | 0.70 | 0.032 | NC | NS |
| U6 snRNA-associated Sm-like protein LSm2 | Lsm2 | Splicing | NC | NS | 2.34 | 0.000 |
| Lumican | Lum | Transcription | 0.35 | 0.005 | NC | NS |
| O-acetyl-ADP-ribose deacetylase MACROD1 | Macrod1 | Histone deacetylation | -0.54 | 0.000 | NC | NS |
| Meiosis arrest female protein 1 | Marf1 | RNA metabolic process | -0.61 | 0.034 | NC | NS |
| Matrin-3 | Matr3 | Transcription, retention of defective RNAs | -1.60 | 0.000 | NC | NS |
| Mitochondrial antiviral-signaling protein | Mavs | Transcription, Nuclear import, mt Transcription | -0.288 | 0.098 | -0.4 | 0.018 |
| Melanocyte-stimulating hormone receptor | Mc1r | Transcription | 3.08 | 0.000 | NC | NS |
| Methyl-CpG-binding protein 2 | Mecp2 | Transcription, Histone deacetylation, DNA methylation | NC | NS | 0.71 | 0.051 |
| Menin | Men1 | Histone methylation, Transcription | 0.66 | 0.057 | NC | NS |
| Muscular LMNA-interacting protein | Mlip | Transcription | -0.31 | 0.07 | 0.85 | 0.000 |

| | | | | | | |
|---|---|---|---|---|---|---|
| 39S ribosomal protein L1, mitochondrial | Mrpl1 | mt Translation, mt rRNA processing | 0.82 | 0.016 | NC | NS |
| 39S ribosomal protein L12, mitochondrial | Mrpl12 | mt Transcription, mt Translation | -0.39 | 0.001 | 0.20 | 0.034 |
| 39S ribosomal protein L23, mitochondrial | Mrpl23 | mt Translation | -1.30 | 0.001 | 0.63 | 0.068 |
| 39S ribosomal protein L28, mitochondrial | Mrpl28 | mt Translation | NC | NS | 3.59 | 0.001 |
| 39S ribosomal protein L38, mitochondrial | Mrpl38 | mt Translation | 0.96 | 0.021 | NC | NS |
| 39S ribosomal protein L4, mitochondrial | Mrpl4 | mt Translation, Translation | 3.11 | 0.000 | NC | NS |
| 39S ribosomal protein L40, mitochondrial | Mrpl40 | mt Translation | NC | NS | 0.39 | 0.001 |
| 39S ribosomal protein L41, mitochondrial | Mrpl41 | mt Translation | -1.21 | 0.000 | -0.41 | 0.076 |
| 39S ribosomal protein L46, mitochondrial | Mrpl46 | mt Translation | -0.59 | 0.026 | NC | NS |
| 39S ribosomal protein L49, mitochondrial | Mrpl49 | mt Translation | -0.55 | 0.000 | 0.24 | 0.003 |
| Peptidyl-tRNA hydrolase ICT1, mitochondrial | Mrpl58 | mt Translation | 2.46 | 0.000 | 0.43 | 0.088 |
| 28S ribosomal protein S14, mitochondrial | Mrps14 | mt Translation | -2.08 | 0.000 | 1.08 | 0.005 |
| 28S ribosomal protein S15, mitochondrial | Mrps15 | mt Translation | -0.99 | 0.086 | 1.07 | 0.047 |
| 28S ribosomal protein S16, mitochondrial | Mrps16 | mt Translation | -0.87 | 0.052 | NC | NS |
| 28S ribosomal protein S22, mitochondrial | Mrps22 | mt Translation | -1.70 | 0.000 | 0.49 | 0.088 |
| 28S ribosomal protein S23, mitochondrial | Mrps23 | mt Translation | -1.35 | 0.003 | 0.95 | 0.002 |
| 28S ribosomal protein S27, mitochondrial | Mrps27 | mt Translation | -3.26 | 0.001 | 3.13 | 0.001 |
| 28S ribosomal protein S5, mitochondrial | Mrps5 | mt Translation | -0.64 | 0.000 | 0.42 | 0.005 |
| 28S ribosomal protein S6, mitochondrial | Mrps6 | mt Translation | -0.90 | 0.006 | NC | NS |
| 28S ribosomal protein S7, mitochondrial | Mrps7 | mt Translation | -0.39 | 0.047 | NC | NS |
| Ribosome-recycling factor, mitochondrial | Mrrf | mt Translation termination | -0.44 | 0.026 | NC | NS |
| RNA-binding protein Musashi homolog 2 | Msi2 | mRNA processing | 0.38 | 0.019 | NC | NS |
| Methionyl-tRNA formyltransferase, mitochondrial | Mtfmt | mt tRNA | 0.81 | 0.000 | 0.52 | 0.003 |

| | | | | | | |
|---|---|---|---|---|---|---|
| Myotrophin | Mtpn | Transcription | 1.93 | 0.000 | NC | NS |
| Muscle-related coiled-coil protein | Murc | Transcription | 0.43 | 0.000 | NC | NS |
| N-alpha-acetyltransferase 10 | Naa10 | Transcription | 1.20 | 0.012 | 0.96 | 0.040 |
| N-alpha-acetyltransferase 15, NatA auxiliary subunit | Naa15 | Transcription | -0.79 | 0.008 | NC | NS |
| Nucleolin | Ncl | rRNA transcription, Transcriptional elongation | -0.35 | 0.000 | 0.22 | 0.008 |
| Nuclear receptor corepressor 2 | Ncor2 | miRNA, Transcription, Histone deacetylation | -1.80 | 0.000 | NC | NS |
| NF-kappa-B inhibitor alpha | Nfkbia | Nuclear import, Transcription | NC | NS | 1.18 | 0.000 |
| NKAP-like protein | Nkapl | Transcription | -1.03 | 0.059 | 1.17 | 0.012 |
| Protein NLRC3 | Nlrc3 | Transcription | 0.32 | 0.007 | NC | NS |
| Non-POU domain-containing octamer-binding protein | Nono | Transcription | -0.94 | 0.004 | NC | NS |
| Neuronal PAS domain-containing protein 4 | Npas4 | Transcription | -0.50 | 0.049 | 0.68 | 0.006 |
| Nucleophosmin | Npm1 | rRNA transcription, rRNA export | 0.40 | 0.001 | NC | NS |
| Cleavage and polyadenylation specificity factor subunit 5 | Nudt21 | mRNA processing | -0.86 | 0.000 | -0.56 | 0.000 |
| Nuclear pore complex protein Nup85 | Nup85 | mRNA export, Transcription | 0.63 | 0.005 | 0.40 | 0.035 |
| UDP-N-acetylglucosamine--peptide N-acetylglucosaminyltransferase 110 kDa subunit | Ogt | Histone acetylation, Transcription | -0.69 | 0.016 | 1.31 | 0.000 |
| Polyadenylate-binding protein 1 | Pabpc1 | Splicing, mRNA processing, mRNA catabolic process | 0.67 | 0.000 | -0.32 | 0.009 |
| Polyadenylate-binding protein-interacting protein 1 | Paip1 | Translation initiation | 1.41 | 0.000 | NC | NS |
| Protein DJ-1 | Park7 | Transcription | 0.47 | 0.035 | NC | NS |
| Pre-B-cell leukemia transcription factor-interacting protein 1 | Pbxip1 | Transcription | -0.59 | 0.018 | NC | NS |
| Pterin-4-alpha-carbinolamine dehydratase | Pcbd1 | Transcription | -0.75 | 0.000 | 0.32 | 0.015 |
| Pterin-4-alpha-carbinolamine dehydratase 2 | Pcbd2 | Transcription | -21.33 | 0.000 | 21.33 | 0.000 |
| Poly(rC)-binding protein 1 | Pcbp1 | mRNA processing, Transcription | 0.42 | 0.001 | NC | NS |
| Poly(rC)-binding protein 3 | Pcbp3 | Transcription | 0.32 | 0.011 | NC | NS |
| PDZ and LIM domain protein 1 | Pdlim1 | Transcription | 0.84 | 0.000 | -0.20 | 0.000 |
| Protein pelota homolog | Pelo | mRNA catabolic process | 0.44 | 0.059 | -0.41 | 0.055 |

| | | | | | | |
|---|---|---|---|---|---|---|
| Prohibitin | Phb | Transcription, mt Transcription | -0.66 | 0.006 | NC | NS |
| Prohibitin-2 | Phb2 | Transcription, mt Transcription | -0.84 | 0.001 | NC | NS |
| Peptidyl-prolyl cis-trans isomerase NIMA-interacting 1 | Pin1 | Transcription | 0.53 | 0.007 | NC | NS |
| Pirin | Pir | Transcription | 0.81 | 0.000 | -0.44 | 0.014 |
| cAMP-dependent protein kinase inhibitor alpha | Pkia | Transcription | 2.94 | 0.000 | -1.05 | 0.005 |
| Polymerase delta-interacting protein 2 | Poldip2 | Transcription | -0.39 | 0.045 | NC | NS |
| Peptidyl-prolyl cis-trans isomerase E | Ppie | Splicing, RNA processing, Transcription | -0.45 | 0.003 | 0.45 | 0.002 |
| Protein phosphatase 1A | Ppm1a | Transcription | 0.37 | 0.042 | -0.30 | 0.069 |
| Serine/threonine-protein phosphatase PP1-alpha catalytic subunit | Ppp1ca | Translation | -0.82 | 0.000 | 0.58 | 0.002 |
| Protein phosphatase 1 regulatory subunit 12A | Ppp1r12a | Transcription | 0.53 | 0.000 | -0.41 | 0.002 |
| Serine/threonine-protein phosphatase 2B catalytic subunit alpha isoform | Ppp3ca | miRNA, Transcription | -0.55 | 0.000 | 0.52 | 0.000 |
| Peroxisome proliferator-activated receptor gamma coactivator-related protein 1 | Pprc1 | Transcription | -1.68 | 0.005 | 1.01 | .0.096 |
| Major prion protein | Prnp | Transcription | -1.10 | 0.009 | NC | NS |
| Protein PRRC2A | Prrc2a | Splicing | 2.77 | 0.000 | -1.10 | 0.037 |
| PC4 and SFRS1-interacting protein | Psip1 | Transcription, Splicing | 1.90 | 0.000 | -0.36 | 0.003 |
| 26S protease regulatory subunit 4 | Psmc1 | Transcription | 0.35 | 0.032 | -0.27 | 0.07 |
| 26S protease regulatory subunit 10B | Psmc6 | Transcription | 0.71 | 0.000 | NC | NS |
| 26S proteasome non-ATPase regulatory subunit 9 | Psmd9 | Transcription | 0.36 | 0.053 | NC | NS |
| Polypyrimidine tract-binding protein 1 | Ptbp1 | Splicing | NC | NS | 0.28 | 0.023 |
| Polypyrimidine tract-binding protein 2 | Ptbp2 | Splicing | -0.46 | 0.034 | 0.78 | 0.000 |
| Pentatricopeptide repeat domain-containing protein 3, mitochondrial | Ptcd3 | mt Translation | -0.66 | 0.002 | 0.62 | 0.003 |
| Prostaglandin E synthase 2 | Ptges2 | Transcription | -0.49 | 0.006 | NC | NS |
| Glutamyl-tRNA(Gln) amidotransferase subunit A, mitochondrial | Qrsl1 | tRNA, mt Translation | -0.40 | 0.034 | NC | NS |

| | | | | | | |
|---|---|---|---|---|---|---|
| RNA-binding protein Raly | Raly | Transcription, Splicing, RNA processing | NC | NS | 26.67 | 0.000 |
| Arginine--tRNA ligase, cytoplasmic | Rars | tRNA | NC | NS | 0.26 | 0.003 |
| RNA binding motif protein, X-linked-like-1 | Rbmxl1 | mRNA processing, splicing | -0.54 | 0.008 | NC | NS |
| Oligoribonuclease, mitochondrial | Rexo2 | Nucleotide recycling | 0.57 | 0.026 | NC | NS |
| DNA-binding protein RFX2 | Rfx2 | Transcription | -1.29 | 0.011 | NC | NS |
| Transforming protein RhoA | Rhoa | Transcription | 0.51 | 0.004 | NC | NS |
| Rho-related GTP-binding protein RhoG | Rhog | Transcription | -0.99 | 0.014 | NC | NS |
| 60S ribosomal protein L10-like | Rpl10l | Translation | 0.76 | 0.001 | -0.38 | 0.051 |
| 60S ribosomal protein L11 | Rpl11 | Translation | 0.65 | 0.000 | NC | NS |
| 60S ribosomal protein L12 | Rpl12 | Translation | 0.43 | 0.001 | NC | NS |
| 60S ribosomal protein L13 | Rpl13 | Translation | 0.35 | 0.000 | NC | NS |
| 60S ribosomal protein L14 | Rpl14 | Translation | 1.06 | 0.000 | NC | NS |
| 60S ribosomal protein L15 | Rpl15 | Translation | -0.47 | 0.008 | NC | NS |
| 60S ribosomal protein L17 | Rpl17 | Translation | 1.24 | 0.000 | NC | NS |
| 60S ribosomal protein L18 | Rpl18 | Translation | -0.32 | 0.009 | NC | NS |
| 60S ribosomal protein L18a | Rpl18a | Translation | NC | NS | -1.10 | 0.002 |
| 60S ribosomal protein L19 | Rpl19 | Translation | -1.19 | 0.000 | NC | NS |
| 60S ribosomal protein L21 | Rpl21 | Translation | 0.78 | 0.000 | NC | NS |
| 60S ribosomal protein L22 | Rpl22 | Translation | 0.62 | 0.026 | NC | NS |
| 60S ribosomal protein L23 | Rpl23 | Translation | 0.43 | 0.002 | -0.48 | 0.000 |
| 60S ribosomal protein L23a | Rpl23a | Translation | 0.41 | 0.005 | -0.39 | 0.004 |
| 60S ribosomal protein L24 | Rpl24 | Translation | 0.74 | 0.000 | NC | NS |
| 60S ribosomal protein L26 | Rpl26 | Translation | 0.83 | 0.000 | NC | NS |
| 60S ribosomal protein L27 | Rpl27 | Translation | 0.64 | 0.005 | NC | NS |
| 60S ribosomal protein L27a | Rpl27a | Translation | 0.35 | 0.011 | NC | NS |
| 60S ribosomal protein L28 | Rpl28 | Translation | 0.86 | 0.000 | -0.36 | 0.003 |
| 60S ribosomal protein L29 | Rpl29 | Translation | 0.36 | 0.004 | -0.25 | 0.027 |
| 60S ribosomal protein L3 | Rpl3 | Translation | 0.83 | 0.006 | -0.57 | 0.033 |
| 60S ribosomal protein L30 | Rpl30 | Translation | 0.71 | 0.000 | NC | NS |
| 60S ribosomal protein L31 | Rpl31 | Translation | 0.41 | 0.002 | NC | NS |
| 60S ribosomal protein L32 | Rpl32 | Translation | 1.24 | 0.001 | -0.99 | 0.002 |
| 60S ribosomal protein L34 | Rpl34 | Translation | 1.02 | 0.003 | -0.84 | 0.007 |
| 60S ribosomal protein L35 | Rpl35 | Translation | 0.87 | 0.000 | NC | NS |

| | | | | | | |
|---|---|---|---|---|---|---|
| 60S ribosomal protein L35a | Rpl35a | Translation | -0.76 | 0.004 | NC | NS |
| 60S ribosomal protein L36a | Rpl36a | Translation | 1.34 | 0.000 | NC | NS |
| 60S ribosomal protein L37a | Rpl37a | Translation | 1.20 | 0.000 | -0.81 | 0.001 |
| Putative 60S ribosomal protein L37a | Rpl37a-ps1 | Translation | 0.92 | 0.012 | -1.42 | 0.000 |
| 60S ribosomal protein L38 | Rpl38 | Translation | 0.57 | 0.013 | -0.67 | 0.002 |
| 60S ribosomal protein L39 | Rpl39 | Translation | 0.69 | 0.000 | -0.25 | 0.061 |
| 60S ribosomal protein L4 | Rpl4 | Translation | -0.47 | 0.049 | NC | NS |
| 60S ribosomal protein L6 | Rpl6 | Translation | 0.82 | 0.000 | NC | NS |
| 60S ribosomal protein L7 | Rpl7 | Translation | -0.56 | 0.004 | NC | NS |
| 60S ribosomal protein L8 | Rpl8 | Translation | NC | NS | -0.29 | 0.057 |
| 60S ribosomal protein L9 | Rpl9 | Translation | 0.40 | 0.005 | -0.40 | 0.003 |
| 60S acidic ribosomal protein P1 | Rplp1 | Translation elongation | NC | NS | -0.26 | 0.040 |
| 40S ribosomal protein S10 | Rps10 | Translation | NC | NS | 0.38 | 0.023 |
| 40S ribosomal protein S12 | Rps12 | Translation | 0.81 | 0.021 | NC | NS |
| 40S ribosomal protein S15 | Rps15 | Translation | NC | NS | -1.05 | 0.001 |
| 40S ribosomal protein S17 | Rps17 | Translation | NC | NS | 0.17 | 0.030 |
| 40S ribosomal protein S2 | Rps2 | Translation | -0.31 | 0.008 | NC | NS |
| 40S ribosomal protein S20 | Rps20 | Translation | -0.43 | 0.001 | 0.37 | 0.002 |
| 40S ribosomal protein S21 | Rps21 | Translation | -3.15 | 0.000 | 2.36 | 0.000 |
| 40S ribosomal protein S23 | Rps23 | Translation | 0.98 | 0.007 | NC | NS |
| 40S ribosomal protein S24 | Rps24 | Translation initiation | 0.30 | 0.001 | NC | NS |
| 40S ribosomal protein S26 | Rps26 | Translation | 0.49 | 0.000 | -0.26 | 0.030 |
| 40S ribosomal protein S27 | Rps27 S27-1 | Translation | 0.41 | 0.016 | -0.29 | 0.059 |
| Ubiquitin-40S ribosomal protein S27a | Rps27a | Translation | 1.20 | 0.047 | -0.9 | 0.066 |
| 40S ribosomal protein S28 | Rps28 | Translation | -1.07 | 0.021 | 1.70 | 0.000 |
| 40S ribosomal protein S29 | Rps29 | Translation | -0.50 | 0.052 | NA | NS |
| 40S ribosomal protein S3a | Rps3a | Translation | 0.26 | 0.022 | NA | NS |
| 40S ribosomal protein S4, X isoform | Rps4x | Translation | NC | NS | -0.23 | 0.053 |
| 40S ribosomal protein S5 | Rps5 | Translation | 0.67 | 0.000 | NC | NS |
| 40S ribosomal protein S6 | Rps6 | Translation, rRNA processing | 0.99 | 0.000 | NC | NS |
| Ribosomal protein S6 kinase alpha-3 | Rps6ka3 | Transcription | NC | NS | -0.92 | 0.002 |
| Ribosomal protein S6 kinase alpha-6 | Rps6ka6 | Transcription | 1.53 | 0.003 | NC | NS |

| | | | | | | |
|---|---|---|---|---|---|---|
| 40S ribosomal protein S7 | Rps7 | Translation | 0.69 | 0.001 | NC | NS |
| 40S ribosomal protein S8 | Rps8 | Translation, rRNA transcription | NC | NS | -0.35 | 0.002 |
| 40S ribosomal protein S9 | Rps9 | Translation fidelity | -0.93 | 0.000 | NC | NS |
| 40S ribosomal protein SA | Rpsa | rRNA export, Translation | -0.27 | 0.002 | 0.19 | 0.018 |
| Runt-related transcription factor 1 | Runx1 | Transcription | -1.35 | 0.003 | 1.09 | 0.007 |
| RuvB-like 2 | Ruvbl2 | Transcription, Histone acetylation | -1.33 | 0.000 | -1.19 | 0.000 |
| RWD domain-containing protein 1 | Rwdd1 | Translation | NC | NS | 0.38 | 0.006 |
| Protein S100-A1 | S100a1 | Transcription | -1.54 | 0.094 | 1.74 | 0.056 |
| Scaffold attachment factor B1 | Safb | Transcription, mRNA processing | 0.27 | 0.008 | 0.43 | 0.000 |
| Scaffold attachment factor B2 | Safb2 | Transcription, mRNA processing | -0.65 | 0.031 | NC | NS |
| SAP domain-containing ribonucleoprotein | Sarnp | mRNA export, Transcription, Translation | NC | NS | -0.74 | 0.048 |
| Serine--tRNA ligase, mitochondrial | Sars2 | tRNA, mt tRNA | -0.98 | 0.001 | 0.93 | 0.001 |
| Protein SEC13 homolog | Sec13 | mRNA transport | 1.15 | 0.000 | NC | NS |
| Selenocysteine insertion sequence-binding protein 2 | Secisbp2 | Translation initiation | -2.41 | 0.001 | -1.15 | 0.095 |
| Sentrin-specific protease 2 | Senp2 | Transcription | NC | NS | 0.27 | 0.037 |
| Protein SET | Set | Transcription | 0.75 | 0.010 | NC | NS |
| Splicing factor 1 | Sf1 | Splicing, Transcription | -3.04 | 0.008 | 4.80 | 0.001 |
| Splicing factor, proline- and glutamine-rich | Sfpq | Histone deacetylation, Transcription | -0.54 | 0.016 | -0.37 | 0.066 |
| Swi5-dependent recombination DNA repair protein 1 homolog | Sfr1 | Transcription | 1.31 | 0.000 | NC | NS |
| NAD-dependent protein deacetylase sirtuin-2 | Sirt2 | Histone deacetylation, Transcription | -0.48 | 0.004 | NC | NS |
| Zinc transporter 9 | Slc30a9 | Transcription, Wnt | -0.69 | 0.005 | 0.45 | 0.058 |
| Staphylococcal nuclease domain-containing protein 1 | Snd1 | Transcription | 0.73 | 0.012 | NC | NS |
| Small nuclear ribonucleoprotein Sm D2 | Snrpd2 | Splicing | -1.24 | 0.002 | 1.18 | 0.002 |
| Small nuclear ribonucleoprotein E | Snrpe | Splicing | -0.40 | 0.008 | -0.70 | 0.000 |
| Small nuclear ribonucleoprotein F | Snrpf | Splicing | 1.98 | 0.000 | -1.66 | 0.000 |
| Sorting nexin-6 | Snx6 | Transcription | -0.31 | 0.005 | 0.37 | 0.001 |
| Sequestosome-1 | Sqstm1 | Transcription | 0.44 | 0.048 | NC | NS |

| | | | | | | |
|---|---|---|---|---|---|---|
| Sterol regulatory element-binding protein 1 | Srebf1 | Transcription | -0.90 | 0.054 | 1.25 | 0.007 |
| Serine/arginine-rich splicing factor 1 | Srsf1 | Splicing, mRNA transport | -1.06 | 0.000 | NC | NS |
| Serine/arginine-rich splicing factor 3 | Srsf3 | Splicing, mRNA transport | -0.93 | 0.001 | NC | NS |
| Single-stranded DNA-binding protein, mitochondrial | Ssbp1 | mt DNA replication | NC | NS | 0.47 | 0.001 |
| Signal transducer and activator of transcription 3 | Stat3 | Transcription | 0.71 | 0.000 | NC | NS |
| Signal transducer and activator of transcription 5B | Stat5b | Transcription | -1.30 | 0.028 | 1.54 | 0.007 |
| Striatin-3 | Strn3 | Transcription | -2.17 | 0.021 | NC | NS |
| Activated RNA polymerase II transcriptional coactivator p15 | Sub1 | Transcription | -1.03 | 0.014 | 1.24 | 0.002 |
| ATP-dependent RNA helicase SUPV3L1, mitochondrial | Supv3l1 | mt MRNA catabolic, mt RNA helicase | -0.90 | 0.008 | NC | NS |
| Translational activator of cytochrome c oxidase 1 | Taco1 | mt translation | NC | NS | 0.43 | 0.005 |
| TAR DNA-binding protein 43 | Tardbp | Splicing, Transcription, mRNA stability, mRNA processing | -0.41 | 0.001 | NC | NS |
| Threonine--tRNA ligase, cytoplasmic | Tars | tRNA | 1.23 | 0.000 | NC | NS |
| Calcineurin B homologous protein 3 | Tesc | Transcription | 0.31 | 0.009 | NC | NS |
| Transcription factor A, mitochondrial | Tfam | mt Transcription | 0.61 | **0.007** | -0.36 | 0.076 |
| Lamina-associated polypeptide 2, isoforms alpha/zeta | Tmpo | Transcription | NC | NS | -0.75 | 0.001 |
| DNA topoisomerase 2-alpha | Top2a | Transcription | -2.28 | 0.037 | NC | NS |
| DNA topoisomerase 2-beta | Top2b | Mitosis | 1.82 | 0.004 | NC | NS |
| Translationally-controlled tumor protein | Tpt1 | Transcription factor binding | 0.26 | 0.043 | NC | NS |
| Transformer-2 protein homolog beta | Tra2b | Splicing | -0.96 | 0.023 | NC | NS |
| E3 ubiquitin/ISG15 ligase TRIM25 | Trim25 | Transcription factor activity | -1.08 | 0.004 | NC | NS |
| Transcription intermediary factor 1-beta | Trim28 | Transcription factor activity | -1.04 | 0.014 | 0.69 | 0.068 |
| Tripartite motif-containing protein 55 | Trim55 | Gene expression | -0.86 | 0.028 | NC | NS |
| TSC22 domain family protein 1 | Tsc22d1 | Transcription factor activity | 1.57 | 0.000 | NC | NS |
| Elongation factor Ts, mitochondrial | Tsfm | mt Translational elongation | NC | NS | 0.43 | 0.001 |

| | | | | | | |
|---|---|---|---|---|---|---|
| Translin | Tsn | RNA-induced silencing | 0.61 | 0.003 | NC | NS |
| Thiosulfate sulfurtransferase | Tst | rRNA transport, mt import of rRNA | -0.76 | 0.000 | 0.34 | 0.003 |
| Thioredoxin | Txn | nuclear import, Transcription | 0.95 | 0.000 | -0.80 | 0.000 |
| Splicing factor U2AF 35 kDa subunit | U2af1 | Splicing | -0.88 | 0.016 | -0.56 | 0.09 |
| NEDD8-activating enzyme E1 catalytic subunit | Uba3 | Transcription | -1.64 | 0.005 | 1.28 | 0.015 |
| Regulator of nonsense transcripts 1 | Upf1 | mRNA catabolic process, Translation termination | -0.65 | 0.003 | 0.37 | 0.054 |
| Ubiquinol-cytochrome-c reductase complex assembly factor 2 | Uqcc2 | mt translation | -1.22 | 0.005 | 1.57 | 0.001 |
| Ubiquitin carboxyl-terminal hydrolase 47 Isoform 2 | Usp47 | Transcription | NC | NS | 1.65 | 0.000 |
| Ubiquitin carboxyl-terminal hydrolase 47 | Usp47 | Transcription | -0.67 | 0.001 | 0.88 | 0.000 |
| Valine--tRNA ligase | Vars | mt tRNA | NC | NS | 0.27 | 0.002 |
| Vascular endothelial growth factor B | Vegfb | negative regulation of gene expression | -1.25 | 0.015 | NC | NS |
| Vacuolar protein-sorting-associated protein 36 | Vps36 | Transcription | NC | NS | -0.71 | 0.008 |
| Tryptophan--tRNA ligase, cytoplasmic | Wars | tRNA | 2.48 | 0.000 | NC | NS |
| Neural Wiskott-Aldrich syndrome protein | Wasl | Transcription | 2.11 | 0.000 | -1.40 | 0.005 |
| WD repeat-containing protein 62 | Wdr62 | Splicing | 2.50 | 0.000 | NC | NS |
| Wolframin | Wfs1 | Transcription | -1.06 | 0.010 | -1.03 | 0.007 |
| WW domain-containing transcription regulator protein 1 | Wwtr1 | Transcription | -2.06 | 0.016 | NC | NS |
| Exportin-1 | Xpo1 | mRNA transport, Transcription, nuclear export, rRNA export | NC | NS | 0.33 | 0.043 |
| Transcriptional coactivator YAP1 | Yap1 | Transcription | 1.13 | 0.003 | -0.31 | 0.083 |
| Tyrosine--tRNA ligase, cytoplasmic | Yars | tRNA | 0.55 | 0.002 | NC | NS |
| Y-box-binding protein 2 | Ybx2 | mRNA stabilization, Translation, Transcription | 0.35 | 0.014 | -0.23 | 0.071 |
| Y-box-binding protein 3 | Ybx3 | Transcription | 0.65 | 0.031 | -0.47 | 0.089 |
| Y-box-binding protein 3 Isoform 2 | Ybx3 | Transcription | NA | NS | -1.20 | 0.000 |
| 14-3-3 protein epsilon | Ywhae | Nuclear export | 0.18 | 0.03 | NC | NS |
| 14-3-3 protein eta | Ywhah | Transcription | 0.33 | 0.000 | NC | NS |

| | | | | | | |
|---|---|---|---|---|---|---|
| 14-3-3 protein theta | Ywhaq | Transcription | 0.54 | 0.000 | -0.16 | 0.004 |
| 14-3-3 protein zeta/delta | Ywhaz | Transcription factor binding | -0.30 | 0.006 | NC | NS |
| Telomere zinc finger-associated protein Isoform 2 | Zbtb48 | Transcription | -1.23 | 0.000 | NC | NS |
| Zinc finger protein 12 | Znf12 | Transcription | -1.21 | 0.000 | 1.23 | 0.000 |
| UPF0568 protein C14orf166 homolog | | tRNA ligation | 0.69 | 0.000 | NC | NS |

NA = no appropriate NC = no change NS = not significant

**Supplemental table 2**: List of MS data showing significant changed protein concentrations in human HFpEF samples verses control in comparison to significant changed protein in rat HFpEF verses control LV. Shown are the foldchange values (log2) of HFpEF verses control for human and rat samples for total protein concentration if significant ($p<0.05$; n(human)=7-9; n(rat)=3-4).

| Protein name | Gene name | Foldchange (log2) human HFpEF vs control | p-value human HFpEF vs control | Foldchange (log2) rat HFpEF vs control | p-value human HFpEF vs control |
|---|---|---|---|---|---|
| 14-3-3 protein epsilon | 1433E | -0.38 | 0.000 | 0.18 | 0.026 |
| 14-3-3 protein eta | 1433F | -0.33 | 0.009 | 0.33 | 0.000 |
| 14-3-3 protein theta | 1433T | -0.40 | 0.000 | 0.26 | 0.038 |
| Alpha-1-acid glycoprotein 1 | A1AG1 | 0.57 | 0.000 | 1.50 | 0.000 |
| Alpha-1-acid glycoprotein 2 | A1AG2 | 0.31 | 0.000 | NC | NS |
| Alpha-aminoadipic semialdehyde synthase | AASS | -0.47 | 0.000 | NC | NS |
| Actin-binding LIM protein 1 | ABLM1 | -0.32 | 0.000 | 0.46 | 0.002 |
| Short-chain specific acyl-CoA dehydrogenase | ACADS | -0.46 | 0.000 | -0.36 | 0.001 |
| Acyl-CoA synthetase family member 2 | ACSF2 | -0.37 | 0.009 | -0.54 | 0.000 |
| Alpha-actinin-2 | ACTN2 | -0.30 | 0.000 | 0.61 | 0.000 |
| ADP/ATP translocase 1 | ADT1 | -0.40 | 0.000 | -0.85 | 0.001 |
| 2-aminoethanethiol dioxygenase | AEDO | -0.62 | 0.001 | NC | NS |
| Fatty aldehyde dehydrogenase | AL3A2 | 0.33 | 0.006 | -0.96 | 0.016 |
| Alanine aminotransferase 1 | ALAT1 | -0.32 | 0.000 | -0.55 | 0.001 |
| Serum albumin | ALBU | 0.36 | 0.000 | NC | NS |
| Aldehyde dehydrogenase | ALDH2 | -0.24 | 0.000 | 1.19 | 0.000 |
| Fructose-bisphosphate aldolase C | ALDOC | -0.28 | 0.000 | NC | NS |
| Protein arginine N-methyltransferase 1 | ANM1 | -0.58 | 0.000 | NC | NS |
| Protein archease | ARCH | -0.51 | 0.009 | NC | NS |
| Aflatoxin B1 aldehyde reductase member 2 | ARK72 | -0.21 | 0.046 | 0.39 | 0.019 |
| ADP-ribosylation factor-like protein 8B | ARL8B | -0.71 | 0.036 | -1.41 | 0.000 |
| Arylsulfatase B | ARSB | -0.45 | 0.000 | NC | NS |
| Acid ceramidase | ASAH1 | -0.31 | 0.000 | 0.47 | 0.047 |
| ATPase ASNA1 | ASNA | -0.58 | 0.001 | NC | NS |
| Sodium/potassium-transporting ATPase subunit beta-1 | AT1B1 | -0.31 | 0.000 | -0.84 | 0.000 |
| ATP synthase F(0) complex subunit B1 | AT5F1 | -0.44 | 0.000 | -0.95 | 0.000 |
| ATP synthase subunit d | ATP5H | -0.23 | 0.000 | -0.57 | 0.000 |
| ATP synthase subunit alpha | ATPA | -0.49 | 0.000 | NC | NS |
| ATP synthase subunit beta | ATPB | -0.39 | 0.000 | -1.17 | 0.000 |
| ATP synthase subunit delta | ATPD | -0.44 | 0.000 | NC | NS |
| ATP synthase subunit gamma | ATPG | -0.48 | 0.000 | 0.22 | 0.009 |
| ATP synthase subunit O | ATPO | -0.41 | 0.000 | NC | NS |
| Barrier-to-autointegration factor | BAF | -0.36 | 0.005 | 1.82 | 0.000 |
| Branched-chain-amino-acid aminotransferase | BCAT2 | -0.45 | 0.000 | NC | NS |

| | | | | | |
|---|---|---|---|---|---|
| [3-methyl-2-oxobutanoate dehydrogenase [lipoamide]] kinase | BCKD | -0.36 | 0.000 | NC | NS |
| 3-hydroxybutyrate dehydrogenase type 2 | BDH2 | -0.43 | 0.000 | NC | NS |
| Valacyclovir hydrolase | BPHL | -0.42 | 0.016 | -0.60 | 0.021 |
| Basic leucine zipper and W2 domain-containing protein 2 | BZW2 | -0.46 | 0.000 | -0.45 | 0.008 |
| Succinate dehydrogenase cytochrome b560 subunit | C560 | -0.52 | 0.008 | NC | NS |
| Putative transferase CAF17 | CAF17 | 0.40 | 0.013 | NC | NS |
| Carbonic anhydrase 1 | CAH1 | 0.51 | 0.000 | -1.07 | 0.000 |
| Carbonic anhydrase 4 | CAH4 | -0.54 | 0.000 | -1.05 | 0.006 |
| Cathepsin D | CATD | -0.47 | 0.000 | NC | NS |
| Cathepsin Z | CATZ | -0.42 | 0.000 | NC | NS |
| Caveolae-associated protein 1 | CAVN1 | -0.18 | 0.000 | NC | NS |
| Mast cell carboxypeptidase A | CBPA3 | -0.52 | 0.000 | -8.74 | 0.000 |
| Platelet glycoprotein 4 | CD36 | -0.37 | 0.000 | -0.25 | 0.058 |
| CD63 antigen | CD63 | -0.65 | 0.000 | NC | NS |
| CD9 antigen | CD9 | -0.57 | 0.043 | -1.32 | 0.001 |
| Cell division control protein 42 homolog | CDC42 | -0.49 | 0.000 | 0.45 | 0.008 |
| Complement factor H | CFAH | 0.49 | 0.000 | NC | NS |
| Coiled-coil-helix-coiled-coil-helix domain-containing protein 1 | CHCH1 | -0.46 | 0.001 | NC | NS |
| Coiled-coil-helix-coiled-coil-helix domain-containing protein 2 | CHCH2 | -0.50 | 0.000 | NC | NS |
| Calcineurin B homologous protein 1 | CHP1 | -0.46 | 0.000 | NC | NS |
| Chromatin accessibility complex protein 1 | CHRC1 | -0.88 | 0.000 | NC | NS |
| CDGSH iron-sulfur domain-containing protein 1 | CISD1 | -0.31 | 0.000 | -0.79 | 0.004 |
| Citramalyl-CoA lyase | CLYBL | -0.65 | 0.009 | NC | NS |
| Chymase | CMA1 | -0.51 | 0.000 | -0.62 | 0.254 |
| Calponin-1 | CNN1 | 1.07 | 0.000 | NC | NS |
| Complement C3 | CO3 | 0.28 | 0.000 | 0.44 | 0.000 |
| Collagen alpha-1(III) chain | CO3A1 | -0.67 | 0.000 | NC | NS |
| Complement component C7 | CO7 | -0.55 | 0.000 | NC | NS |
| Cofilin-1 | COF1 | -0.30 | 0.000 | 0.54 | 0.001 |
| Cofilin-2 | COF2 | -0.24 | 0.000 | 0.75 | 0.000 |
| Coatomer subunit beta | COPB | -0.62 | 0.000 | 1.11 | 0.002 |
| Cytochrome c oxidase copper chaperone | COX17 | -0.46 | 0.000 | 4.05 | 0.000 |
| Cytochrome c oxidase subunit 7C | COX7C | -0.47 | 0.000 | 0.68 | 0.028 |
| Cytochrome c oxidase subunit 8A | COX8A | -0.66 | 0.000 | NC | NS |
| Copine-1 | CPNE1 | -0.34 | 0.027 | NC | NS |
| Serine/threonine-protein phosphatase CPPED1 | CPPED | 0.33 | 0.033 | 1.09 | 0.001 |
| Carnitine O-palmitoyltransferase 1 | CPT1A | -0.51 | 0.009 | -0.84 | 0.025 |
| Death domain-containing protein CRADD | CRADD | -0.38 | 0.014 | NC | NS |
| Cysteine-rich with EGF-like domain protein 1 | CREL1 | -0.65 | 0.000 | -0.49 | 0.006 |
| Cysteine-rich protein 1 | CRIP1 | 0.89 | 0.005 | NC | NS |
| Cysteine and glycine-rich protein 1 | CSRP1 | 0.57 | 0.036 | NC | NS |
| Choline transporter-like protein 2 | CTL2 | -0.59 | 0.000 | NC | NS |
| Cytochrome c oxidase subunit 7A1 | CX7A1 | -1.04 | 0.000 | -1.25 | 0.000 |
| Dolichyl-diphosphooligosaccharide--protein glycosyltransferase subunit DAD1 | DAD1 | -0.45 | 0.038 | NC | NS |
| DAZ-associated protein 1 | DAZP1 | -0.59 | 0.000 | NC | NS |
| Dynactin subunit 2 | DCTN2 | -0.33 | 0.000 | NC | NS |
| Uroporphyrinogen decarboxylase | DCUP | -0.45 | 0.000 | NC | NS |
| L-xylulose reductase | DCXR | -0.35 | 0.000 | NC | NS |
| N(G)-dimethylarginine dimethylaminohydrolase 1 | DDAH1 | -0.49 | 0.000 | 0.57 | 0.000 |
| 2,4-dienoyl-CoA reductase, mitochondrial | DECR | -0.46 | 0.000 | -0.64 | 0.000 |

| | | | | | |
|---|---|---|---|---|---|
| Dermatopontin | DERM | -0.30 | 0.000 | -1.03 | 0.002 |
| Desmin | DESM | 0.24 | 0.000 | NC | NS |
| Peroxisomal multifunctional enzyme type 2 | DHB4 | -0.31 | 0.001 | 0.79 | 0.000 |
| Dehydrogenase/reductase SDR family member 11 | DHR11 | -0.40 | 0.035 | 1.20 | 0.000 |
| Dehydrogenase/reductase SDR family member 2 | DHRS2 | 0.55 | 0.022 | NC | NS |
| Sorbitol dehydrogenase | DHSO | 0.56 | 0.000 | NC | NS |
| Putative ATP-dependent RNA helicase DHX30 | DHX30 | -0.43 | 0.017 | NC | NS |
| H/ACA ribonucleoprotein complex subunit 4 | DKC1 | -2.00 | 0.000 | NC | NS |
| Dihydrolipoyl dehydrogenase | DLDH | -0.35 | 0.000 | NC | NS |
| DnaJ homolog subfamily B member 2 | DNJB2 | 0.60 | 0.006 | NC | NS |
| Deoxyribonuclease-2-alpha | DNS2A | -0.56 | 0.001 | NC | NS |
| DNA polymerase alpha subunit B | DPOA2 | 0.75 | 0.000 | NC | NS |
| Dihydropyrimidinase-related protein 2 | DPYL2 | -0.22 | 0.004 | NC | NS |
| Dynein light chain 1 | DYL1 | -0.46 | 0.000 | NC | NS |
| Dynein light chain 2 | DYL2 | -0.25 | 0.000 | 0.15 | 0.044 |
| Trifunctional enzyme subunit alpha | ECHA | -0.23 | 0.000 | -0.26 | 0.000 |
| Trifunctional enzyme subunit beta | ECHB | -0.23 | 0.000 | NC | NS |
| Enoyl-CoA hydratase domain-containing protein 2 | ECHD2 | -0.48 | 0.000 | NC | NS |
| Endothelial differentiation-related factor 1 | EDF1 | -0.72 | 0.000 | NC | NS |
| Elongation factor 1-alpha 2 | EF1A2 | -0.47 | 0.000 | NC | NS |
| EF-hand domain-containing protein D1 | EFHD1 | 0.44 | 0.016 | NC | NS |
| EH domain-containing protein 2 | EHD2 | -0.58 | 0.000 | -0.68 | 0.000 |
| Emerin | EMD | -0.49 | 0.000 | -0.76 | 0.058 |
| Transcription and mRNA export factor ENY2 | ENY2 | -0.61 | 0.001 | NC | NS |
| Mammalian ependymin-related protein 1 | EPDR1 | -0.54 | 0.000 | 1.69 | 0.000 |
| Electron transfer flavoprotein subunit alpha | ETFA | -0.30 | 0.000 | -0.30 | 0.000 |
| Electron transfer flavoprotein-ubiquinone oxidoreductase | ETFD | -0.33 | 0.000 | NC | NS |
| Coagulation factor XIII B chain | F13B | -0.45 | 0.045 | NC | NS |
| Fatty acid-binding protein | FABP5 | 0.29 | 0.000 | 0.61 | 0.001 |
| Fatty acid-binding protein | FABPH | 0.39 | 0.000 | 0.38 | 0.004 |
| Acylpyruvase FAHD1 | FAHD1 | 0.38 | 0.000 | NC | NS |
| Fibulin-7 | FBLN7 | -0.73 | 0.047 | NC | NS |
| Formin-binding protein 1-like | FBP1L | 0.92 | 0.000 | NC | NS |
| Alpha-2-HS-glycoprotein | FETUA | 0.33 | 0.000 | -0.86 | 0.000 |
| Four and a half LIM domains protein 2 | FHL2 | -0.35 | 0.000 | -0.73 | 0.042 |
| Fibrinogen alpha chain | FIBA | 0.45 | 0.000 | 0.74 | 0.003 |
| Peptidyl-prolyl cis-trans isomerase FKBP1A | FKB1A | -0.17 | 0.000 | 0.68 | 0.001 |
| Peptidyl-prolyl cis-trans isomerase FKBP2 | FKBP2 | -0.29 | 0.001 | NC | NS |
| Protein flightless-1 homolog | FLII | -0.51 | 0.000 | NC | NS |
| Filamin-A | FLNA | 0.55 | 0.000 | 0.38 | 0.033 |
| Flotillin-1 | FLOT1 | -0.54 | 0.000 | NC | NS |
| Flotillin-2 | FLOT2 | -0.56 | 0.000 | NC | NS |
| Fructosamine-3-kinase | FN3K | -0.34 | 0.000 | NC | NS |
| Glyceraldehyde-3-phosphate dehydrogenase | G3P | 0.35 | 0.000 | NC | NS |
| Guanine nucleotide-binding protein G(I)/G(S)/G(T) subunit beta-1 | GBB1 | -0.32 | 0.048 | NC | NS |
| Guanine nucleotide-binding protein G(I)/G(S)/G(T) subunit beta-2 | GBB2 | -0.38 | 0.000 | NC | NS |

| | | | | | |
|---|---|---|---|---|---|
| Guanine nucleotide-binding protein G(I)/G(S)/G(O) subunit gamma-7 | GBG7 | -0.67 | 0.000 | NC | NS |
| Glutaryl-CoA dehydrogenase | GCDH | -0.23 | 0.000 | NC | NS |
| Glycogen debranching enzyme | GDE | 0.43 | 0.000 | NC | NS |
| Hydroxyacylglutathione hydrolase | GLO2 | 0.25 | 0.000 | 0.35 | 0.006 |
| Glutaredoxin-1 | GLRX1 | 0.18 | 0.000 | NC | NS |
| D-glutamate cyclase | GLUCM | -1.06 | 0.000 | NC | NS |
| Glycogenin-1 | GLYG | 0.35 | 0.000 | 0.79 | 0.000 |
| Guanine nucleotide-binding protein G(i) subunit alpha-2 | GNAI2 | -0.54 | 0.000 | -0.59 | 0.039 |
| Glycerol-3-phosphate acyltransferase 3 | GPAT3 | -0.35 | 0.012 | NC | NS |
| Glycerol-3-phosphate acyltransferase 4 | GPAT4 | -0.55 | 0.002 | NC | NS |
| Glycerol-3-phosphate dehydrogenase 1-like protein | GPD1L | -0.33 | 0.000 | NC | NS |
| Glutathione S-transferase kappa 1 | GSTK1 | -0.40 | 0.000 | -0.68 | 0.000 |
| Glutathione S-transferase Mu 2 | GSTM2 | 0.32 | 0.000 | NC | NS |
| Glutathione S-transferase Mu 3 | GSTM3 | 0.40 | 0.000 | NC | NS |
| Glutathione S-transferase omega-1 | GSTO1 | -0.29 | 0.000 | NC | NS |
| Glycogen [starch] synthase | GYS1 | 0.36 | 0.000 | -0.22 | 0.043 |
| Histone H1.0 | H10 | -0.81 | 0.000 | NC | NS |
| Histone H1x | H1X | -0.63 | 0.000 | NC | NS |
| Core histone macro-H2A.1 | H2AY | -0.60 | 0.000 | NC | NS |
| Histone H4 | H4 | -0.39 | 0.000 | 0.81 | 0.000 |
| Hepatoma-derived growth factor | HDGF | -0.41 | 0.000 | NC | NS |
| Hepatoma-derived growth factor-related protein 3 | HDGR3 | -0.49 | 0.000 | NC | NS |
| Hemopexin | HEMO | 0.28 | 0.000 | NC | NS |
| HLA class II histocompatibility antigen gamma chain | HG2A | -0.75 | 0.001 | NC | NS |
| High mobility group protein HMG-I/HMG-Y | HMGA1 | -0.98 | 0.000 | -1.52 | 0.000 |
| High mobility group protein B1 | HMGB1 | -0.42 | 0.000 | 0.65 | 0.001 |
| Hepatocyte nuclear factor 1-beta | HNF1B | -0.69 | 0.040 | NC | NS |
| Heterogeneous nuclear ribonucleoprotein H | HNRH1 | -0.34 | 0.000 | -0.38 | 0.000 |
| Heterogeneous nuclear ribonucleoprotein H3 | HNRH3 | -0.39 | 0.000 | NC | NS |
| Heterogeneous nuclear ribonucleoprotein U-like protein 1 | HNRL1 | -0.47 | 0.003 | NC | NS |
| Heterogeneous nuclear ribonucleoproteins C1/C2 | HNRPC | -0.65 | 0.000 | NC | NS |
| Heterogeneous nuclear ribonucleoprotein D0 | HNRPD | -0.59 | 0.000 | 0.99 | 0.004 |
| Heterogeneous nuclear ribonucleoprotein K | HNRPK | -0.28 | 0.000 | NC | NS |
| Heterogeneous nuclear ribonucleoprotein M | HNRPM | -0.34 | 0.001 | -1.03 | 0.003 |
| Heterogeneous nuclear ribonucleoprotein U | HNRPU | -0.45 | 0.000 | NC | NS |
| Heterochromatin protein 1-binding protein 3 | HP1B3 | -0.62 | 0.000 | 0.79 | 0.003 |
| Hypoxanthine-guanine phosphoribosyltransferase | HPRT | 0.32 | 0.000 | 0.36 | 0.005 |
| Heat shock 70 kDa protein 1-like | HS71L | -0.50 | 0.000 | NC | NS |
| Heat shock protein beta-6 | HSPB6 | 0.49 | 0.000 | 0.80 | 0.000 |
| Bifunctional epoxide hydrolase 2 | HYES | 0.43 | 0.000 | NC | NS |
| Protein-S-isoprenylcysteine O-methyltransferase | ICMT | -0.96 | 0.029 | NC | NS |
| Isocitrate dehydrogenase [NAD] subunit alpha | IDH3A | -0.32 | 0.000 | NC | NS |
| Eukaryotic translation initiation factor 2 subunit 2 | IF2B | -0.45 | 0.000 | NC | NS |
| Eukaryotic initiation factor 4A-III | IF4A3 | -0.61 | 0.000 | -2.96 | 0.000 |
| Eukaryotic translation initiation factor 4E | IF4E | -0.44 | 0.006 | 0.86 | 0.000 |

| | | | | | |
|---|---|---|---|---|---|
| Immunoglobulin heavy constant gamma 1 | IGHG1 | 0.29 | 0.000 | NC | NS |
| Immunoglobulin heavy constant gamma 2 | IGHG2 | 0.33 | 0.000 | NC | NS |
| Immunoglobulin kappa constant | IGKC | 0.36 | 0.000 | NC | NS |
| Integrin-linked protein kinase | ILK | -0.40 | 0.000 | -0.61 | 0.000 |
| Inorganic pyrophosphatase | IPYR | -0.38 | 0.000 | 0.30 | 0.055 |
| Integrin beta-1 | ITB1 | -0.34 | 0.000 | NC | NS |
| Keratin | K2C5 | 0.51 | 0.042 | 1.16 | 0.000 |
| Keratin | K2C6A | 1.23 | 0.003 | 1.16 | 0.000 |
| KN motif and ankyrin repeat domain-containing protein 2 | KANK2 | -0.38 | 0.000 | -0.67 | 0.002 |
| Creatine kinase M-type | KCRM | 0.22 | 0.000 | NC | NS |
| Creatine kinase S-type | KCRS | -0.39 | 0.000 | -0.58 | 0.000 |
| Pyruvate kinase PKM | KPYM | 0.24 | 0.000 | 0.42 | 0.015 |
| Ribosomal protein S6 kinase beta-1 | KS6B1 | -0.61 | 0.040 | NC | NS |
| Lupus La protein | LA | -0.53 | 0.000 | NC | NS |
| Laminin subunit alpha-2 | LAMA2 | -0.52 | 0.000 | NC | NS |
| Laminin subunit beta-1 | LAMB1 | -0.52 | 0.000 | NC | NS |
| Laminin subunit gamma-1 | LAMC1 | -0.34 | 0.000 | NC | NS |
| L-lactate dehydrogenase A chain | LDHA | 0.31 | 0.000 | 0.38 | 0.007 |
| Galectin-1 | LEG1 | -0.26 | 0.000 | 0.19 | 0.059 |
| Lipoyl synthase | LIAS | -0.50 | 0.011 | NC | NS |
| Platelet-activating factor acetylhydrolase IB subunit alpha | LIS1 | -0.36 | 0.000 | 0.30 | 0.040 |
| Lamin-B2 | LMNB2 | -0.40 | 0.000 | NC | NS |
| U6 snRNA-associated Sm-like protein LSm4 | LSM4 | -0.55 | 0.000 | NC | NS |
| U6 snRNA-associated Sm-like protein LSm6 | LSM6 | -0.52 | 0.000 | NC | NS |
| Ragulator complex protein LAMTOR1 | LTOR1 | -0.46 | 0.000 | NC | NS |
| Ragulator complex protein LAMTOR5 | LTOR5 | -0.47 | 0.014 | NC | NS |
| Lysophospholipase-like protein 1 | LYPL1 | -0.49 | 0.000 | NC | NS |
| Lysozyme C | LYSC | 0.46 | 0.000 | NC | NS |
| Mitochondrial 2-oxoglutarate/malate carrier protein | M2OM | -0.44 | 0.000 | -0.92 | 0.000 |
| Alpha-mannosidase 2C1 | MA2C1 | -0.40 | 0.000 | NC | NS |
| Maleylacetoacetate isomerase | MAAI | 0.43 | 0.000 | -0.58 | 0.001 |
| NADP-dependent malic enzyme | MAOX | 0.41 | 0.000 | NC | NS |
| Mitochondrial carnitine/acylcarnitine carrier protein | MCAT | -0.55 | 0.041 | -1.24 | 0.000 |
| Malignant T-cell-amplified sequence 1 | MCTS1 | -0.39 | 0.000 | NC | NS |
| Malate dehydrogenase | MDHC | 0.16 | 0.000 | 0.22 | 0.008 |
| Mitochondrial fission regulator 1-like | MFR1L | -0.79 | 0.001 | -1.95 | 0.000 |
| MICOS complex subunit MIC13 | MIC13 | -0.50 | 0.000 | NC | NS |
| Migration and invasion enhancer 1 | MIEN1 | 0.55 | 0.000 | NC | NS |
| Melanoma antigen recognized by T-cells 1 | MLANA | 0.84 | 0.021 | NC | NS |
| Methylmalonate-semialdehyde dehydrogenase [acylating] | MMSA | -0.33 | 0.000 | -0.52 | 0.000 |
| Phosphate carrier protein | MPCP | -0.69 | 0.000 | -0.86 | 0.001 |
| Mannose-6-phosphate isomerase | MPI | 0.36 | 0.000 | NC | NS |
| Methyltransferase-like 26 | MTL26 | -0.52 | 0.045 | NC | NS |
| Myosin-11 | MYH11 | 0.79 | 0.000 | 0.31 | 0.006 |
| Myosin regulatory light polypeptide 9 | MYL9 | 0.31 | 0.001 | 0.73 | 0.000 |
| Nicotinate-nucleotide pyrophosphorylase [carboxylating] | NADC | 0.52 | 0.001 | NC | NS |
| N-acetyl-D-glucosamine kinase | NAGK | -0.48 | 0.000 | NC | NS |
| Neutral cholesterol ester hydrolase 1 | NCEH1 | -0.30 | 0.001 | -1.36 | 0.001 |
| NADH dehydrogenase [ubiquinone] 1 alpha subcomplex subunit 1 | NDUA1 | -0.35 | 0.003 | -0.98 | 0.038 |

| | | | | | |
|---|---|---|---|---|---|
| NADH dehydrogenase [ubiquinone] 1 alpha subcomplex subunit 2 | NDUA2 | 0.32 | 0.001 | -0.62 | 0.040 |
| Cytochrome c oxidase subunit NDUFA4 | NDUA4 | -0.47 | 0.000 | -0.84 | 0.000 |
| NADH dehydrogenase [ubiquinone] 1 alpha subcomplex subunit 9 | NDUA9 | -0.39 | 0.000 | -0.17 | 0.056 |
| NADH dehydrogenase [ubiquinone] 1 alpha subcomplex subunit 10 | NDUAA | -0.31 | 0.000 | NC | NS |
| NADH dehydrogenase [ubiquinone] 1 alpha subcomplex subunit 13 | NDUAD | -0.51 | 0.000 | -0.70 | 0.008 |
| NADH dehydrogenase [ubiquinone] 1 alpha subcomplex assembly factor 3 | NDUF3 | -0.34 | 0.000 | 1.16 | 0.000 |
| NADH dehydrogenase [ubiquinone] 1 alpha subcomplex assembly factor 4 | NDUF4 | -0.44 | 0.000 | -0.62 | 0.000 |
| NADH dehydrogenase [ubiquinone] iron-sulfur protein 3 | NDUS3 | -0.24 | 0.000 | NC | NS |
| NEDD8 | NEDD8 | -0.31 | 0.000 | 1.32 | 0.000 |
| Nidogen-2 | NID2 | -0.30 | 0.000 | -0.63 | 0.000 |
| Protein NipSnap homolog 1 | NIPS1 | 0.46 | 0.009 | NC | NS |
| NPC intracellular cholesterol transporter 2 | NPC2 | -0.35 | 0.001 | NC | NS |
| Nucleophosmin | NPM | -0.56 | 0.000 | 0.40 | 0.001 |
| Protein NipSnap homolog 3A | NPS3A | 0.87 | 0.000 | NC | |
| Nuclear transport factor 2 | NTF2 | -0.24 | 0.000 | 0.34 | 0.025 |
| N-terminal Xaa-Pro-Lys N-methyltransferase 1 | NTM1A | -0.69 | 0.000 | NC | NS |
| NADH-ubiquinone oxidoreductase chain 2 | NU2M | -0.39 | 0.003 | -1.23 | 0.007 |
| NADH-ubiquinone oxidoreductase chain 6 | NU6M | -0.63 | 0.000 | NC | NS |
| Iron-sulfur protein NUBPL | NUBPL | 0.28 | 0.019 | NC | NS |
| Endonuclease G | NUCG | -0.53 | 0.003 | -0.25 | 0.030 |
| Nuclear ubiquitous casein and cyclin-dependent kinase substrate 1 | NUCKS | -1.06 | 0.000 | NC | NS |
| ADP-sugar pyrophosphatase | NUDT5 | -0.66 | 0.002 | NC | NS |
| Nucleoporin p58/p45 | NUP58 | -0.79 | 0.048 | NC | NS |
| Immunoglobulin heavy variable 3-7 | P01781 | 0.54 | 0.000 | NC | NS |
| Polyadenylate-binding protein 2 | PABP2 | -0.49 | 0.021 | NC | NS |
| Protein kinase C and casein kinase substrate in neurons protein 3 | PACN3 | -0.41 | 0.000 | NC | NS |
| Poly [ADP-ribose] polymerase 1 | PARP1 | -0.51 | 0.000 | NC | NS |
| Poly(rC)-binding protein 1 | PCBP1 | -0.50 | 0.000 | 0.42 | 0.001 |
| Poly(rC)-binding protein 2 | PCBP2 | -0.41 | 0.031 | NC | NS |
| Prenylcysteine oxidase 1 | PCYOX | 0.30 | 0.000 | NC | NS |
| PDZ and LIM domain protein 3 | PDLI3 | 0.60 | 0.000 | 1.34 | 0.000 |
| Astrocytic phosphoprotein PEA-15 | PEA15 | -0.43 | 0.000 | NC | NS |
| Phosphatidylethanolamine-binding protein 1 | PEBP1 | 0.15 | 0.000 | 0.47 | 0.004 |
| Phosphoglycerate mutase 1 | PGAM1 | 0.25 | 0.000 | 0.66 | 0.005 |
| Phosphoglycerate kinase 1 | PGK1 | 0.18 | 0.000 | 0.17 | 0.033 |
| 14 kDa phosphohistidine phosphatase | PHP14 | -0.41 | 0.000 | NC | NS |
| Plakophilin-2 | PKP2 | -0.41 | 0.000 | NC | NS |
| Junction plakoglobin | PLAK | -0.44 | 0.000 | NC | NS |
| Proteolipid protein 2 | PLP2 | -0.48 | 0.028 | NC | NS |
| Lysosomal protective protein | PPGB | -0.48 | 0.000 | 0.33 | 0.000 |
| Peptidyl-prolyl cis-trans isomerase-like 1 | PPIL1 | -0.45 | 0.000 | NC | NS |
| Palmitoyl-protein thioesterase 1 | PPT1 | -0.64 | 0.000 | 0.94 | 0.000 |
| PRA1 family protein 3 | PRAF3 | -0.61 | 0.000 | -0.66 | 0.003 |
| Peroxiredoxin-2 | PRDX2 | 0.42 | 0.000 | NC | NS |

| | | | | | |
|---|---|---|---|---|---|
| Peroxiredoxin-5 | PRDX5 | -0.27 | 0.000 | -0.28 | 0.009 |
| Prostaglandin-H2 D-isomerase | PTGDS | 0.31 | 0.000 | NC | NS |
| Adenylosuccinate synthetase isozyme 1 | PURA1 | -0.49 | 0.000 | -0.85 | 0.000 |
| Glycogen phosphorylase | PYGB | 0.31 | 0.000 | 0.28 | 0.012 |
| Glycogen phosphorylase | PYGM | 0.36 | 0.001 | 0.24 | 0.012 |
| Cytochrome b-c1 complex subunit 2 | QCR2 | -0.27 | 0.000 | -1.10 | 0.055 |
| Cytochrome b-c1 complex subunit 6 | QCR6 | 0.27 | 0.000 | NC | NS |
| Cytochrome b-c1 complex subunit 8 | QCR8 | -0.21 | 0.012 | -1.18 | 0.000 |
| Cytochrome b-c1 complex subunit 9 | QCR9 | -0.26 | 0.004 | -1.68 | 0.000 |
| Ras-related protein Rab-2A | RAB2A | -0.47 | 0.000 | -0.38 | 0.000 |
| Ras-related protein Rab-7a | RAB7A | -0.41 | 0.000 | NC | NS |
| Ras-related C3 botulinum toxin substrate 1 | RAC1 | -0.40 | 0.000 | 0.32 | 0.021 |
| Receptor of activated protein C kinase 1 | RACK1 | -0.28 | 0.000 | 0.49 | 0.000 |
| RNA-binding protein Raly | RALY | -0.95 | 0.010 | NC | NS |
| Protein RCC2 | RCC2 | -0.93 | 0.024 | NC | NS |
| Retinoid-binding protein 7 | RET7 | -0.52 | 0.000 | NC | NS |
| Ribonuclease inhibitor | RINI | -0.25 | 0.000 | 0.37 | 0.003 |
| Ribonucleoside-diphosphate reductase large subunit | RIR1 | 1.22 | 0.000 | NC | NS |
| 60S ribosomal protein L12 | RL12 | -0.27 | 0.000 | 0.43 | 0.001 |
| 60S ribosomal protein L17 | RL17 | -0.47 | 0.000 | 1.24 | 0.000 |
| 60S ribosomal protein L22 | RL22 | -0.32 | 0.000 | 0.62 | 0.026 |
| 60S ribosomal protein L23 | RL23 | -0.56 | 0.000 | 0.43 | 0.002 |
| 60S ribosomal protein L23a | RL23A | -0.36 | 0.000 | 0.41 | 0.005 |
| 60S ribosomal protein L24 | RL24 | -0.38 | 0.000 | 0.74 | 0.000 |
| 60S ribosomal protein L27 | RL27 | -0.31 | 0.000 | 0.64 | 0.005 |
| 60S ribosomal protein L30 | RL30 | -0.40 | 0.000 | 0.71 | 0.000 |
| 60S ribosomal protein L9 | RL9 | -0.39 | 0.000 | 0.40 | 0.005 |
| 60S acidic ribosomal protein P0 | RLA0 | -0.72 | 0.011 | NC | NS |
| 60S acidic ribosomal protein P1 | RLA1 | -0.38 | 0.003 | NC | NS |
| 39S ribosomal protein L14 | RM14 | -1.39 | 0.000 | NC | NS |
| 39S ribosomal protein L34 | RM34 | -0.75 | 0.003 | NC | NS |
| 39S ribosomal protein L40 | RM40 | -0.52 | 0.001 | NC | NS |
| 39S ribosomal protein L53 | RM53 | -0.64 | 0.006 | NC | NS |
| 39S ribosomal protein L55 | RM55 | -0.58 | 0.003 | NC | NS |
| 60 kDa SS-A/Ro ribonucleoprotein | RO60 | -0.32 | 0.039 | NC | NS |
| Heterogeneous nuclear ribonucleoproteins A2/B1 | ROA2 | -0.47 | 0.000 | -0.75 | 0.000 |
| Heterogeneous nuclear ribonucleoprotein A3 | ROA3 | -0.48 | 0.000 | -0.16 | 0.043 |
| Heterogeneous nuclear ribonucleoprotein A/B | ROAA | -0.50 | 0.000 | NC | NS |
| Reactive oxygen species modulator 1 | ROMO1 | 0.33 | 0.002 | NC | NS |
| Ras-related protein R-Ras | RRAS | -0.43 | 0.013 | NC | NS |
| 40S ribosomal protein S10 | RS10 | -0.48 | 0.000 | NC | NS |
| 40S ribosomal protein S11 | RS11 | -0.34 | 0.000 | NC | NS |
| 40S ribosomal protein S12 | RS12 | -0.34 | 0.001 | 0.81 | 0.021 |
| 40S ribosomal protein S13 | RS13 | -0.31 | 0.005 | NC | NS |
| 40S ribosomal protein S14 | RS14 | -0.30 | 0.000 | NC | NS |
| 40S ribosomal protein S15 | RS15 | -0.62 | 0.000 | NC | NS |
| 40S ribosomal protein S18 | RS18 | -0.33 | 0.000 | NC | NS |
| 40S ribosomal protein S20 | RS20 | -0.50 | 0.000 | -0.43 | 0.001 |
| 40S ribosomal protein S25 | RS25 | -0.37 | 0.000 | NC | NS |
| Ubiquitin-40S ribosomal protein S27a | RS27A | -0.31 | 0.000 | 1.20 | 0.047 |
| 40S ribosomal protein S3 | RS3 | -0.46 | 0.000 | 0.21 | 0.026 |
| 40S ribosomal protein S3a | RS3A | -0.38 | 0.000 | 0.26 | 0.022 |
| 40S ribosomal protein S8 | RS8 | -0.32 | 0.000 | NC | NS |
| Reticulon-4 | RTN4 | 0.43 | 0.008 | 1.10 | 0.000 |
| Small nuclear ribonucleoprotein E | RUXE | -0.57 | 0.000 | -0.40 | 0.008 |
| Protein S100-B | S100B | -0.69 | 0.000 | NC | NS |
| Protein S100-A1 | S10A1 | 0.46 | 0.000 | NC | NS |

| | | | | | |
|---|---|---|---|---|---|
| Protein S100-A8 | S10A8 | 0.69 | 0.000 | 4.78 | 0.000 |
| Protein S100-A9 | S10A9 | 0.79 | 0.000 | 5.62 | 0.000 |
| Protein S100-A10 | S10AA | -0.25 | 0.001 | NC | NS |
| Prosaposin | SAP | -0.40 | 0.000 | -5.13 | 0.001 |
| SAP domain-containing ribonucleoprotein | SARNP | -0.66 | 0.000 | NC | NS |
| Diamine acetyltransferase 2 | SAT2 | -0.55 | 0.006 | 3.50 | 0.000 |
| Sec1 family domain-containing protein 1 | SCFD1 | -0.70 | 0.004 | NC | NS |
| Lysosome membrane protein 2 | SCRB2 | -0.56 | 0.000 | -1.79 | 0.000 |
| Selenoprotein H | SELH | -0.57 | 0.032 | NC | NS |
| Protein SET | SET | -0.40 | 0.000 | 0.75 | 0.010 |
| SH3 domain-binding glutamic acid-rich-like protein 2 | SH3L2 | 0.62 | 0.015 | NC | NS |
| Endophilin-B1 | SHLB1 | -0.43 | 0.000 | 0.28 | 0.000 |
| Sialic acid synthase | SIAS | -0.29 | 0.000 | 0.95 | 0.007 |
| Structural maintenance of chromosomes protein 6 | SMC6 | -0.61 | 0.034 | NC | NS |
| Small nuclear ribonucleoprotein Sm D2 | SMD2 | -0.33 | 0.000 | -1.24 | 0.002 |
| Small nuclear ribonucleoprotein Sm D3 | SMD3 | -0.37 | 0.000 | NC | NS |
| Alpha-soluble NSF attachment protein | SNAA | -0.38 | 0.006 | -1.16 | 0.000 |
| Superoxide dismutase [Cu-Zn] | SODC | 0.36 | 0.000 | 0.48 | 0.002 |
| Signal peptidase complex subunit 2 | SPCS2 | -0.50 | 0.000 | NC | NS |
| Signal peptidase complex subunit 3 | SPCS3 | -0.59 | 0.001 | NC | NS |
| SPARC-like protein 1 | SPRL1 | -0.63 | 0.000 | -1.31 | 0.002 |
| Spectrin beta chain | SPTB2 | -0.42 | 0.000 | -1.09 | 0.000 |
| Sulfide:quinone oxidoreductase | SQOR | -0.48 | 0.000 | NC | |
| Sarcalumenin | SRCA | -0.32 | 0.000 | -1.15 | 0.000 |
| Sarcoplasmic reticulum histidine-rich calcium-binding protein | SRCH | 0.48 | 0.000 | NC | NS |
| Signal recognition particle 9 kDa protein | SRP09 | -0.55 | 0.001 | NC | NS |
| Signal recognition particle 14 kDa protein | SRP14 | -0.76 | 0.000 | NC | NS |
| Succinate--CoA ligase [ADP-forming] subunit beta | SUCB1 | -0.58 | 0.000 | -0.47 | 0.000 |
| Aspartate--tRNA ligase | SYDM | 0.41 | 0.000 | NC | NS |
| Transaldolase | TALDO | -0.33 | 0.000 | NC | NS |
| T-complex protein 1 subunit beta | TCPB | -0.55 | 0.004 | NC | NS |
| T-complex protein 1 subunit delta | TCPD | -0.75 | 0.001 | -0.18 | 0.006 |
| T-complex protein 1 subunit epsilon | TCPE | -0.56 | 0.000 | -0.26 | 0.009 |
| Translationally-controlled tumor protein | TCTP | -0.27 | 0.000 | 0.32 | 0.004 |
| Prostaglandin E synthase 3 | TEBP | -0.56 | 0.000 | 0.68 | 0.008 |
| Dual specificity testis-specific protein kinase 2 | TESK2 | -0.60 | 0.025 | NC | NS |
| Acetyl-CoA acetyltransferase | THIL | -0.26 | 0.000 | NC | NS |
| 3-ketoacyl-CoA thiolase | THIM | -0.19 | 0.000 | -0.52 | 0.000 |
| Mitochondrial import inner membrane translocase subunit Tim13 | TIM13 | 0.22 | 0.000 | NC | NS |
| Talin-2 | TLN2 | -0.38 | 0.000 | NC | NS |
| Transmembrane protein 245 | TM245 | -0.61 | 0.027 | NC | NS |
| Calcium load-activated calcium channel | TMCO1 | -0.89 | 0.008 | NC | NS |
| Triosephosphate isomerase | TPIS | 0.21 | 0.000 | 0.33 | 0.003 |
| Tropomyosin beta chain | TPM2 | 0.42 | 0.000 | 1.79 | 0.000 |
| Tropomyosin alpha-3 chain | TPM3 | 0.82 | 0.000 | 1.37 | 0.001 |
| Tripeptidyl-peptidase 1 | TPP1 | -0.28 | 0.000 | NC | NS |
| Serotransferrin | TRFE | 0.32 | 0.000 | -0.89 | 0.000 |
| Lactotransferrin | TRFL | 1.06 | 0.000 | NC | NS |
| Transthyretin | TTHY | -0.31 | 0.001 | 1.28 | 0.009 |
| NEDD8-conjugating enzyme Ubc12 | UBC12 | 0.39 | 0.026 | NC | NS |

| | | | | | |
|---|---|---|---|---|---|
| SUMO-conjugating enzyme UBC9 | UBC9 | -0.38 | 0.000 | NC | NS |
| Ubiquitin carboxyl-terminal hydrolase 14 | UBP14 | -0.22 | 0.032 | NC | NS |
| Ubiquitin carboxyl-terminal hydrolase 8 | UBP8 | 1.19 | 0.017 | NC | NS |
| Ubiquinol-cytochrome-c reductase complex assembly factor 3 | UQCC3 | -0.66 | 0.023 | NC | NS |
| Vesicle-associated membrane protein-associated protein B/C | VAPB | -0.46 | 0.000 | NC | NS |
| Synaptic vesicle membrane protein VAT-1 homolog | VAT1 | -0.33 | 0.000 | NC | NS |
| Vinculin | VINC | -0.49 | 0.000 | 0.22 | 0.000 |
| Visinin-like protein 1 | VISL1 | -0.50 | 0.037 | NC | NS |
| Wings apart-like protein homolog | WAPL | -0.30 | 0.033 | NC | NS |
| Nuclease-sensitive element-binding protein 1 | YBOX1 | -0.56 | 0.000 | NC | NS |
| Y-box-binding protein 3 | YBOX3 | -0.50 | 0.000 | 0.65 | 0.032 |
| Zinc finger CCCH-type antiviral protein 1-like | ZCCHL | -0.72 | 0.019 | NC | NS |

NA = no appropriate NC = no change NS = not significant

**Supplemental table3:** List of MS data quantifying phosphorylation of proteins able to regulate protein expression. Shown are the fold change values (log2) of HFpEF compared to control and for CDC compared to placebo treated HFpEF for total protein concentration if significant (p<0.05; n=3–4) and significant phosphorylation changes (n=6).

| Protein Name | Gene Name | Role | HFpEF vs control FC (log2) | HFpEF vs control p-value | CDC vs placebo FC (log2) | CDC vs placebo p-value | Phospho-site | HFpEF vs control FC (log2) | HFpEF vs control p-value | CDC vs placebo FC (log2) | CDC vs placebo p-value |
|---|---|---|---|---|---|---|---|---|---|---|---|
| Alanyl-tRNA editing protein Aarsd1 | Aarsd 1 | Translation | NC | NS | NC | NS | S88 | 2.06 | 0.008 | NC | NS |
| Actin-binding LIM protein 1 | Ablim1 | Transcription | 0.46 | 0.003 | -0.40 | 0.005 | S115 | NC | NS | 1.31 | 0.041 |
| Arf-GAP domain and FG repeat-containing protein 1 | Agfg1 | RNA trafficking | -0.83 | 0.060 | NC | NS | T177 | 0.79 | 0.020 | NC | NS |
| Neuroblast differentiation-associated protein AHNAK | Ahnak | RNA splicing | NC | NS | NC | NS | S116 (hu) | 0.95 | 0.040 | NC | NS |
| Neuroblast differentiation-associated protein AHNAK | Ahnak | RNA splicing | NC | NS | NC | NS | S211 (hu) | 0.91 | 0.010 | NC | NS |
| Neuroblast differentiation-associated protein AHNAK | Ahnak | RNA splicing | NC | NS | NC | NS | S217 (hu) | 1.57 | 0.057 | NC | NS |
| Neuroblast differentiation-associated protein AHNAK | Ahnak | RNA splicing | NC | NS | NC | NS | T219 (hu) | 1.57 | 0.059 | NC | NS |
| Neuroblast differentiation- | Ahnak | RNA splicing | NC | NS | NC | NS | S889 (hu) | 0.95 | 0.054 | NC | NS |

| | | | | | | | | | | | |
|---|---|---|---|---|---|---|---|---|---|---|---|
| associated protein AHNAK | | | | | | | | | | | |
| Neuroblast differentiation-associated protein AHNAK | Ahnak | RNA splicing | NC | NS | NC | NS | S2050 (hu) | 1.94 | 0.037 | NC | NS |
| Neuroblast differentiation-associated protein AHNAK | Ahnak | RNA splicing | NC | NS | NC | NS | S2245 (hu) | 1.73 | 0.009 | NC | NS |
| Neuroblast differentiation-associated protein AHNAK | Ahnak | RNA splicing | NC | NS | NC | NS | S2499 (hu) | 2.26 | 0.005 | NC | NS |
| Neuroblast differentiation-associated protein AHNAK | Ahnak | RNA splicing | NC | NS | NC | NS | S4893 (hu) | NC | NS | -0.88 | 0.059 |
| Neuroblast differentiation-associated protein AHNAK | Ahnak | RNA splicing | NC | NS | NC | NS | S5218 (hu) | 1.07 | 0.013 | NC | NS |
| Neuroblast differentiation-associated protein AHNAK | Ahnak | RNA splicing | NC | NS | NC | NS | S5233 (hu) | 2.44 | 0.021 | NC | NS |
| Neuroblast differentiation-associated protein AHNAK | Ahnak | RNA splicing | NC | NS | NC | NS | S5243 (hu) | 1.47 | 0.011 | NC | NS |
| Neuroblast differentiation-associated protein AHNAK | Ahnak | RNA splicing | NC | NS | NC | NS | S5244 (hu) | 2.44 | 0.021 | NC | NS |
| Neuroblast differentiation-associated protein AHNAK | Ahnak | RNA splicing | NC | NS | NC | NS | S5271 (hu) | 2.43 | 0.035 | NC | NS |
| Neuroblast differentiation-associated protein AHNAK | Ahnak | RNA splicing | NC | NS | NC | NS | T5275 (hu) | 2.43 | 0.035 | NC | NS |
| Neuroblast differentiation-associated protein AHNAK | Ahnak | RNA splicing | NC | NS | NC | NS | T5275 (hu) | 1.48 | 0.014 | NC | NS |
| RAC-alpha serine/threonine-protein kinase | Akt1 | Transcription | -0.29 | 0.030 | NC | NS | S126 | NC | NS | -2.09 | 0.021 |
| RAC-alpha serine/threonine-protein kinase | Akt1 | Transcription | -0.29 | 0.030 | NC | NS | S129 | NC | NS | -2.09 | 0.021 |

| | | | | | | | | | | | |
|---|---|---|---|---|---|---|---|---|---|---|---|
| DNA-(apurinic or apyrimidinic site) lyase | Apex1 | mRNA stability | NC | NS | NC | NS | S13 | 0.79 | 0.039 | NC | NS |
| Rho GTPase-activating protein 35 | Arhgap35 | Transcription | NC | NS | NC | NS | S1179 | 1.00 | 0.025 | NC | NS |
| Armadillo repeat-containing X-linked protein 3 | Armcx3 | Transcription | NC | NS | NC | NS | S61 | 1.00 | 0.028 | NC | NS |
| BAG family molecular chaperone regulator 3 | Bag3 | Transcription | 0.67 | 0.000 | NC | NS | S176 | 0.86 | 0.059 | NC | NS |
| BAG family molecular chaperone regulator 3 | Bag3 | Transcription | 0.67 | 0.000 | NC | NS | S267 | NC | NS | -1.40 | 0.058 |
| BAG family molecular chaperone regulator 3 | Bag3 | Transcription | 0.67 | 0.000 | NC | NS | S388 | 0.78 | 0.029 | NC | NS |
| BAG family molecular chaperone regulator 3 | Bag3 | Transcription | 0.67 | 0.000 | NC | NS | S401 | 1.12 | 0.016 | NC | NS |
| BAG family molecular chaperone regulator 3 | Bag3 | Transcription | 0.67 | 0.000 | NC | NS | S563 | 0.94 | 0.012 | NC | NS |
| BAG family molecular chaperone regulator 3 | Bag3 | Transcription | 0.67 | 0.000 | NC | NS | T557 | 0.62 | 0.024 | NC | NS |
| Bcl-2-associated transcription factor 1 | Bclaf1 | Transcription | NC | NS | NC | NS | S658 | 0.86 | 0.030 | NC | NS |
| Bcl-2-associated transcription factor 1 | Bclaf1 | Transcription | NC | NS | NC | NS | S660 | 0.92 | 0.030 | NC | NS |
| Calcium-regulated heat stable protein 1 | Carhsp1 | mRNA stability | 0.42 | 0.010 | NC | NS | S32 | NC | NS | 1.07 | 0.098 |
| Calcium-regulated heat stable protein 1 | Carhsp1 | mRNA stability | 0.42 | 0.010 | NC | NS | S41 | 0.65 | 0.036 | NC | NS |
| Caveolae-associated protein 1 | Cavin1 | Transcription | NC | NS | NC | NS | S38 | 1.52 | 0.002 | -0.93 | 0.045 |
| Caveolae-associated protein 1 | Cavin1 | Transcription | NC | NS | NC | NS | T40 | 1.52 | 0.002 | -0.93 | 0.045 |

| | | | | | | | | | | | |
|---|---|---|---|---|---|---|---|---|---|---|---|
| Caveolae-associated protein 1 | Cavin1 | Transcription | NC | NS | NC | NS | S38 | 1.18 | 0.018 | NC | NS |
| Caveolae-associated protein 1 | Cavin1 | Transcription | NC | NS | NC | NS | S42 | 1.18 | 0.018 | NC | NS |
| Cell cycle and apoptosis regulator protein 2 | Ccar2 | mRNA processing | NC | NS | NC | NS | S612 | 0.79 | 0.049 | NC | NS |
| Cold shock domain-containing protein C2 | Csdc2 | mRNA processing | NC | NS | -7.11 | 0.000 | S48 | NC | NS | -1.33 | 0.002 |
| Cysteine and glycine-rich protein 3 | Csrp3 | Transcription | 1.54 | 0.000 | -0.63 | 0.000 | S111 | 1.43 | 0.019 | NC | NS |
| Catenin beta-1 | Ctnnb 1 | Transcription | NC | NS | NC | NS | S191 | 0.92 | 0.042 | NC | NS |
| DnaJ homolog subfamily C member 21 | Dnajc2 1 | rRNA regulation | NC | NS | NC | NS | S46 | 3.79 | 0.013 | NC | NS |
| Segment polarity protein dishevelled homolog DVL-3 | Dvl3 | Transcription | NC | NS | NC | NS | S598 | 1.98 | 0.054 | NC | NS |
| Segment polarity protein dishevelled homolog DVL-3 | Dvl3 | Transcription | NC | NS | NC | NS | T608 | 1.98 | 0.054 | NC | NS |
| Segment polarity protein dishevelled homolog DVL-3 | Dvl3 | Transcription | NC | NS | NC | NS | T609 | 1.98 | 0.054 | NC | NS |
| Enhancer of mRNA-decapping protein 4 | Edc4 | mRNA degradation | NC | NS | NC | NS | T732 | 0.72 | 0.015 | NC | NS |
| Enhancer of mRNA-decapping protein 4 | Edc4 | mRNA degradation | NC | NS | NC | NS | S734 | 0.73 | 0.014 | NC | NS |
| Enhancer of mRNA-decapping protein 4 | Edc4 | mRNA degradation | NC | NS | NC | NS | S678 | 0.70 | 0.041 | NC | NS |
| Elongation factor 2 | Eef2 | Translation | 0.36 | 0.000 | NC | NS | T59 | NC | NS | -2.16 | 0.072 |
| Eukaryotic translation initiation factor 4 gamma 1 | Eif4g1 | Translation | NC | NS | NC | NS | S1079 | 1.30 | 0.058 | NC | NS |

| | | | | | | | | | | | |
|---|---|---|---|---|---|---|---|---|---|---|---|
| Eukaryotic translation initiation factor 4 gamma 1 | Eif4g1 | Translation | NC | NS | NC | NS | S1595 | -2.14 | 0.025 | NC | NS |
| Eukaryotic translation initiation factor 4 gamma 3 | Eif4g3 | Translation | 1.67 | 0.002 | NC | NS | S305 | 1.57 | 0.007 | NC | NS |
| Eukaryotic translation initiation factor 5B | Eif5b | Translation | NC | NS | NC | NS | S137 | 3.55 | 0.010 | -2.13 | 0.058 |
| Alpha-enolase | Eno1 | Transcription | 0.51 | 0.000 | -0.12 | 0.051 | S419 | NC | NS | -0.80 | 0.078 |
| FH1/FH2 domain-containing protein 1 | Fhod1 | Transcription | NC | NS | NC | NS | S141 | 1.19 | 0.034 | NC | NS |
| Fragile X mental retardation syndrome-related protein 1 | Fxr1 | Translation | 0.36 | 0.049 | NC | NS | S433 | 1.45 | 0.029 | NC | NS |
| Ras GTPase-activating protein-binding protein 1 | G3bp1 | Cleaves mRNA | 0.31 | 0.042 | -0.32 | 0.025 | S229 | 0.75 | 0.034 | NC | NS |
| Ras GTPase-activating protein-binding protein 1 | G3bp1 | Cleaves mRNA | 0.31 | 0.042 | -0.32 | 0.025 | S231 | 0.74 | 0.036 | NC | NS |
| Glyceraldehyde-3-phosphate dehydrogenase | Gapdh | Translation | NC | NS | NC | NS | T180 | 0.96 | 0.057 | NC | NS |
| Glyceraldehyde-3-phosphate dehydrogenase | Gapdh | Translation | NC | NS | NC | NS | S208 | NC | NS | -2.20 | 0.037 |
| Histone H1.4 | Hist1h1e | Transcription | NC | NS | NC | NS | S36 | NC | NS | 0.70 | 0.099 |
| Heterogeneous nuclear ribonucleoprotein K | Hnrnpk | RNA splicing | NC | NS | NC | NS | S284 | NC | NS | -0.77 | 0.092 |
| HIV TAT specific factor 1 | Htatsf1 | mRNA splicing | NC | NS | NC | NS | S455 | 2.41 | 0.043 | NC | NS |
| Junction plakoglobin | Jup | Transcription | NC | NS | NC | NS | T78 | 1.86 | 0.059 | NC | NS |
| La-related protein 1 | Larp1 | Translation | NC | NS | NC | NS | S610 | 0.78 | 0.041 | NC | NS |
| Leucine-rich repeat flightless-interacting protein 2 | Lrrfip2 | Transcription | NC | NS | NC | NS | S133 | 1.26 | 0.035 | NC | NS |

| | | | | | | | | | | | |
|---|---|---|---|---|---|---|---|---|---|---|---|
| Matrin-3 | Matr3 | Posttranscriptional regulation | -1.60 | 0.000 | NC | NS | S598 | NC | NS | -1.50 | 0.062 |
| Myocyte-specific enhancer factor 2C | Mef2c | Transcription | NC | NS | NC | NS | S222 | 1.15 | 0.020 | NC | NS |
| Myeloid leukemia factor 1 | Mlf1 | Transcription | NC | NS | NC | NS | S34 | 0.99 | 0.020 | NC | NS |
| Muscular LMNA-interacting protein | Mlip | Transcription | NC | NS | 0.85 | 0.000 | S157 | 0.66 | 0.053 | NC | NS |
| Muscular LMNA-interacting protein | Mlip | Transcription | NC | NS | 0.85 | 0.000 | S721 | 1.69 | 0.037 | NC | NS |
| Metastasis-associated protein MTA2 | Mta2 | Transcription | NC | NS | NC | NS | S435 | 2.20 | 0.021 | NC | NS |
| Nascent polypeptide-associated complex subunit alpha, muscle-specific form | Naca | Transcription | NC | NS | NC | NS | T1533 | NC | NS | -1.47 | 0.025 |
| Nuclear factor NF-kappa-B p105 subunit | Nfkb1 | Transcription | NC | NS | NC | NS | S493 | 0.82 | 0.033 | NC | NS |
| Protein Niban | Niban 1 | Translation | NC | NS | NC | NS | S580 | 0.78 | 0.026 | NC | NS |
| Nuclear speckle splicing regulatory protein 1 | Nsrp1 | mRNA processing | NC | NS | NC | NS | S33 | 0.95 | 0.043 | NC | NS |
| Nuclear ubiquitous casein and cyclin-dependent kinase substrate 1 | Nucks 1 | Transcription | NC | NS | NC | NS | S58 | 1.60 | 0.048 | NC | NS |
| Nuclear ubiquitous casein and cyclin-dependent kinase substrate 1 | Nucks 1 | Transcription | NC | NS | NC | NS | S61 | 1.60 | 0.048 | NC | NS |
| Putative transcription factor Ovo-like 1 | Ovol1 | Transcription | NC | NS | NC | NS | S239 (ms) | 1.60 | 0.055 | NC | NS |
| Poly(rC)-binding protein 2 | Pcbp2 | Intracellular ribonucleo-protein complex | NC | NS | NC | NS | S183 | 0.93 | 0.021 | NC | NS |
| Poly(rC)-binding protein 2 | Pcbp2 | Intracellular ribonucleopr | NC | NS | NC | NS | S185 | 0.92 | 0.020 | NC | NS |

| | | | | | | | | | | | |
|---|---|---|---|---|---|---|---|---|---|---|---|
| | | otein complex | | | | | | | | | |
| PDZ and LIM domain protein 1 | Pdlim1 | Transcription | 0.81 | 0.000 | -0.20 | 0.000 | Y142 | 0.79 | 0.037 | NC | NS |
| PDZ and LIM domain protein 1 | Pdlim1 | Transcription | 0.81 | 0.000 | -0.20 | 0.000 | S144 | 0.81 | 0.030 | NC | NS |
| Prohibitin-2 | Phb2 | Transcription | -0.84 | 0.001 | NC | NS | S151 | 1.48 | 0.056 | NC | NS |
| DNA-directed RNA polymerase III subunit RPC7 | Polr3g | Transcription | NC | NS | NC | NS | S462 | 1.60 | 0.021 | NC | NS |
| Protein phosphatase 1 regulatory subunit 12A | Ppp1r 12a | Transcription | 0.53 | 0.000 | -0.41 | 0.002 | S873 | 0.81 | 0.041 | NC | NS |
| RelA-associated inhibitor | Ppp1r 13l | Transcription | NC | NS | NC | NS | S397 | 0.91 | 0.056 | NC | NS |
| Pumilio homolog 2 | Pum2 | mRNA stability | NC | NS | NC | NS | S136 | 1.38 | 0.015 | NC | NS |
| Transcriptional activator protein Pur-beta | Purb | Transcription | NC | NS | NC | NS | S104 | NC | NS | -0.85 | 0.083 |
| R3H domain-containing protein 2 | R3hd m2 | RNA binding | NC | NS | NC | NS | S362 | NC | NS | 1.16 | 0.095 |
| R3H domain-containing protein 2 | R3hd m2 | RNA binding | NC | NS | NC | NS | S363 | 1.22 | 0.022 | NC | NS |
| R3H domain-containing protein 2 | R3hd m2 | RNA binding | NC | NS | NC | NS | T873 | -0.68 | 0.037 | 0.64 | 0.036 |
| Ral GTPase-activating protein subunit alpha-1 | Ralga pa1 | Transcription | NC | NS | NC | NS | S772 | 1.19 | 0.009 | NC | NS |
| Ral GTPase-activating protein subunit alpha-1 | Ralga pa1 | Transcription | NC | NS | NC | NS | T797 | NC | NS | -1.00 | 0.060 |
| Putative RNA-binding protein 15 | Rbm1 5 | Transcription | NC | NS | NC | NS | S293 | 1.21 | 0.051 | NC | NS |
| RNA-binding protein 20 | Rbm2 0 | RNA splicing | NC | NS | NC | NS | S789 | NC | NS | -0.69 | 0.090 |
| RNA-binding protein 20 | Rbm2 0 | RNA splicing | NC | NS | NC | NS | S1034 | NC | NS | -0.59 | 0.083 |
| 60S ribosomal protein L23a | Rpl23 a | rRNA binding | 0.41 | 0.005 | -0.39 | 0.004 | S43 | 1.18 | 0.033 | NC | NS |
| 60S acidic ribosomal protein P0 | Rplp0 | Ribosome biogenesis | NC | NS | NC | NS | S304 | NC | NS | 2.69 | 0.096 |

| | | | | | | | | | | | |
|---|---|---|---|---|---|---|---|---|---|---|---|
| 60S acidic ribosomal protein P0 | Rplp0 | Ribosome biogenesis | NC | NS | NC | NS | S307 | NC | NS | 2.69 | 0.096 |
| 40S ribosomal protein S17 | Rps17 | rRNA processing | NC | NS | 0.17 | 0.030 | S115 | 1.57 | 0.038 | NC | NS |
| 40S ribosomal protein S3 | Rps3 | Translation | NC | NS | NC | NS | T221 | 1.26 | 0.005 | NC | NS |
| Helicase SKI2W | Skiv2l | mRNA catabolic process | NC | NS | NC | NS | S250 | 0.93 | 0.022 | NC | NS |
| SWI/SNF complex subunit SMARCC2 | Smarc c2 | Transcription | NC | NS | NC | NS | S347 | 1.46 | 0.035 | NC | NS |
| U5 small nuclear ribonucleoprotei n 200 kDa helicase | Snrnp 200 | mRNA splicing | NC | NS | NC | NS | S225 | 1.77 | 0.038 | NC | NS |
| Helicase SRCAP | Srcap | Transcription | NC | NS | NC | NS | S196 | 2.21 | 0.031 | NC | NS |
| Serine/arginine repetitive matrix protein 1 | Srrm1 | mRNA splicing | NC | NS | NC | NS | S400 | NC | NS | -1.89 | 0.063 |
| Serine/arginine repetitive matrix protein 1 | Srrm1 | mRNA splicing | NC | NS | NC | NS | S401 | NC | NS | -1.89 | 0.063 |
| Serine/arginine repetitive matrix protein 1 | Srrm1 | mRNA splicing | NC | NS | NC | NS | S620 | NC | NS | -2.67 | 0.016 |
| Serine/arginine repetitive matrix protein 1 | Srrm1 | mRNA splicing | NC | NS | NC | NS | S741 | 1.58 | 0.028 | NC | NS |
| Serine/arginine repetitive matrix protein 2 | Srrm2 | mRNA splicing | NC | NS | NC | NS | T234 | 1.13 | 0.020 | NC | NS |
| Serine/arginine repetitive matrix protein 2 | Srrm2 | mRNA splicing | NC | NS | NC | NS | S235 | 0.83 | 0.045 | NC | NS |
| Serine/arginine repetitive matrix protein 2 | Srrm2 | mRNA splicing | NC | NS | NC | NS | S348 | NC | NS | -1.94 | 0.091 |
| Serine/arginine repetitive matrix protein 2 | Srrm2 | mRNA splicing | NC | NS | NC | NS | S350 | NC | NS | -1.94 | 0.091 |
| Serine/arginine repetitive matrix protein 2 | Srrm2 | mRNA splicing | NC | NS | NC | NS | S1069 | 1.46 | 0.060 | NC | NS |
| Serine/arginine repetitive matrix protein 2 | Srrm2 | mRNA splicing | NC | NS | NC | NS | S1070 | 1.51 | 0.056 | NC | NS |

| | | | | | | | | | | | |
|---|---|---|---|---|---|---|---|---|---|---|---|
| Serine/arginine repetitive matrix protein 2 | Srrm2 | mRNA splicing | NC | NS | NC | NS | S1153 | 1.22 | 0.030 | NC | NS |
| Serine/arginine repetitive matrix protein 2 | Srrm2 | mRNA splicing | NC | NS | NC | NS | S1280 | 0.79 | 0.031 | NC | NS |
| Serine/arginine-rich splicing factor 6 | Srsf6 | mRNA splicing | NC | NS | NC | NS | S303 | 1.63 | 0.039 | NC | NS |
| Striatin-3 | Strn3 | Transcription | -1.56 | 0.067 | NC | NS | S255 | 1.43 | 0.022 | NC | NS |
| Transcription initiation factor TFIID subunit 9B | Taf9b | Transcription | NC | NS | NC | NS | S377 | NC | NS | -1.53 | 0.092 |
| Thyroid hormone receptor-associated protein 3 | Thrap 3 | RNA splicing | -0.49 | 0.070 | NC | NS | S243 | 0.96 | 0.050 | NC | NS |
| Thyroid hormone receptor-associated protein 3 | Thrap 3 | RNA splicing | -0.49 | 0.070 | NC | NS | S679 | 0.62 | 0.027 | NC | NS |
| Spermatogenesis-defective protein 39 homolog | Vipas39 | Transcription | NC | NS | NC | NS | S119 | 1.13 | 0.045 | NC | NS |
| Transcriptional coactivator YAP1 | Yap1 | Transcription | 1.13 | 0.003 | NC | NS | S145 | NC | NS | -0.88 | 0.066 |
| Zinc finger MYM-type protein 4 | Zmym 4 | Transcription | NC | NS | NC | NS | S245 | 1.15 | 0.059 | NC | NS |

NA = no appropriate NC = no change NS = not significant

**Supplemental table 4**: Listed are significant changed NSPs following PKC isoform inhibition. Proteins were enriched via L-Azidohomoalanine labeling and quantified by LC-MS/MS. Shown are the fold change values (log2) of PKC isoform inhibition compared to control for hypertrophic and WT H9C2 cells if significant ($p<0.05$; n=6) and trending ($p<0.1$; n=6) changes.

| Protein name | Gene name | PKCalpha + PE vs PE | | PKCalpha vs control | | PKCbeta + PE vs PE | | PKCbeta vs control | | PKCdelta + PE vs PE | | PKCdelta vs control | |
|---|---|---|---|---|---|---|---|---|---|---|---|---|---|
| | | Fold-change (log2) | p-value | Fold-change (log2) | p-value | Fold-change (log2) | p-value | Fold-change (log2) | p-value | Fold-change (log2) | p-value | Fold-change (log2) | p-value |
| Actin, alpha skeletal muscle | Acta1 | NC | NS | NC | NS | NC | NS | 1.17 | 0.05 | NC | NS | NC | NS |
| Actin, cytoplasmic 2 | Actg1 | NC | NS | NC | NS | NC | NS | 1.44 | 0.05 | NC | NS | NC | NS |
| Actin, gamma-enteric | Actg2 | NC | NS | 1.13 | 0.07 | NC | NS | 1.17 | 0.04 | NC | NS | NC | NS |

| | | | | | | | | | | | | | |
|---|---|---|---|---|---|---|---|---|---|---|---|---|---|
| smooth muscle | | | | | | | | | | | | | |
| Alpha-actinin-1 | Actn1 | NC | NS | NC | NS | NC | NS | 2.12 | 0.04 | NC | NS | NC | NS |
| Alpha-actinin-4 | Actn4 | NC | NS | NC | NS | NC | NS | 0.79 | 0.07 | NC | NS | NC | NS |
| Annexin A2 | Anxa2 | NC | NS | NC | NS | NC | NS | 0.84 | 0.07 | NC | NS | NC | NS |
| ADP-ribosylation factor 2 | Arf2 | NC | NS | NC | NS | NC | NS | 0.82 | 0.10 | NC | NS | NC | NS |
| Atlastin-3 | Atl3 | NC | NS | 1.02 | 0.02 | NC | NS | 0.98 | 0.02 | NC | NS | NC | NS |
| Sodium/potassium-transporting ATPase subunit alpha-1 | Atp1a1 | NC | NS | 0.88 | 0.05 | NC | NS | 0.79 | 0.07 | NC | NS | NC | NS |
| Sarcoplasmic/endoplasmic reticulum calcium ATPase 1 | Atp2a1 | NC | NS | NC | NS | NC | NS | 2.26 | 0.03 | NC | NS | NC | NS |
| Sarcoplasmic/endoplasmic reticulum calcium ATPase 2 | Atp2a2 | NC | NS | NC | NS | NC | NS | 0.72 | 0.09 | NC | NS | NC | NS |
| ATP synthase subunit beta | Atp5f1b | NC | NS | 3.27 | 0.04 | NC | NS | 3.73 | 0.01 | NC | NS | 2.49 | 0.03 |
| V-type proton ATPase catalytic subunit A | Atp6v1a | Inf | NA | NC | NS | NC | NS | NC | NS | NC | NS | NC | NS |
| Calumenin | Calu | NC | NS | NC | NS | NC | NS | 1.60 | 0.07 | NC | NS | NC | NS |
| Collagen alpha-2 | Col1a1 | NC | NS | NC | NS | NC | NS | 0.93 | 0.09 | 0.91 | 0.09 | 2.44 | 0.02 |
| Collagen alpha-2 | Col1a2 | NC | NS | NC | NS | NC | NS | NC | NS | NC | NS | 0.76 | 0.09 |
| Collagen alpha-1 | Col5a1 | NC | NS | 0.90 | 0.07 | NC | NS | 0.80 | 0.09 | NC | NS | Inf | NA |
| Cathepsin D | Ctsd | NC | NS | NC | NS | 0.98 | 0.07 | NC | NS | 1.19 | 0.03 | NC | NS |
| ATP-dependent | Dhx9 | NC | NS | NC | NS | NC | NS | 1.16 | 0.03 | NC | NS | NC | NS |

| | | | | | | | | | | | | | |
|---|---|---|---|---|---|---|---|---|---|---|---|---|---|
| RNA helicase A | | | | | | | | | | | | | |
| Dihydrolipoyl dehydrogenase, mitochondrial | Dld | NC | NS | NC | NS | NC | NS | NC | NS | NC | NS | 0.78 | 0.09 |
| Elongation factor 1-alpha 1 | Eef1a1 | NC | NS | NC | NS | NC | NS | NC | NS | 2.17 | 0.02 | NC | NS |
| Elongation factor 2 | Eef2 | NC | NS | NC | NS | NC | NS | 1.60 | 0.09 | NC | NS | 0.85 | 0.08 |
| ELAV-like protein 1 | Elavl1 | NC | NS | NC | NS | NC | NS | NC | NS | NC | NS | 0.96 | 0.06 |
| Alpha-enolase | Eno1 | NC | NS | -1.42 | 0.09 | NC | NS | 0.91 | 0.07 | NS | NC | NS | NC |
| Bifunctional glutamate/proline--tRNA ligase | Eprs | NC | NS | NC | NS | NC | NS | NC | NS | NC | NS | 0.97 | 0.09 |
| Neutral alpha-glucosidase AB | Ganab | NC | NS | 1.01 | 0.04 | NC | NS | 1.65 | 0.06 | NC | NS | NC | NS |
| Glyceraldehyde-3-phosphate dehydrogenase | Gapdh | NC | NS | 1.34 | 0.09 | NC | NS | NC | NS | NC | NS | NC | NS |
| Rab GDP dissociation inhibitor beta | Gdi2 | NC | NS | NC | NS | NC | NS | 1.25 | 0.05 | NC | NS | 1.62 | 0.01 |
| Golgi apparatus protein 1 | Glg1 | NC | NS | NC | NS | NC | NS | 1.05 | 0.04 | NC | NS | NC | NS |
| Glutamate dehydrogenase 1 | Glud1 | NC | NS | 1.28 | 0.01 | NC | NS | NC | NS | NC | NS | 0.95 | 0.06 |
| Heterogeneous nuclear ribonucleoprotein H2 | Hnrnph2 | NC | NS | NC | NS | NC | NS | 0.94 | 0.10 | NC | NS | NC | NS |
| Heterogeneous nuclear ribonucleoprotein K | Hnrnpk | NC | NS | NC | NS | NC | NS | 1.72 | 0.03 | NC | NS | NC | NS |
| Heterogeneous nuclear ribonucleoprotein U | Hnrnpu | NC | NS | NC | NS | NC | NS | 0.78 | 0.08 | NC | NS | NC | NS |

| | | | | | | | | | | | | | |
|---|---|---|---|---|---|---|---|---|---|---|---|---|---|
| Endoplasmin | Hsp90b1 | NC | NS | NC | NS | NC | NS | 1.29 | 0.04 | NC | NS | NC | NS |
| Stress-70 protein, mitochondrial | Hspa9 | NC | NS | 0.86 | 0.06 | NC | NS | NC | NS | NC | NS | 2.08 | 0.02 |
| 60 kDa heat shock protein, mitochondrial | Hspd1 | NC | NS | NC | NS | NC | NS | 3.03 | 0.01 | NC | NS | 1.38 | 0.02 |
| Basement membrane-specific heparan sulfate proteoglycan core protein | Hspg1 | NC | NS | NC | NS | NC | NS | 1.02 | 0.03 | -0.91 | 0.06 | NC | NS |
| Hypoxia up-regulated protein 1 | Hyou1 | NC | NS | NC | NS | NC | NS | 0.87 | 0.08 | NC | NS | NC | NS |
| LIM domain and actin-binding protein 1 | Lima1 | NC | NS | Inf | NA | NC | NS | Inf | NA | NC | NS | NC | NS |
| Prolow-density lipoprotein receptor-related protein 1 | Lrp1 | NC | NS | 1.14 | 0.03 | NC | NS | 2.03 | 0.01 | NC | NS | NC | NS |
| Matrin-3 | Matr3 | NC | NS | NC | NS | NC | NS | 1.14 | 0.07 | NC | NS | NC | NS |
| Malate dehydrogenase | Mdh2 | 1.48 | 0.05 | NC | NS | NC | NS | NC | NS | NC | NS | NC | NS |
| Lactadherin | Mfge8 | NC | NS | NC | NS | NC | NS | 0.88 | 0.09 | NC | NS | NC | NS |
| Myosin-10 | Myh10 | NC | NS | NC | NS | NC | NS | 1.05 | 0.06 | NC | NS | NC | NS |
| Myosin-9 | Myh9 | NC | NS | 0.88 | 0.07 | NC | NS | 1.03 | 0.05 | NC | NS | Inf | NA |
| Myosin regulatory light chain 12B | Myl12b | NC | NS | NC | NS | NC | NS | 2.18 | 0.07 | NC | NS | NC | NS |
| Myoferlin | Myof | NC | NS | 1.12 | 0.01 | NC | NS | NC | NS | NC | NS | 1.08 | 0.03 |
| Protein Niban 1 | Niban1 | NC | NS | NC | NS | NC | NS | 1.00 | 0.09 | NC | NS | NC | NS |
| Nucleoside diphosphate kinase B | Nme2 | -0.95 | 0.08 | NC | NS | NC | NS | NC | NS | NC | NS | NC | NS |

| | | | | | | | | | | | | | |
|---|---|---|---|---|---|---|---|---|---|---|---|---|---|
| Non-POU domain-containing octamer-binding protein | Nono | NC | NS | NC | NS | NC | NS | 0.87 | 0.09 | NC | NS | NC | NS |
| Nucleophosmin | Npm1 | NC | NS | NC | NS | NC | NS | 1.08 | 0.06 | NC | NS | NC | NS |
| Nucleobindin-1 | Nucb1 | | | | | | | | | -1.33 | 0.01 | -1.30 | 0.01 |
| Procollagen C-endopeptidase enhancer 1 | Pcolce | NC | NS | NC | NS | NC | NS | NC | NS | 1.47 | 0.06 | NC | NS |
| Prenylcysteine oxidase | Pcyox1 | NC | NS | NC | NS | NC | NS | 1.60 | 0.10 | NC | NS | NC | NS |
| Platelet-derived growth factor receptor beta | Pdgfrb | NC | NS | 2.13 | 0.00 | NC | NS | 1.71 | 0.01 | NC | NS | NC | NS |
| Profilin-1 | Pfn1 | NC | NS | 0.90 | 0.08 | NC | NS | 1.18 | 0.02 | NC | NS | 1.14 | 0.02 |
| Pyruvate kinase PKM | Pkm | NC | NS | NC | NS | 1.14 | 0.07 | NC | NS | NC | NS | NC | NS |
| Procollagen-lysine,2-oxoglutarate 5-dioxygenase 1 | Plod1 | NC | NS | NC | NS | NC | NS | 0.77 | 0.07 | NC | NS | NC | NS |
| Peripherin | Prph | NC | NS | NC | NS | NC | NS | 1.21 | 0.05 | NC | NS | NC | NS |
| Prostacyclin synthase | Ptgis | NC | NS | 1.22 | 0.02 | NC | NS | 2.01 | 0.00 | NC | NS | 1.08 | 0.04 |
| Dolichyl-diphosphooligosaccharide--protein glycosyltransferase subunit 1 | Rpn1 | NC | NS | NC | NS | NC | NS | 1.14 | 0.06 | NC | NS | 1.20 | 0.06 |
| RNA-splicing ligase RtcB homolog | Rtcb | NC | NS | NC | NS | NC | NS | 1.09 | 0.10 | NC | NS | 1.26 | 0.06 |
| Vesicle-trafficking protein SEC22b | Sec22b | NC | NS | NC | NS | NC | NS | NC | NS | NC | NS | 1.07 | 0.02 |
| Protein transport | Sec31a | NC | NS | NC | NS | NC | NS | NC | NS | NC | NS | 1.12 | 0.04 |

| | | | | | | | | | | | | | |
|---|---|---|---|---|---|---|---|---|---|---|---|---|---|
| protein Sec31A | | | | | | | | | | | | | |
| Serpin H1 | Serpinh1 | NC | NS | NC | NS | NC | NS | 2.50 | 0.04 | NC | NS | NC | NS |
| ADP/ATP translocase 1 | Slc25a4 | NC | NS | 1.01 | 0.04 | NC | NS | 0.86 | 0.07 | NC | NS | NC | NS |
| Staphylococcal nuclease domain-containing protein 1 | Snd1 | NC | NS | NC | NS | NC | NS | 0.84 | 0.08 | NC | NS | NC | NS |
| Sequestosome-1 | Sqstm1 | NC | NS | -1.28 | 0.08 | NC | NS | NC | NS | 2.19 | 0.01 | NC | NS |
| Transgelin | Tagln | -1.55 | 0.03 | NC | NS | NC | NS | 0.70 | 0.09 | NC | NS | 0.83 | 0.05 |
| Transgelin-2 | Tagln2 | NC | NS | NC | NS | NC | NS | NC | NS | NC | NS | -0.85 | 0.09 |
| T-complex protein 1 subunit alpha | Tcp1 | NC | NS | 1.51 | 0.01 | NC | NS | 1.55 | 0.01 | NC | NS | 1.68 | 0.00 |
| Thrombospondin-1 | Thbs1 | NC | NS | NC | NS | NC | NS | 0.94 | 0.08 | NC | NS | NC | NS |
| Transmembrane emp24 domain-containing protein 10 | Tmed10 | NC | NS | NC | NS | NC | NS | 1.05 | 0.06 | NC | NS | NC | NS |
| Tubulin alpha-3 chain | Tuba3a | NC | NS | NC | NS | NC | NS | 1.12 | 0.09 | NC | NS | NC | NS |
| Tubulin beta-3 chain | Tubb3 | NC | NS | NC | NS | -1.28 | 0.07 | NC | NS | NC | NS | NC | NS |
| Thioredoxin domain-containing protein 12 | Txndc12 | NC | NS | NC | NS | NC | NS | 0.75 | 0.09 | NC | NS | NC | NS |
| Transitional endoplasmic reticulum ATPase | Vcp | NC | NS | NC | NS | NC | NS | 1.14 | 0.04 | NC | NS | 1.05 | 0.06 |
| Vimentin | Vim | NC | NS | NC | NS | NC | NS | 1.36 | 0.03 | NC | NS | NC | NS |

NA = no appropriate NC = no change NS = not significant Inf = only identified in CDC treated samples

**Supplemental table 5:** Listed are mitochondrial proteins changed following CDC treatment, that are able to regulate mitochondrial protein expression. Shown are relevant gene ontology functions and the foldchanges (log2) and p-values for HFpEF compared to control rats and CDC treatment of HFpEF rats compared to placebo treatment (n=6).

| Protein names | Gene name | Function | Foldchange (log2) HFpEF vs control | p-value HFpEF vs control | Foldchange (log2) CDC vs placebo | p-value CDC vs placebo |
|---|---|---|---|---|---|---|
| DNA replication ATP-dependent helicase/nuclease DNA2 | Dna2 | mitochondrial DNA replication and repair [11, 12] | -2.75 | 0.000 | 0.97 | 0.000 |
| Endonuclease G, mitochondrial | Endog | mitochondrial apoptotic DNA fragmentation [13] | -0.25 | 0.030 | -0.25 | 0.021 |
| FAST kinase domain-containing protein 2, mitochondrial | Fastkd2 | mitochondrial large ribosomal subunit assembly and controlling 16S mt-rRNA abundance [14, 15] | 4.85 | 0.000 | -5.68 | 0.000 |
| Glutamyl-tRNA(Gln) amidotransferase subunit C, mitochondrial | Gatc | mitochondrial translation [16], glutaminyl-tRNAGln biosynthesis via transamidation [16] | -1.20 | 0.023 | 1.02 | 0.036 |
| Elongation factor G, mitochondrial | Gfm1 | mitochondrial translational elongation [17] | -0.34 | 0.000 | 0.26 | 0.002 |
| Ribosome-releasing factor 2, mitochondrial | Gfm2 | mitochondrial translational termination [17] | -0.81 | 0.010 | 0.80 | 0.007 |
| Lon protease homolog, mitochondrial | Lonp1 | Regulation of mt gene expression and maintenance of mt genome integrity. Binds to mitochondrial promoters and RNA in a single-stranded, site-specific, and strand-specific manner. Regulation of mt DNA replication and/or gene expression using site-specific, single-stranded DNA binding to target the degradation of regulatory proteins binding to adjacent sites in mt promoters [18-21] | NC | NS | 0.28 | 0.003 |
| Phosphatidate phosphatase LPIN1 | Lpin1 | Involved in mitochondrial fission by converting phosphatidic acid to diacylglycerol [22] | 0.54 | 0.000 | -0.42 | 0.000 |
| Leucine-rich PPR motif-containing protein, mitochondrial | Lrpprc | Involved in mt RNA metabolism, plays a role in translation or stability of mitochondrially encoded cytochrome c oxidase (COX) subunits, cooperates with PPARGC1A to regulate certain mitochondrially encoded genes [23] | -0.32 | 0.000 | 0.33 | 0.000 |
| 39S ribosomal protein L12, mitochondrial | Mrpl12 | Associates with mitochondrial RNA polymerase to activate transcription [24] | -0.39 | 0.001 | 0.20 | 0.034 |
| 39S ribosomal protein L28, mitochondrial | Mrpl28 | Structural constituent of ribosome [13] | NC | NS | 3.59 | 0.001 |
| 39S ribosomal protein L40, mitochondrial | Mrpl40 | mitochondrial Translation elongation (Source: Reactome) | NC | NS | 0.39 | 0.001 |

| | | | | | | |
|---|---|---|---|---|---|---|
| 39S ribosomal protein L49, mitochondrial | Mrpl49 | Component of the mitochondrial large ribosomal subunit (mt-LSU) [25-27] | -0.55 | 0.000 | 0.24 | 0.003 |
| 28S ribosomal protein S14, mitochondrial | Mrps14 | mitochondrial translation [28] | -2.08 | 0.000 | 1.08 | 0.005 |
| 28S ribosomal protein S15, mitochondrial | Mrps15 | mitochondrial translation [29] | NC | NS | 1.07 | 0.047 |
| 28S ribosomal protein S23, mitochondrial | Mrps23 | structural constituent of ribosome [30] | -1.35 | 0.003 | 0.95 | 0.002 |
| 28S ribosomal protein S27, mitochondrial | Mrps27 | RNA-binding component of the mitochondrial small ribosomal subunit (mt-SSU) that plays a role in mitochondrial protein synthesis [31]<br>Stimulates mitochondrial mRNA translation of subunit components of the mitochondrial electron transport chain [31] | -3.26 | 0.001 | 3.13 | 0.001 |
| 28S ribosomal protein S5, mitochondrial | Mrps5 | structural constituent of ribosome [13] | -0.64 | 0.000 | 0.42 | 0.005 |
| Methionyl-tRNA formyltransferase, mitochondrial | Mtfmt | Formylates methionyl-tRNA in mitochondria and the cytoplasm [32, 33]<br>Responsible for the formylation of the 8 N-terminally formylated (Nt-formylated) mitochondrial matrix proteins that are encoded by mitochondrial DNA [32, 34] | 0.81 | 0.000 | 0.52 | 0.003 |
| Pentatricopeptide repeat domain-containing protein 3, mitochondrial | Ptcd3 | Mitochondrial RNA-binding protein that has a role in mitochondrial translation [35] | -0.66 | 0.002 | 0.62 | 0.003 |
| Serine--tRNA ligase, mitochondrial | Sars2 | Catalyzes the attachment of serine to tRNA(Ser). Is also probably able to aminoacylate tRNA(Sec) with serine, to form the misacylated tRNA L-seryl-tRNA(Sec), which will be further converted into selenocysteinyl-tRNA(Sec) [36] | -0.98 | 0.001 | 0.93 | 0.001 |
| Translational activator of cytochrome c oxidase 1 | Taco1 | Acts as a translational activator of mitochondrially-encoded cytochrome c oxidase 1 [37] | NC | NS | 0.43 | 0.005 |
| Elongation factor Ts, mitochondrial | Tsfm | Regulation of mitochondrial translation [38] | NC | NS | 0.43 | 0.001 |
| Thiosulfate sulfurtransferase | Tst | Together with MRPL18, acts as a mitochondrial import factor for the cytosolic 5S rRNA [39] | -0.76 | 0.000 | 0.34 | 0.003 |
| Ubiquinol-cytochrome-c reductase complex assembly factor 2 | Uqcc2 | Required for the assembly mitochondrial respiratory chain complex III. Involved in cytochrome b translation and/or stability [40, 41] | -1.22 | 0.005 | 1.57 | 0.001 |

NC = not changed NS= not significant

**Supplemental table 6**: Listed are the dominant kinases and their regulators sharing consensus sequences with changed phosphorylated residues of proteins regulating protein expression following CDC treatment.

| Protein Name | Gene Name | Regulates Upstream Kinase | Function | Phospho-site | CDC vs Placebo (log2FC) | p-value |
|---|---|---|---|---|---|---|
| Cysteine and glycine-rich protein 3 | Csrp3 | PKC | In vitro can inhibit PKC/PRKCA activity. Proposed to be involved in cardiac stress signaling by down-regulating excessive PKC/PRKCA signaling [42] | | -0.63 | <0.001 |
| PDZ and LIM domain protein 5 | Pdlim5 | PKC | Scaffolding PKC to the Z-disk region [43-45] | S228 | -2.63 | 0.059 |
| PDZ and LIM domain protein 5 | Pdlim5 | PKC | Scaffolding PKC to the Z-disk region [43-45] | | -0.43 | 0.042 |
| Protein kinase C delta type | Prkcd | PKC | Scaffolding PKC to the Z-disk region [43-45] | | -1.81 | <0.001 |
| Receptor of activated protein C kinase 1 | Rack1 | PKC | Binds to and stabilizes activated protein kinase C (PKC), increasing PKC-mediated phosphorylation [46] | | -0.29 | 0.012 |
| 14-3-3 protein sigma | Sfn | PKC / MTOR | Protein kinase C inhibitor activity [47] Regulates with KRT17 protein synthesis and epithelial cell growth by stimulating Akt/mTOR pathway [48] | | 3.38 | 0.002 |
| Protein SEC13 homolog | Sec13 | MTOR | Indirectly activates mTORC1 and the TORC1 signaling pathway through the inhibition of the GATOR1 subcomplex [49] | | 0.55 | 0.059 |
| N-alpha-acetyltransferase 10 | Naa10 | MTOR | Acetylates, and stabilizes TSC2, thereby repressing mTOR activity and suppressing cancer development [50] | | 0.96 | 0.04 |
| Keratin, type I cytoskeletal 17 | KRT17 | Akt/MTOR | Stimulating Akt/mTOR pathway [51] | | -2.56 | <0.001 |
| Glycogen synthase kinase-3 alpha | GSK3A | | | | 1.15 | <0.001 |
| Protein MEMO1 | Memo1 | GSK3B | Regulates GSK3B activity [52, 53] | | -0.33 | 0.039 |
| Integrin-linked protein kinase | Ilk | Akt/GSK3B | Phosphorylates Akt1 and GSK3B [54] | | -0.27 | 0.02 |
| RAC-alpha serine/threonine-protein kinase | Akt | | | | -15 ± 9.8% (WB*) | <0.05 |
| RAC-alpha serine/threonine-protein kinase | Akt | | | S126, S129 | -2.09 | 0.02 |
| A-kinase anchor protein 12 | Akap12 | PKC | Anchoring protein that mediates the subcellular compartmentation of protein kinase A (PKA) and protein kinase C (PKC) [55] | | 0.86 | 0.005 |

| | | | | | | |
|---|---|---|---|---|---|---|
| cAMP-dependent protein kinase inhibitor alpha | Pkia | PKA | Extremely potent competitive inhibitor of cAMP-dependent protein kinase activity [56] | | -1.13 | 0.04 |
| cAMP-dependent protein kinase type I-beta regulatory subunit | Prkab1 | | | | 0.38 | 0.01 |
| cAMP-dependent protein kinase catalytic subunit beta | Prkacb | | | | -0.34 | <0.001 |
| cAMP-dependent protein kinase type II-beta regulatory subunit | Prkar2b | | | | 1.09 | 0.01 |

WB* = foldchange was estimated with Western blot analysis.

**Supplemental table 7:** Listed are proteins identified in CDC exosomes relevant for regulating the identified key regulators of CDC therapy. Shown are the function related to the identified upstream regulator, the protein probability showing the significance for the identified protein and amount of identified unique peptides (n=3). Only proteins with an FDR lower than 5% (protein probability >0.95) were included). Nonunique, overlapping sequences were excluded from the analysis as well.

| Protein Name | Gen Name | Protein ID | Regulator function | Protein Probability | Distinct Peptides |
|---|---|---|---|---|---|
| Complement component 1 Q subcomponent-binding protein | C1QBP | Q07021 | Signaling involved in inhibition of innate immune response is implicating the PI3K-AKT/PKB pathway [57]. | 1 | 2 |
| Complement C3 | C3 | P01024 | Appears to stimulate TG synthesis via activation of the PLC, MAPK and AKT signaling pathways [58, 59]. | 1 | 80 |
| Caveolin-1 | CAV1 | Q03135 | Negatively regulates TGFB1-mediated activation of SMAD2/3 by mediating the internalization of TGFBR1 from membrane rafts leading to its subsequent degradation [60]. | 1 | 1 |
| CD109 antigen | CD109 | Q6YHK3 | Modulates negatively TGFB1 signaling in keratinocytes [61]. | 1 | 4 |
| CD63 antigen | CD63 | P08962 | Plays a role in the activation of ITGB1 and integrin signaling, leading to the activation of AKT, FAK/PTK2 and MAP kinases. Promotes cell survival, reorganization of the actin cytoskeleton, cell adhesion, spreading and migration, via its role in the activation of AKT and FAK/PTK2 [62]. | 1 | 1 |
| Chitinase-3-like protein 1 | CHI3L1 | P36222 | Mediates activation of AKT1 signaling pathway and subsequent IL8 production in colonic epithelial cells [63, 64]. | 1 | 3 |
| Dipeptidyl peptidase 4 | DPP4 | P27487 | binding to CAV1 and CARD11 induces T-cell proliferation and NF-kappa-B activation in a T-cell receptor/CD3-dependent manner [65, 66]. | 1 | 8 |
| Eukaryotic translation initiation factor 5A-1 | EIF5A | P63241 | With syntenin SDCBP, functions as a regulator of p53/TP53 and p53/TP53-dependent apoptosis [67]. | 1 | 1 |
| Endoglin | ENG | P17813 | Acts as TGF-beta coreceptor and is involved in the TGF-beta/BMP signaling cascade leading ultimately to the activation of SMAD transcription factors [68-71]. | 1 | 1 |
| Heat shock 70 kDa protein 1A | HSPA1A | P0DMV8 | Maintains protein homeostasis during cellular stress binding to HOPX which assists in chaperone-mediated protein refolding [72]. | 0.999 | 4 |
| Heat shock cognate 71 kDa protein | HSPA8 | P11142 | HOPX (HSPA8) is HSP70-associated co-chaperone [73, 74]. | 1 | 14 |
| Integrin alpha-V | ITGAV | P06756 | Integrin alpha-V mediates release of TGF-beta-1 from regulatory Latency-associated peptide (LAP) and thus mediates TGF-beta-1 activation [75, 76]. HMGB1 inhibits macrophage activity in efferocytosis through binding to the alphavbeta3-integrin [77]. | 1 | 6 |

| | | | | | |
|---|---|---|---|---|---|
| Keratin, type I cytoskeletal 17 | KRT17 | Q04695 | Regulates protein synthesis and epithelial cell growth through binding to the adapter protein SFN and by stimulating Akt/mTOR pathway [51]. | 1 | 3 |
| Latent-transforming growth factor beta-binding protein 1 | LTBP1 | Q14766 | Critical role in controlling and directing the activity of TGFB1 [78-80]. | 1 | 3 |
| Tyrosine-protein kinase Lyn | LYN | P07948 | Regulates phosphatidylinositol 3-kinase activity and AKT1 activation [81-100]. | 0.994 | 1 |
| Nucleoside diphosphate kinase B | NME2 | P22392 | Acts as a transcriptional activator of the MYC gene [101]. | 1 | 2 |
| Nucleophosmin | NPM1 | P06748 | Regulation of tumor suppressors p53/TP53. In complex with MYC enhances the transcription of MYC target genes [102-110] | 1 | 2 |
| Profilin-1 | PFN1 | P07737 | Inhibits androgen receptor (AR) and HTT aggregation and binding of G-actin is essential for its inhibition of AR [111]. | 1 | 4 |
| Retinoic acid receptor responder protein 2 | RARRES2 | Q99969 | The effect is mediated via inhibiting activation of NF-kappa-B and CRK/p38 through stimulation of AKT1/NOS3 signaling and nitric oxide production [112-118]. | 1 | 1 |
| 60S ribosomal protein L11 | RPL11 | P62913 | It also couples ribosome biogenesis to p53/TP53 activation. As part of the 5S RNP it accumulates in the nucleoplasm and inhibits MDM2, when ribosome biogenesis is perturbed, mediating the stabilization and the activation of TP53 [119]. | 0.982 | 1 |
| 40S ribosomal protein S3 | RPS3 | P23396 | Binds to and protects TP53/p53 from MDM2-mediated ubiquitination [120]. | 1 | 2 |
| Syntenin-1 | SDCBP | O00560 | Positively regulates TGFB1-mediated SMAD2/3 activation and TGFB1-induced epithelial-to-mesenchymal transition (EMT) and cell migration in various cell types. May increase TGFB1 signaling by enhancing cell-surface expression of TGFR1 by preventing the interaction between TGFR1 and CAV1 and subsequent CAV1-dependent internalization and degradation of TGFR1 [60]. | 1 | 4 |
| 14-3-3 protein sigma | SFN | P31947 | When bound to KRT17, regulates protein synthesis and epithelial cell growth by stimulating Akt/mTOR pathway. May also regulate MDM2 autoubiquitination and degradation and thereby activate p53/TP53 [48].; FUNCTION: p53-regulated inhibitor of G2/M progression [48]. | 0.981 | 3 |
| 4F2 cell-surface antigen heavy chain | SLC3A2 | P08195 | When associated with LAPTM4B, recruits SLC3A2 and SLC7A5 to lysosomes to promote leucine uptake into these organelles and is required for mTORC1 activation [121]. | 1 | 5 |
| Transitional endoplasmic reticulum ATPase | VCP | P55072 | Acts as a negative regulator of type I interferon production by interacting with DDX58/RIG-I: interaction takes place when DDX58/RIG-I is ubiquitinated via 'Lys-63'-linked ubiquitin on its CARD domains}leading to recruit RNF125 and promote ubiquitination and degradation of DDX58/RIG-I [122]. Is involved in DNA damage response: recruited to double-strand breaks | 1 | 3 |

| | | | | | |
|---|---|---|---|---|---|
| | | | (DSBs) sites in a RNF8- and RNF168-dependent manner and promotes the recruitment of TP53BP1 at DNA damage sites [123, 124]. | | |
| Tryptophan--tRNA ligase | WARS | P23381 | Regulates ERK, Akt, and eNOS activation pathways that are associated with angiogenesis, cytoskeletal reorganization, and shear stress-responsive gene expression [125-128]. | 0.976 | 1 |
| Nuclease-sensitive element-binding protein 1 | YBX1 | P22392 | Component of the CRD-mediated complex that promotes MYC mRNA stability [129]. | 0.998 | 1 |

**Supplemental table 8:** Demographic, structural, and hemodynamic data of human HFpEF patients and their referent control.

| | **Referent Control** | **HFpEF** | **p-value** |
|---|---|---|---|
| **Age (years)** | 70 ± 10 | 72 ± 9 | 0.752 |
| **BSA ($m^2$)** | 2.0 ± 0.3 | 2.2 ± 0.2 | 0.233 |
| **SBP (mmHg)** | 125 ± 11 | 135 ± 11 | 0.171 |
| **DBP (mmHg)** | 69 ± 10 | 77 ± 11 | 0.271 |
| **HR (bpm)** | 71 ± 4 | 69 ± 12 | 0.666 |
| **LV EDV (mL)** | 104 ± 38 | 107 ± 7 | 0.838 |
| **LV $EDV_i$ ($mL/m^2$)** | 46 ±12 | 51 ± 19 | 0.599 |
| **LV ESV (mL)** | 32 ±14 | 51 ± 21 | 0.215 |
| **LV $ESV_i$ ($mL/m^2$)** | 16 ± 6 | 24 ± 10 | 0.199 |
| **LV SV (mL)** | 72 ± 26 | 57 ± 16 | 0.319 |
| **LV EF (%)** | 67 ± 8 | 64 ± 6 | 0.529 |
| **LV Mass (g)** | 152 ± 12 | 254 ± 47 | ***0.006*** |
| **LV $Mass_i$ ($g/m^2$)** | 77 ± 5 | 117 ± 23 | ***0.016*** |
| **E (cm/sec)** | 69 ±14 | 95 ± 28 | 0.104 |
| **e' (cm/sec)** | 10 ± 1 | 7.5 ± 3.4 | 0.163 |
| **E/e'** | 6.8 ± 1.2 | 13.2 ± 2.8 | ***0.004*** |
| **PCWP (mmHg)** | 10.5 ± 1.7 | 19.2 ± 4.3 | ***0.003*** |
| **LV EDP (mmHg)** | 9.0 ± 1.7 | 20.0 ± 7.1 | ***0.010*** |
| **LA dimension (cm)** | 3.7 ± 0.5 | 4.7 ± 0.6 | ***0.016*** |
| **LA Vol (mL)** | 44 ± 6 | 70 ± 23 | ***0.010*** |
| **LA $Vol_i$ ($mL/m^2$)** | 22 ± 3 | 37 ± 9 | ***0.040*** |
| **CVF (%)** | 0.68 ± 0.16 | 1.46 ± 0.17 | ***0.001*** |

**Abbreviations:** Data=Mean±St.Dev; BSA=body surface area; SBP=systolic blood pressure; DBP=diastolic blood pressure; HR=heart rate; LV=left ventricular; EDV=end diastolic volume; EDVi=end diastolic volume index; ESV=end systolic volume; ESVi=end systolic volume index; SV=stroke volume; EF=ejection fraction;

E=transmitral Doppler early filling velocity; e'=tissue Doppler early diastolic mitral annular velocity; PCWP=Pulmonary capillary wedge pressure; EDP=end diastolic pressure; LA=left atrium; CVF=collagen volume fraction.

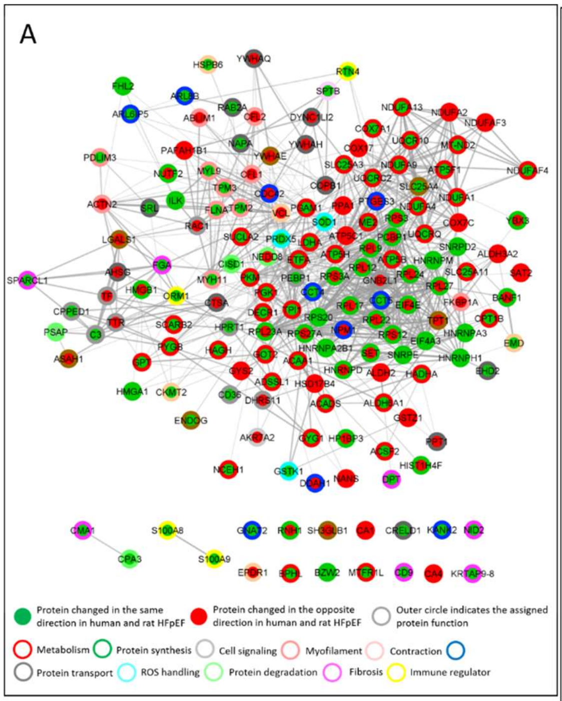

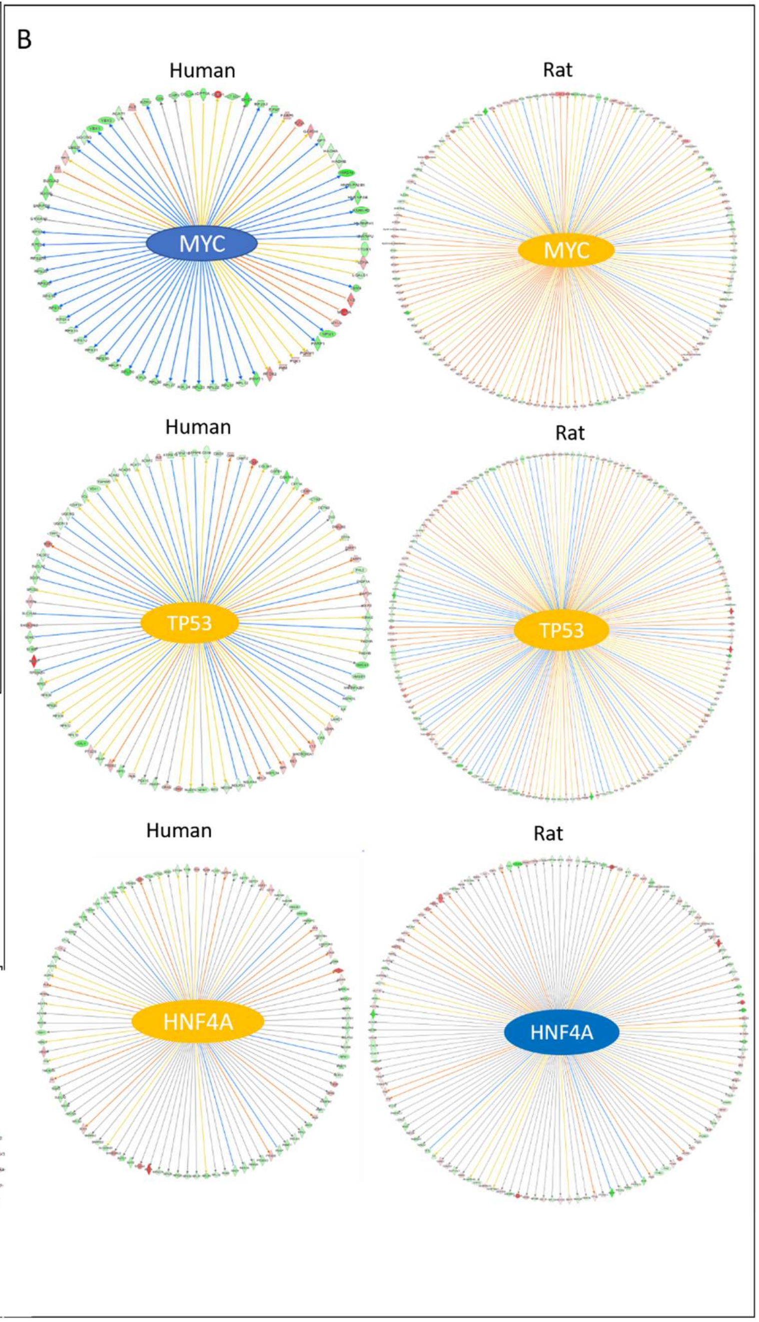

C

| Top upstream regulators predicted for overlap DEPs for HFpEF vs control hearts comparison | |
|---|---|
| **Human** | **Rat** |
| Rictor | MYC |
| MAPT | TP53 |
| LARP1 | MLXIPL |
| HNF4A | MYCN |
| TLE3 | YAP1 |
| MYC | CTNNB1 |
| TP53 | PPARA |
| CD437 | HNF4A |

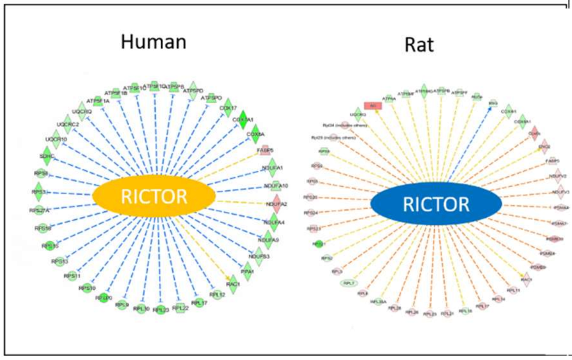

**Supplemental figure 1:** In panel A is a network of the overlapping DEPs from rat and human HFpEF hearts compared to control hearts shown. The color of the inner circle show change in same direction (green) or opposite direction (red) while the color of the outer circle indicates protein function (A). Panel B shows proteins downstream of the major relevant upstream regulator MYC, TP53, HNF4A, and RICTOR in human verses rat HFpEF. The color red indicates an increase and the color green a decrease in target protein concentration. The color orange an activation and the color blue an inhibition of key regulatory proteins.

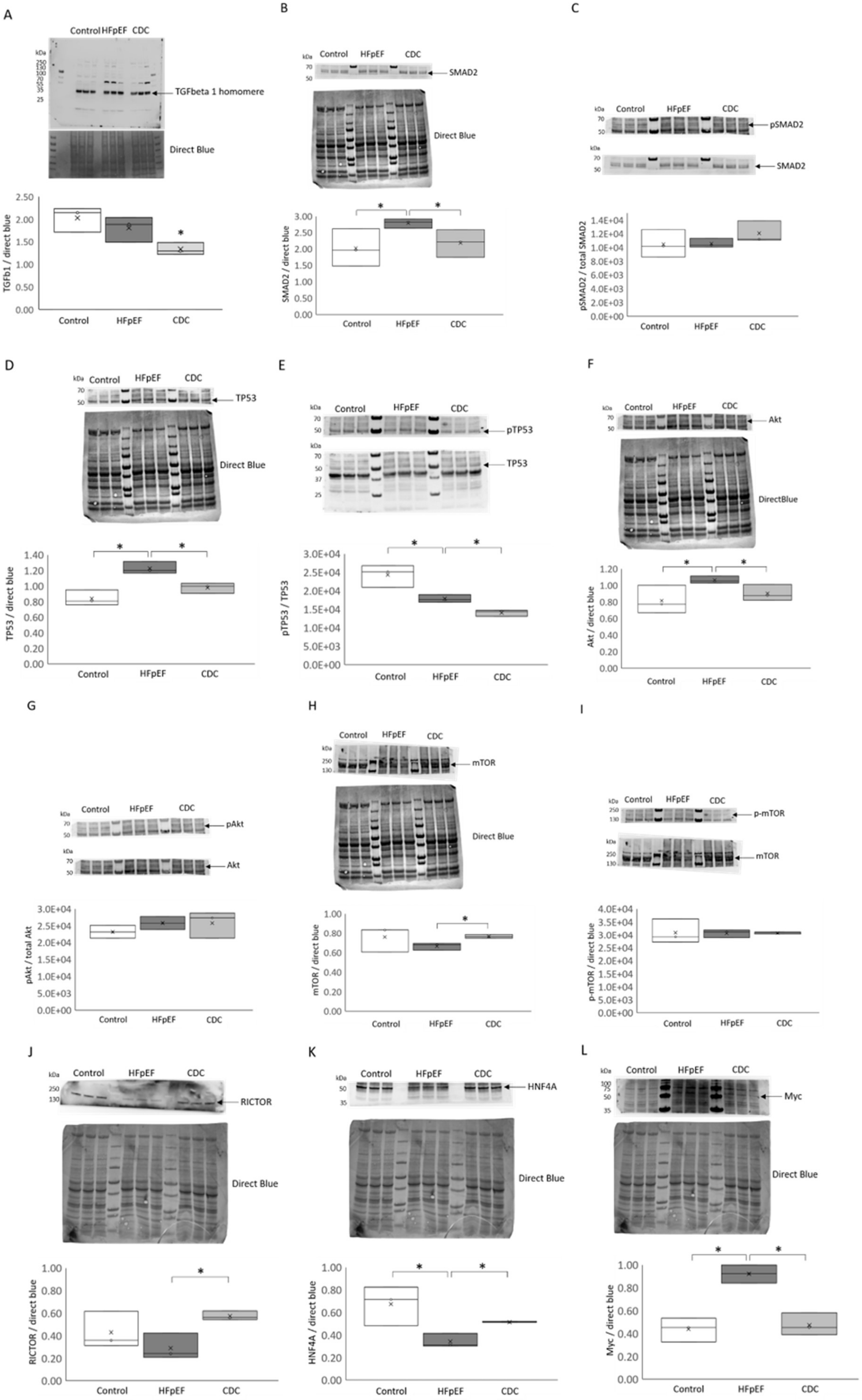

A
Control HFpEF CDC
kDa
250
130
100
70
55
35
25
TGFbeta 1 homomere
Direct Blue
TGFb1 / direct blue
2.50
2.00
1.50
1.00
0.50
0.00
*
Control
HFpEF
CDC
B
kDa
Control
HFpEF
CDC
70
50
SMAD2
Direct Blue
SMAD2 / direct blue
3.00
2.50
2.00
1.50
1.00
0.50
0.00
*
*
Control
HFpEF
CDC
C
kDa
Control
HFpEF
CDC
70
50
pSMAD2
SMAD2
pSMAD2 / total SMAD2
1.4E+04
1.2E+04
1.0E+04
8.0E+03
6.0E+03
4.0E+03
2.0E+03
0.0E+00
Control
HFpEF
CDC
D
kDa
Control
HFpEF
CDC
70
50
TP53
Direct Blue
TP53 / direct blue
1.40
1.20
1.00
0.80
0.60
0.40
0.20
0.00
*
*
Control
HFpEF
CDC
E
kDa
Control
HFpEF
CDC
70
50
pTP53
kDa
70
50
37
25
TP53
pTP53 / TP53
3.0E+04
2.5E+04
2.0E+04
1.5E+04
1.0E+04
5.0E+03
0.0E+00
*
*
Control
HFpEF
CDC
F
kDa
Control
HFpEF
CDC
70
50
Akt
DirectBlue
Akt / direct blue
1.20
1.00
0.80
0.60
0.40
0.20
0.00
*
*
Control
HFpEF
CDC
G
kDa
Control
HFpEF
CDC
70
50
pAkt
kDa
70
50
Akt
pAkt / total Akt
3.0E+04
2.5E+04
2.0E+04
1.5E+04
1.0E+04
5.0E+03
0.0E+00
Control
HFpEF
CDC
H
Control
HFpEF
CDC
kDa
250
130
mTOR
Direct Blue
mTOR / direct blue
1.00
0.80
0.60
0.40
0.20
0.00
*
Control
HFpEF
CDC
I
kDa
Control
HFpEF
CDC
250
130
p-mTOR
250
130
mTOR
p-mTOR / direct blue
4.0E+04
3.5E+04
3.0E+04
2.5E+04
2.0E+04
1.5E+04
1.0E+04
5.0E+03
0.0E+00
Control
HFpEF
CDC
J
kDa
Control
HFpEF
CDC
250
130
RICTOR
Direct Blue
RICTOR / direct blue
1.00
0.80
0.60
0.40
0.20
0.00
*
Control
HFpEF
CDC
K
kDa
Control
HFpEF
CDC
50
35
HNF4A
Direct Blue
HNF4A / direct blue
1.00
0.80
0.60
0.40
0.20
0.00
*
*
Control
HFpEF
CDC
L
kDa
Control
HFpEF
CDC
100
75
50
35
Myc
Direct Blue
Myc / direct blue
1.00
0.80
0.60
0.40
0.20
0.00
*
*
Control
HFpEF
CDC

**Supplemental figure 2:** Immunoreactivity assays of rat LV quantifying identified key regulators of CDC therapy following CDC treatment of HFpEF. WB analysis were performed for TGFb1 (A), SMAD (B), phospho-SMAD (Ser465/467) (C), TP53 (D), phospho-TP53 (Ser15) (E), Akt (F), phospho-Akt (G), mTOR (H), phospho-mTOR (I), RICTOR (J), HNF4A (K), and Myc (L). Intensities were either normalized to total protein staining with direct blue or phospho-intensities were normalized to the equivalent total. * indicates significance $p \leq 0.05$ among treatment groups.

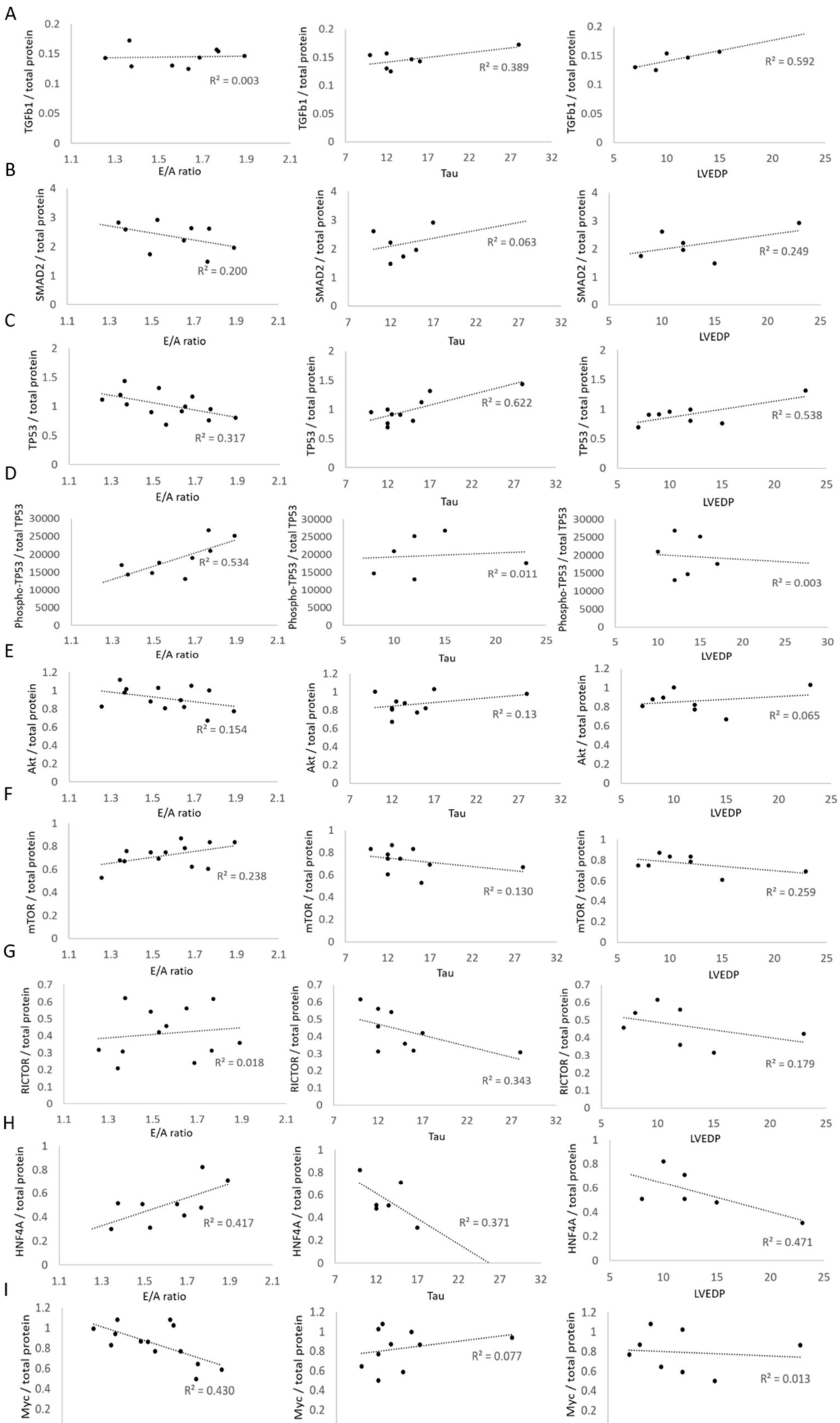

A
TGFb1 / total protein
E/A ratio
R² = 0.003
Tau
R² = 0.389
LVEDP
R² = 0.592
B
SMAD2 / total protein
E/A ratio
R² = 0.200
Tau
R² = 0.063
LVEDP
R² = 0.249
C
TP53 / total protein
E/A ratio
R² = 0.317
Tau
R² = 0.622
LVEDP
R² = 0.538
D
Phospho-TP53 / total TP53
E/A ratio
R² = 0.534
Tau
R² = 0.011
LVEDP
R² = 0.003
E
Akt / total protein
E/A ratio
R² = 0.154
Tau
R² = 0.13
LVEDP
R² = 0.065
F
mTOR / total protein
E/A ratio
R² = 0.238
Tau
R² = 0.130
LVEDP
R² = 0.259
G
RICTOR / total protein
E/A ratio
R² = 0.018
Tau
R² = 0.343
LVEDP
R² = 0.179
H
HNF4A / total protein
E/A ratio
R² = 0.417
Tau
R² = 0.371
LVEDP
R² = 0.471
I
Myc / total protein
E/A ratio
R² = 0.430
Tau
R² = 0.077
LVEDP
R² = 0.013

**Supplemental figure 3:** The Graphs are showing the correlation between functional data (E/A ratio, Tau, and LVEDP) and key regulatory protein concentrations of specific animals (HFpEF + placebo, HFpEF + CDC, and control) used in this study.

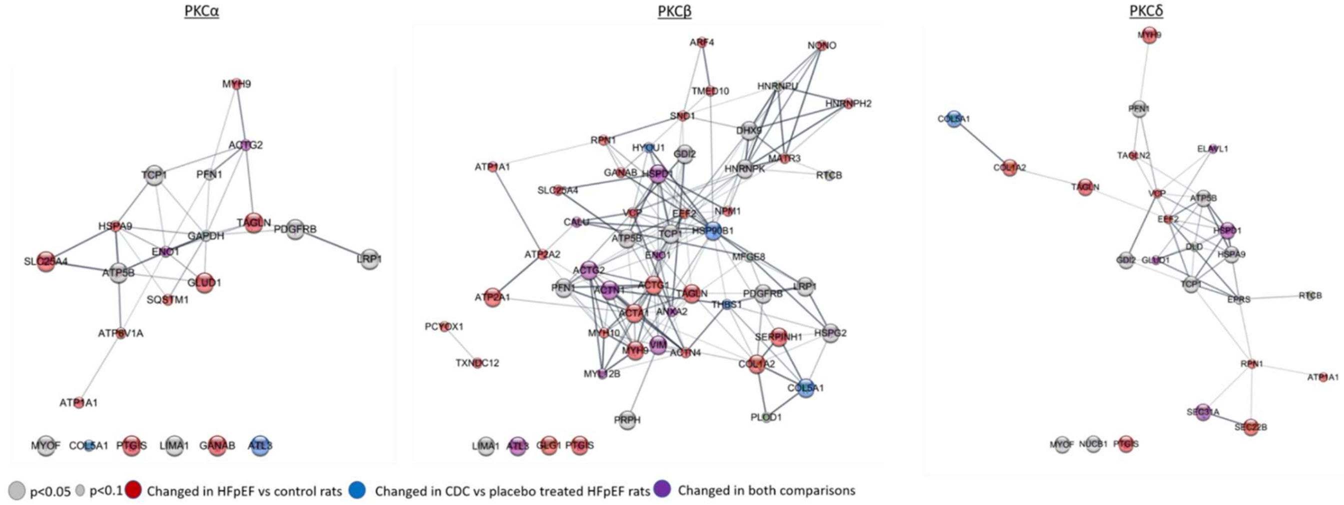

**Supplemental figure 4:** NSPs were quantified in H9C2 cells following PKC inhibition. Shown is an interaction network of significantly changed NSPs following PKC isoform inhibition overplayed with changed proteins from CDC treated HFpEF rats.

### Online Supplements – References
